# Nitric oxide feedback to ciliary photoreceptor cells gates a UV avoidance circuit

**DOI:** 10.1101/2023.08.02.551600

**Authors:** Kei Jokura, Piotr Słowiński, Nobuo Ueda, Martin Gühmann, Luis Alfonso Yañez-Guerra, Emily Savage, Kyle C. A. Wedgwood, Gáspár Jékely

**Affiliations:** Living Systems Institute, University of Exeter, Exeter EX4 4QD, United Kingdom; Biosciences, Faculty of Health and Life Sciences, University of Exeter, Exeter EX4 4QD, United Kingdom; Okinawa Institute of Science and Technology, Okinawa, Japan; School of Biological Sciences, University of Bristol, Bristol, United Kingdom; School of Biological Sciences, University of Southampton, Southampton, United Kingdom; Centre for Organismal Studies (COS), Heidelberg University, 69120 Heidelberg, Germany

## Abstract

Nitric oxide (NO) produced by nitric-oxide synthase (NOS) is a key regulator of animal physiology. Here we uncover a function for NO in the integration of UV exposure and the gating of a UV-avoidance circuit. We studied UV/violet avoidance mediated by brain ciliary photoreceptors (cPRCs) in larvae of the annelid *Platynereis dumerilii*. In the larva, NOS is expressed in interneurons (INNOS) postsynaptic to cPRCs. UV stimulation of cPRCs triggers INNOS activation and NO production. NO signals retrogradely to cPRCs to induce their sustained post-stimulus activation through an unconventional guanylate cyclase. This late activation drives ciliary beating through serotonergic ciliomotor neurons during UV avoidance. In *NOS* mutants, retrograde signalling, ciliary activation and UV avoidance are defective. By mathematical modelling, we recapitulate phototransduction and circuit dynamics in wild-type and mutant larvae. Our results reveal how NO-mediated retrograde signalling gates a synaptic circuit and induces short-term memory of UV exposure to orchestrate light-avoidance behaviour.

## Introduction

In nervous systems, synaptic transmission and volume transmission together shape circuit dynamics (Bargmann and Marder, 2013). While synaptic transmission occurs at specialised contact sites, volume transmission is characterised by the delocalised release of diverse diffusive neuromodulators.

Nitric oxide (NO) is one such modulator with unique physical and signalling properties. This free radical synthesized from L-arginine by nitric oxide synthase (NOS) is short-lived and can diffuse across biological membranes (Cudeiro and Rivadulla, 1999; Thomas, 2015). Canonical NO signalling involves the Ca^2+^/calmodulin-dependent activation of NOS, NO production and diffusion, and the NO-dependent activation of soluble guanylate cyclases (sGC) leading to cGMP production (Bredt et al., 1990; Hölscher, 1997). Given that NOS activation is Ca^2+^ dependent and NOS shows neuron-type-specific expression (Aso et al., 2019; Gibbs and Truman, 1998; Mobley et al., 2022; Wildemann and Bicker, 1999), NO action can lead to the activity-dependent modulation of neural circuits at specific sites (Aso et al., 2019; Jacoby et al., 2018; Vielma et al., 2014; Wang et al., 2007).

In the vertebrate retina, NOS is expressed in amacrine, ganglion and other cells and the actions of NO can be diverse (Cudeiro and Rivadulla, 1999; Jacoby et al., 2018; Wang et al., 2007). For example, defective NO signalling in NOS knockout mice leads to a decreased sensitivity of retinal ganglion cells to light stimulation (Wang et al., 2007). In the retina, NO signalling can also involve pathways other than canonical sGC signalling (Jacoby et al., 2018; Tooker et al., 2013; Wei et al., 2012). Due to the complex expression of NOS in vertebrates and the diversity of its functions, it has been challenging to link the neurophysiological effects of NO to signalling mechanisms and behaviour change.

In the *Drosophila* brain, NO signalling is also involved in tuning circuit activity at diverse and specific sites. In the ellipsoid body of the central complex, NO is involved in visual working memory (Kuntz et al., 2017). NO signalling also tunes the dynamics of associative memories in mushroom body circuits. NOS is expressed in PPL1-γ1pedc mushroom-body neurons where it is involved in shortening memory retention while promoting fast memory updating in response to new experiences (Aso et al., 2019).

In the whole organism context, NO signalling has often been studied in diverse marine invertebrates, where NO can regulate larval settlement and metamorphosis (Leise et al., 2001; Locascio et al., 2022; Song et al., 2021; Ueda et al., 2016; Zhang et al., 2012). While NOS- expressing neurons potentially responsible for these effects have been reported (Bishop and Brandhorst, 2007; Locascio et al., 2022), it has not been possible to link these NO-dependent effects on behaviour or life-cycle transitions to neuronal activity and function.

Overall, we still know little about how NO production relates to stimulus conditions, how it shapes circuit activity at specific neuron types, and how NO-dependent modulation relates to behaviour.

To investigate NO function in neural circuit dynamics and behaviour, here we study larvae of the marine annelid model *Platynereis dumerilii* (Ozpolat et al., 2021). *Platynereis* has emerged as a model for systems neuroscience with a toolkit that enables the combination of behavioural analysis and neuronal activity imaging with genetic manipulations. A whole-body synaptic connectome and gene expression atlases are also available and can be integrated with functional approaches in live animals thanks to the cellular-level stereotypy of larvae of the same developmental stage (Ozpolat et al., 2021; Verasztó et al., 2025; Verasztó et al., 2017; Vergara et al., 2021).

The 3-day-old larva has over 9,000 cells classified into 202 neuronal and 92 non-neuronal cell types (Verasztó et al., 2025). The synapticly connected subset of the cells in the body form a connectome of over 2,000 cells. Besides synapses, neurons in the larva also signal by volume transmission mediated by a rich repertoire of neuropeptides (Williams et al., 2017) and other modulators (Bauknecht and Jékely, 2017). The transparent and experimentally accessible *Platynereis* larvae could therefore inform how synaptic and volume signalling interact to mediate behaviour (Jékely and Yuste, 2024).

Here we uncovered an essential function for NO signalling in larval UV/violet-light avoidance behaviour. In *Platynereis* larvae, UV/violet avoidance is mediated by brain ciliary photoreceptor cells (cPRCs) and is characterised by downward swimming (Verasztó et al., 2018). The cPRCs express a ciliary-type opsin, c-opsin1 (Arendt et al., 2004) that forms a UV-absorbing bistable photopigment with an absorption maximum around 384 nm (Tsukamoto et al., 2017; Veedin Rajan et al., 2021; Verasztó et al., 2018). Upon UV/violet exposure, the cPRCs show a characteristic biphasic Ca^2+^ response that is c-opsin1 dependent. UV/violet avoidance is also c- opsin1 dependent and is defective in *c-opsin1* mutants (Verasztó et al., 2018). Here we show that *Platynereis* NOS is expressed in interneurons of the cPRC circuit and is required for UV/violet- avoidance. By combining Ca^2+^ imaging across the fully-mapped cPRC circuit (Verasztó et al., 2018) with genetic perturbations and mathematical modelling, we describe how NO tunes circuit dynamics through non-synaptic retrograde signalling to cPRCs. This delayed neuroendocrine feedback integrates UV/violet exposure to induce a short-term memory manifested in altered circuit activity and an aversive behavioural response.

## Results

### Nitric oxide synthase is expressed in interneurons of the UV-avoidance circuit

We identified a single *nitric oxide synthase* (*NOS*) gene in the *Platynereis dumerilii* genome and transcriptome data. Phylogenetic analysis of NOS proteins indicate that *Platynereis* NOS belongs to an orthology group of bilaterian NOS sequences (Figure 1—figure supplement 1). To characterise the expression pattern of *NOS*, we used in situ hybridization chain reaction (HCR) and transient transgenesis. In 2- and 3-day-old larvae, we detected *NOS* expression in four cells (two of them weakly expressing) in the apical organ region (Figure 1E and Figure 1—figure supplement 2A, B). *NOS* was also expressed in the region of the visual eyes (adult eyes) and the pigmented eyespots (Figure 1—figure supplement 2A, B). The four apical organ cells, but not the eyes, were also labelled with a transiently expressed *NOS*-reporter transgene. This transgene contains the upstream promoter sequence of the *NOS* gene that drives a membrane- targeted palmitoylated tdTomato reporter (Figure 1F). This reporter also revealed the axonal projections of these central NOS-expressing neurons. The position and morphology of the four *NOS+* cells allowed us to identify the same four cells as four interneurons (INNOS) in our 3-day- old whole-body *Platynereis* volume EM data (Verasztó et al., 2025; Verasztó et al., 2018; Williams et al., 2017) (Figure 1C, D). In the synaptic connectome, the INNOS cells are postsynaptic to the UV-sensory cPRCs and presynaptic to the INRGWa neurons (cholinergic interneurons that express an RGW neuropeptide), which are also cPRC targets (Figure 1C, H). The projections of INNOS cells are segregated into input (dendritic) and output (axonal) compartments, with cPRC inputs occurring on the dendritic part in the dense neurosecretory plexus (Figure 1D and Figure 1—figure supplement 3A). Immunostaining with an affinity-purified antibody specific to *Platynereis* NOS revealed that the protein is restricted to the dendritic compartment of INNOS neurons (Figure 1G). INNOS output-synapses form in the more ventral projection region (Figure 1D and Figure 1—figure supplement 3). The second type of interneurons of the cPRC circuit, the INRGWa neurons, synapse on the head serotonergic ciliomotor neurons (Ser-h1). These in turn synapse on the cholinergic MC ciliomotor neuron and the multiciliated cells of the prototroch ciliary band (Figure 1C, H) (Verasztó et al., 2017).

**Figure 1.**
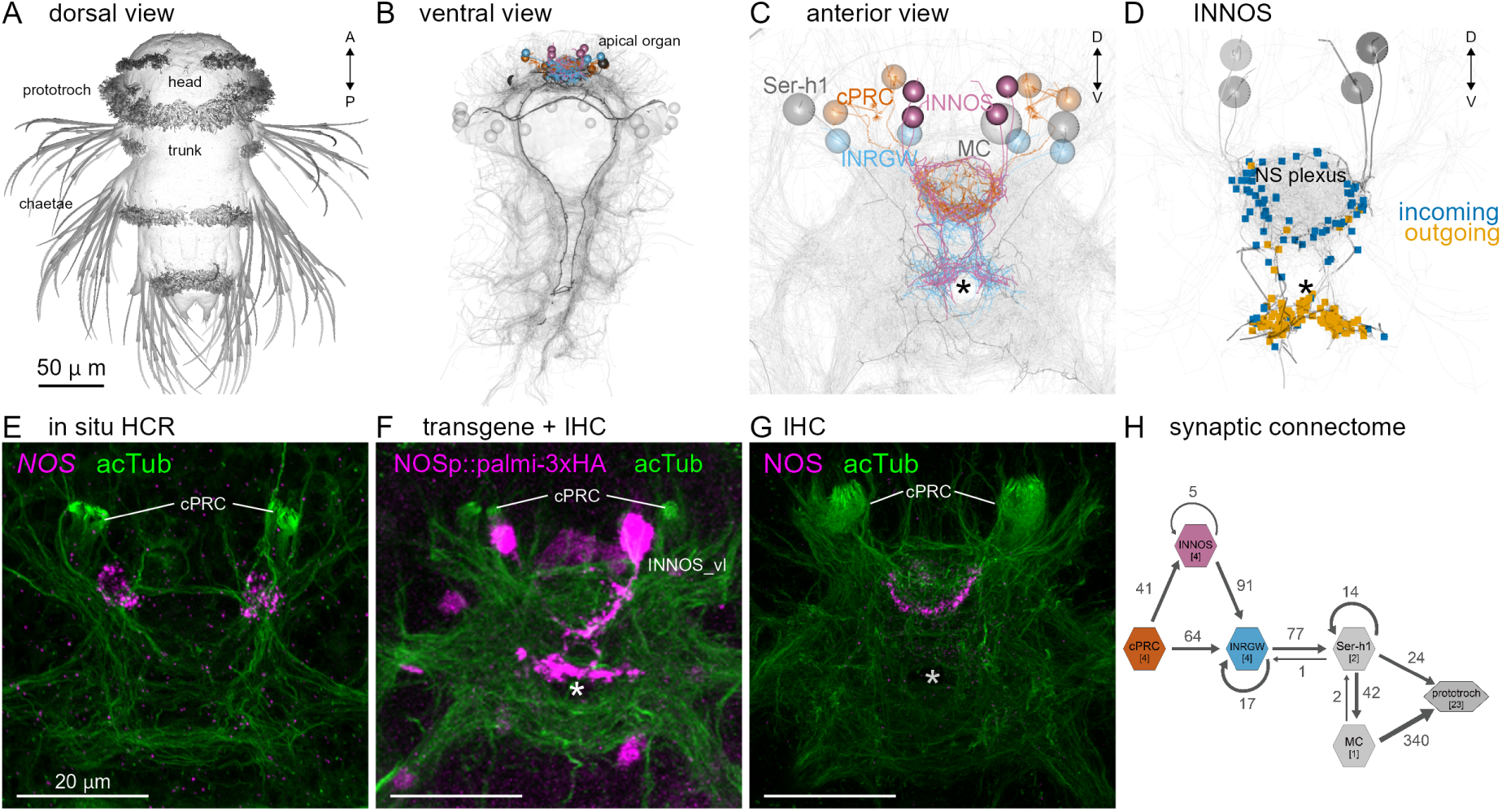
Identification of *NOS*-expressing interneurons (INNOS) within the cPRC circuit. **(A)** Scanning electron microscopy image of a 3-day-old *Platynereis* larva. **(B, C)** Volume rendering of the neuron types (cPRC, INNOS, INRGWa, Ser-h1 and MC) in the cPRC circuit reconstructed from a whole-body transmission electron microscopy volume of a 3-day-old larva. Neurite skeletons are shown with cell-body positions represented by spheres. Projections of all neurons in the body are shown in grey to highlight the neuropils. The outline of the yolk is also indicated in grey. In B, nuclei positions of the prototroch head ciliary band are shown as grey spheres. **(D)** Volume rendering of the four INNOS neurons with incoming and outgoing synapses. **(E)** Expression of the *NOS* gene detected by in situ HCR (magenta) in a 2-day-old larva (anterior view). Antibody staining for acetylated α-tubulin (acTub: green) highlights cPRC cilia and the neuropil. **(F)** Expression of a membrane-targeted reporter driven by the *NOS* regulatory region (NOSp::palmi-3xHA-Tomato; magenta) labelled with an anti-HA antibody in a 2-day-old larva (anterior view). Antibody staining for acetylated α-tubulin (acTub; green) highlights cPRC cilia and the neuropil. **(G)** Immunostaining for NOS (magenta), co-stained for acetylated α-tubulin (acTub; green). **(H)** Synaptic wiring diagram of the cPRC circuit. Hexagons represent cell groups, with the number of cells per group shown in square brackets. Arrows represent the summed number of synaptic contacts between cell groups. Arrow thickness is proportional to the number of synapses.

### Nitric oxide is produced during UV/violet stimulation of the cPRCs

The expression of *NOS* in the INNOS interneurons in the cPRC circuit suggests that NO signalling may be involved in UV/violet-avoidance. To test this, first we asked whether NO is produced during UV/violet stimulation of the larvae. We injected the fluorescent NO-reporter DAF-FM into zygotes and imaged 2-day-old larvae while exposing the region of cPRC cilia to 405 nm violet light (Figure 2A, B). To mark cell outlines, we coinjected mRNA encoding a red-fluorescent reporter (RGECO), allowing the identification of the cPRCs. Following light stimulation in the region of the ramified cilia of the cPRCs, we detected an increase in DAF-FM fluorescence in the anterior neurosecretory neuropil, the region of INNOS projections (Figure 2C). This increase did not occur in larvae in which we illuminated a control area (Figure 2D).

**Figure 2.**
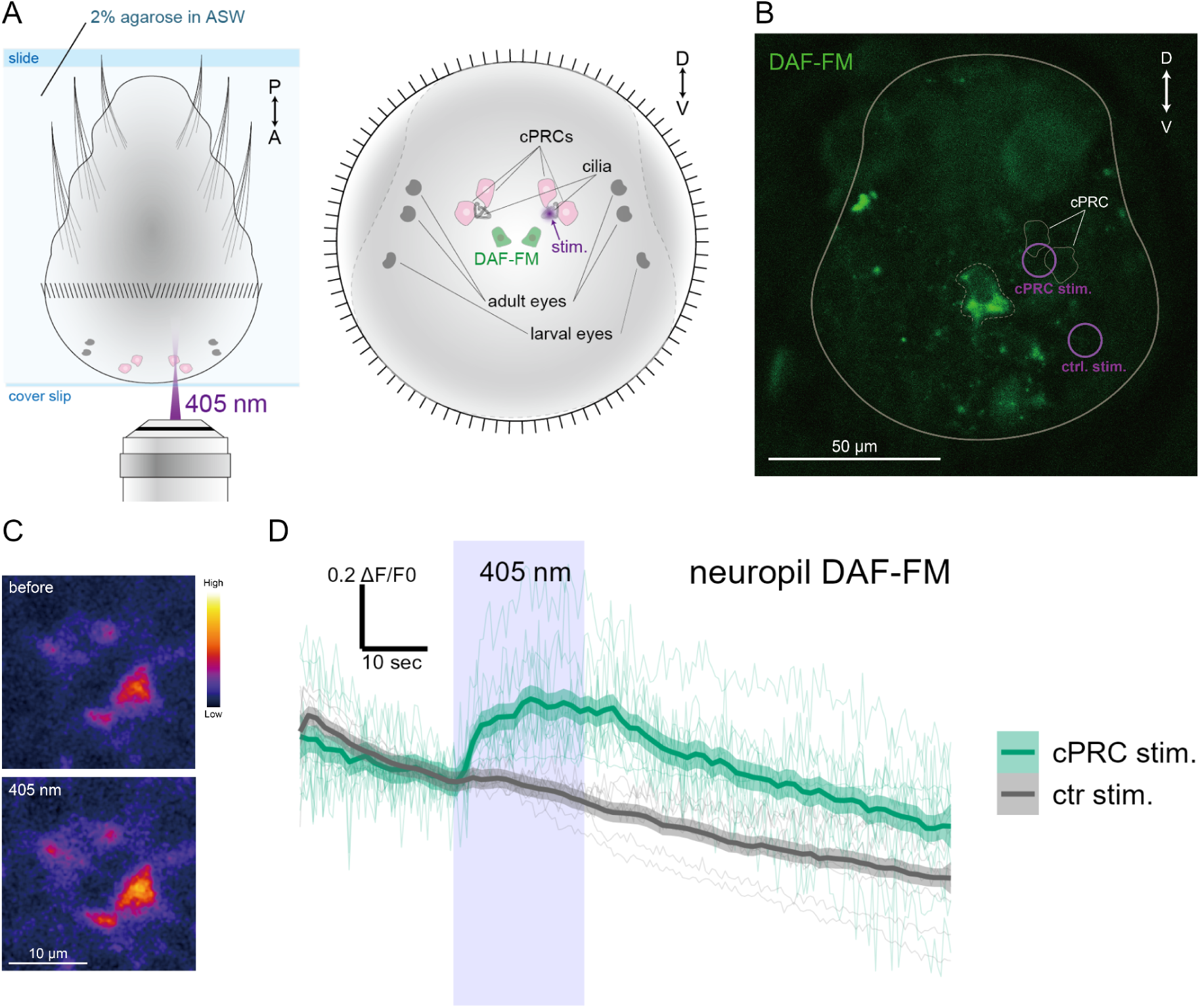
NO produced by UV/violet stimulation to cPRCs. **(A)** Left: The larvae were embedded in 2% agarose seawater with the anterior side down. Right: Changes in DAF-FM fluorescence intensities were examined when the ciliary region of the cPRC was stimulated with a 405 nm laser.**(B)** DAF-FM fluorescence in the region of the neurosecretory neuropil. The white line indicates the outline of the larva. The dashed line corresponds to the area where fluorescence was quantified. The circles indicate the location of cPRC and control stimulation. The cPRCs are marked by thin lines. **(C)** DAF-FM fluorescence before and during 405 nm light stimulation. **(D)** Changes in DAF-FM fluorescence over time during 405 nm stimulation of the cPRCs or a control area (ctr stim.). The purple box indicates the duration of 405 nm stimulation. Individual normalized traces (ΔF/F0) are shown as thin lines. Thick lines show the mean value with 0.95 confidence intervals. N = 9 larvae for control and 11 for cPRC stimulation. Figure 2 – source data 1. DAF-FM fluorescence reads.

### Nitric oxide signalling mediates UV-avoidance behaviour

We next tested whether NO signalling regulates UV/violet avoidance. To achieve this, we generated two *Platynereis NOS* knockout lines using the CRISPR/Cas9 method. We recovered two deletions (NOSΔ11/Δ11 and Δ23/Δ23), both frame-shift mutations leading to an early stop codon and thus likely representing null alleles (Figure 3—figure supplement 1A). We could establish a homozygous line for both mutations indicating that NOS is not an essential gene in *Platynereis* (Figure 3—figure supplement 1B, C).

To quantify UV avoidance, we recorded the trajectories of freely swimming wild type and mutant larvae in vertical columns, illuminated laterally from two opposite sides with 395 nm UV light (Figure 3A and Figure 3—figure supplement 2A, B). As previously shown, wild-type larvae swim downward following non-directional UV/violet light stimulation (Verasztó et al., 2018). In contrast, both 2- and 3-day-old homozygous *NOS*-mutant larvae showed a significantly diminished UV-avoidance response (Figure 3B, C and Figure 3—figure supplement 1C, D). This phenotype is similar to the defective UV-avoidance of *c-opsin1* mutant larvae (Verasztó et al., 2018) and reveals a requirement for NOS in UV-avoidance behaviour. Wild type but not mutant larvae also showed an increase in ciliary beat frequency and swimming speed under UV light (Figure 3—figure supplement 3). We also tested directional phototaxis, by exposing larvae to 480 nm directional collimated light from the top of the column (Figure 3—figure supplement 2A). 3-day-old but not 2-day-old *NOS*-mutant larvae also showed reduced phototactic behaviour. Given that phototaxis in 1 and 2-day-old trochophore larvae is mediated by their non-visual eyespots (Jékely et al., 2008) and in 3-day-old nectochaete larvae by the visual eyes (Randel et al., 2014), these data suggest a function for *NOS* in the visual eyes (Figure 3D and Figure 3— figure supplement 1E).

**Figure 3.**
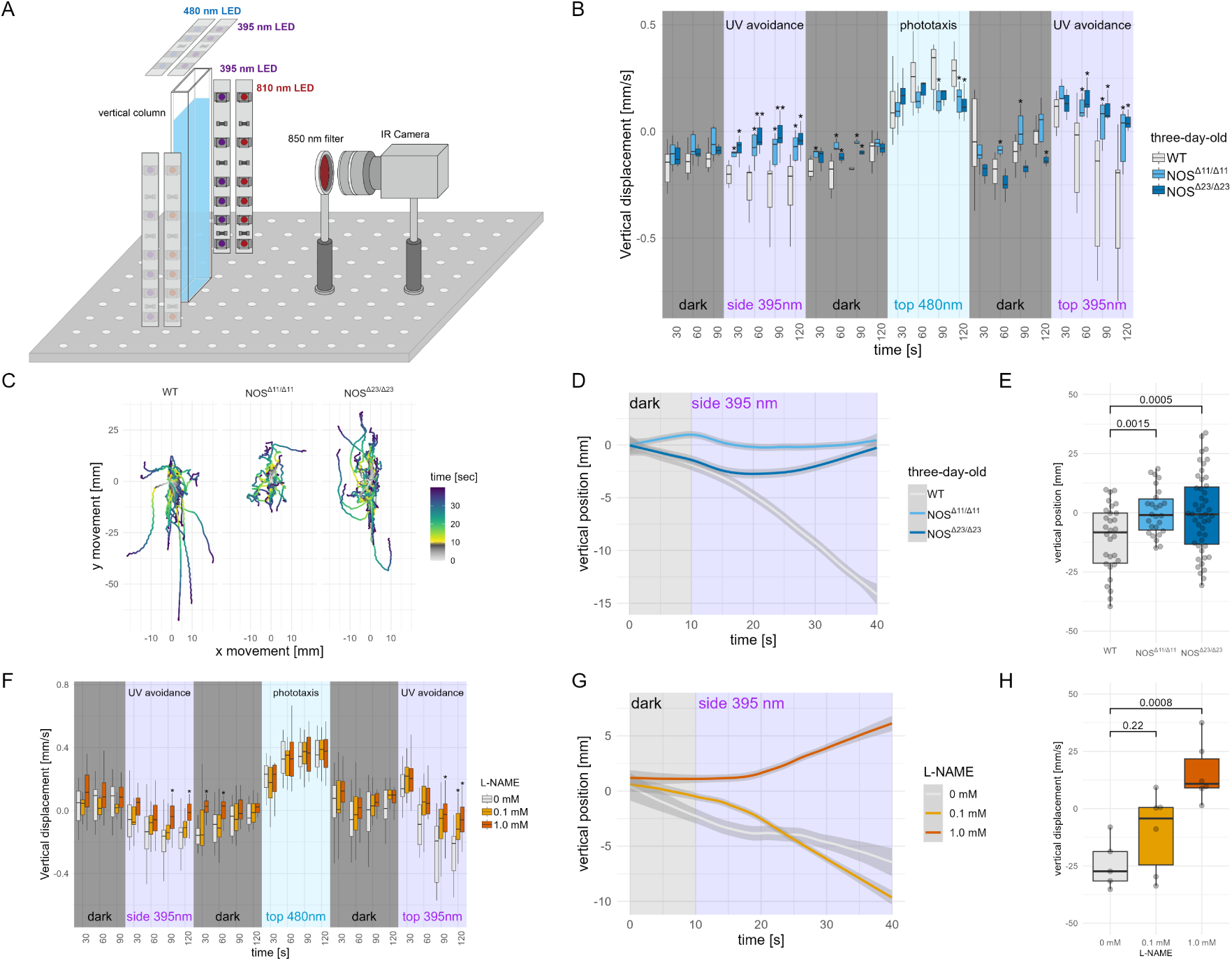
NOS is required for UV avoidance in *Platynereis* larvae. **(A)** Schematic diagram of the set-up of the behavioural experiment. **(B)** Vertical displacement in 30 sec bins of wild type (10 batches) and mutant (*NOSΔ11/Δ11*, 5 batches and *NOSΔ23/Δ23*, 7 batches) 3-day-old larvae stimulated with 395 nm light from the side, 488 nm light from the top and 395 nm light from the top. One-way ANOVA with Dunnett’s multiple-comparison test are shown. *P < 0.05, **P < 0.01. **(C)** Swimming trajectories of wild type (WT, n=32) and *NOS* mutant (*NOSΔ11/Δ11*, n=26 and *NOSΔ23/Δ23*, n=47) 3-day-old larvae. All trajectories start at 0 x and y position and time 0 corresponding to 10 sec before the onset of 395 nm stimulation from the side. **(D)** Vertical position of batches of wild type and mutant 3-day-old larvae over time under 395 nm UV stimulation. The starting position of each larval trajectory was set to 0. **(E)** Comparison of vertical position of wild type and mutant 3-day-old larvae after 30 seconds of UV stimulation. One-way ANOVA with Dunnett’s multiple-comparison test are shown. The wild type (WT, n=32) and *NOS* mutant (*NOSΔ11/Δ11*, n=26 and *NOSΔ23/Δ23*, n=47) larvae. **(F)** Vertical displacement in 30 sec bins of control (15 batches) and L-NAME-treated (0.1 and 1 mM, 15 batches each) 3-day-old larvae stimulated with 395 nm light from the side, 488 nm light from the top and 395 nm light from the top. One-way ANOVA with Dunnett’s multiple-comparison test are shown. *P < 0.05. **(G)** Vertical position of batches of control and L-NAME-treated (0.1 and 1 mM) 3-day-old larvae over time under 395 nm UV stimulation. The starting position of each larval trajectory was set to 0. **(H)** Comparison of vertical position of control (n=6) and L-NAME-treated (0.1 and 1 mM, n=6 each) 3-day-old larvae after 30 seconds of UV stimulation. One-way ANOVA with Dunnett’s multiple-comparison test are shown. Figure 3 – source data 1. Vertical displacement data for wild type and *NOS* mutant larvae. Figure 3 – source data 2. Swimming trajectories of wild type and *NOS* mutant larvae. Figure 3 – source data 3. Vertical position of wild type and *NOS* mutant larvae. Figure 3 – source data 4. Vertical position of wild type and *NOS* mutant larvae after 30 sec UV exposure. Figure 3 – source data 5. Vertical displacement of control and L-NAME-treated larvae. Figure 3 – source data 6. Vertical position of control and L-NAME-treated larvae. Figure 3 – source data 7. Vertical position of control and L-NAME-treated larvae after 30 sec UV exposure.

To distinguish between an acute and developmental function of NOS in light responses, we also tested larvae exposed to the NOS inhibitor L-NAME (Figure 3F). Larvae incubated for 5 min in 0.1 mM or 1 mM L-NAME showed a dose-dependent inhibition of UV avoidance (Figure3G, H). In contrast, phototaxis was not affected (Figure 3F). Finally, as a non-visual behaviour, we also quantified the emergence of sexually mature (epitok) swimming worms from their tubes. The rhythmicity of this behaviour and sexual maturation is controlled by an artificial lunar cycle in the lab. *NOS* mutant worms showed periodic emergence, similar to wild-type worms (Figure 3— figure supplement 4). Overall, our results indicate an acute requirement for NOS signalling in UV-avoidance and a possible indirect, developmental role in the visual system, reminiscent of the function of NO signalling in *Drosophila* eye development (Gibbs and Truman, 1998).

### NO retrograde signalling drives a late-phase cPRC response after UV/violet stimulation

To investigate how NO signalling alters the dynamics of the cPRC circuit, we carried out Ca^2+^ imaging experiments. We ubiquitously expressed the Ca^2+^ sensor GCaMP6s in larvae and imaged Ca^2+^ signals during 405 nm stimulation of the cPRCs. As we have shown previously, a 20 sec local stimulation of cPRC cilia led to a transient increase in cPRC Ca^2+^ levels, followed by a transient decrease (Verasztó et al., 2018). After ∼20-sec, Ca^2+^ levels in cPRCs were raising again, reaching higher levels than at the start of the stimulus – a response that may involve cPRC depolarisation (Figure 4A, B). This activation phase occurs after the 20 sec stimulation period and is likely due to a delayed neuroendocrine feedback (Verasztó et al., 2018). To determine whether NO mediates such a feedback, we repeated the experiment in *NOS*-mutant larvae. While we detected the initial activation phase followed by inhibition, in homozygous *NOS*-mutants for both CRISPR alleles this was not followed by delayed activation. Instead, Ca^2+^ levels dropped to a low steady-state level (Figure 4A, B). We thus identified a requirement for NO signalling in the late-phase activation of cPRCs.

**Figure 4.**
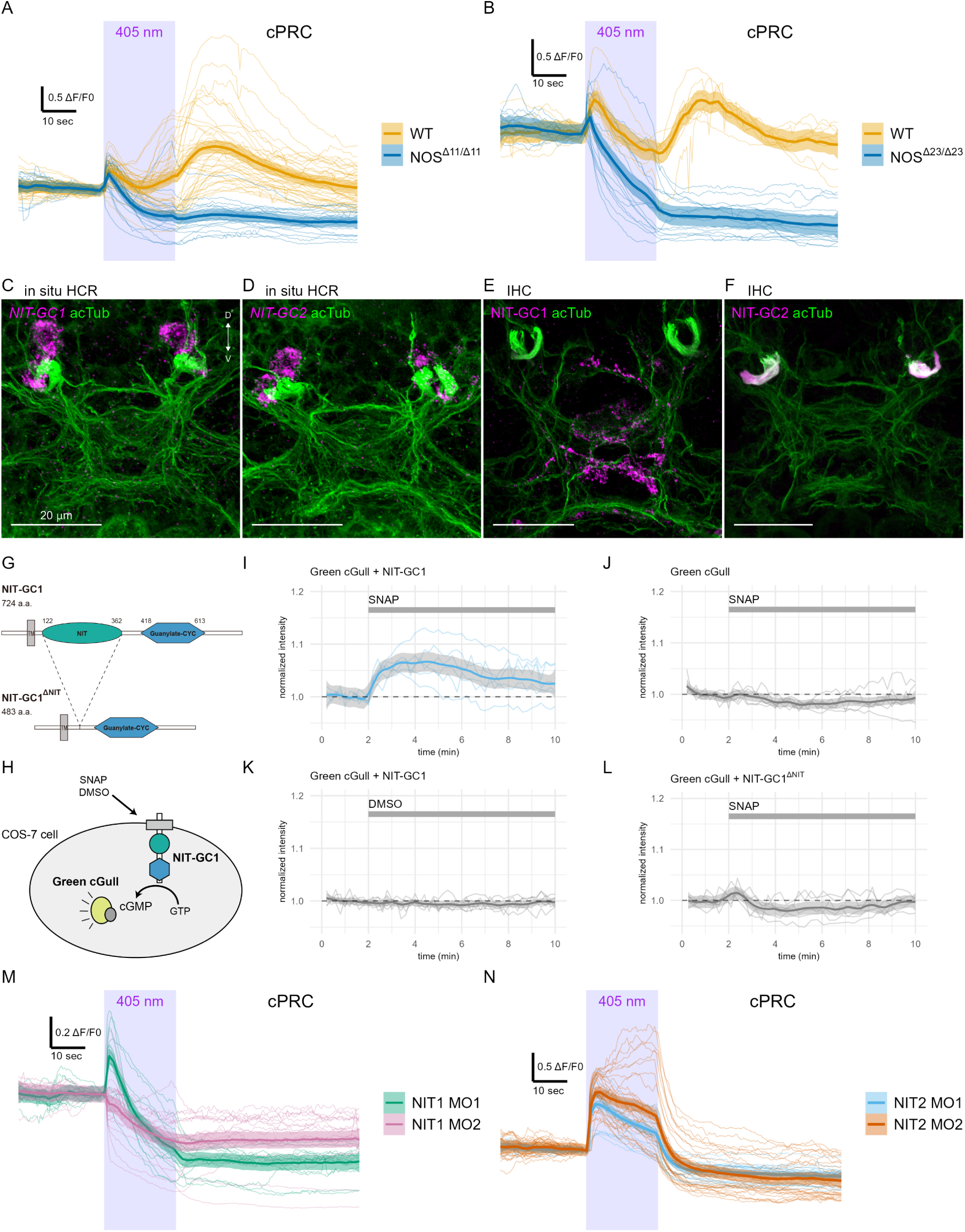
NOS and two NIT-GCs shape Ca^2+^ signals during cPRC UV/violet response. **(A, B)** GCaMP6s signals in cPRCs in wild type and *NOS* mutant (A, *NOSΔ11/Δ11*, B, *NOSΔ23/Δ23*) larvae during 405 nm light stimulation. **(C, D)** In situ HCR for (C) *NIT-GC1* and (D) *NIT-GC2* (magenta) in 3-day-old *Platynereis* larvae. Larvae were co-stained with an antibody against acetylated α-tubulin to label cPRC cilia and the neuropil (green). **(E, F)** Immunostaining for (E) NIT-GC1 and (F) NIT-GC2 (magenta), co-stained for acetylated α- tubulin (green). **(G)** The domain structure of *Platynereis* NIT-GC1 and the truncated NIT-GC1ΔNIT protein lacking the NIT domain. A predicted transmembrane region (TM) is shown in grey. (H) Schematic of the cell-based assay to detect cGMP production following the addition of an NO donor SNAP or DMSO as control. **(I-L)** Green cGull fluorescence over time for the four conditions tested. Individual responses and their mean with 0.95 confidence interval are shown (n > 6 cells). Intensities are normalized (ΔF/F0). The indicated chemicals were added at 2 min after the start of imaging (grey bars). **(M, N)** GCaMP6s signals in cPRCs in (M) NIT-GC1 and (N) *NIT-GC2*-morphant larvae during 405 nm light stimulation. Individual responses and their mean with 0.95 confidence interval are shown. Figure 4 – source data 1. GCaMP6s data for cPRCs in wild type during 405 nm stimulation. Figure 4 – source data 2. GCaMP6s data for cPRCs in *NOS* mutant larvae during 405 nm stimulation. Figure 4 – source data 3. Green cGull fluorescence data of cells with NIT-GC1 after SNAP treatment. Figure 4 – source data 4. Green cGull fluorescence data of cells without NIT-GC1 after SNAP treatment. Figure 4 – source data 5. Green cGull fluorescence data of cells with NIT-GC1 after DMSO treatment. Figure 4 – source data 6. Green cGull fluorescence data of cells with NIT-GC1ΔNIT after SNAP treatment. Figure 4 – source data 7. GCaMP6s data for cPRCs in NIT-GC1 morphant larvae during 405 nm stimulation. Figure 4 – source data 8. GCaMP6s data for cPRCs in NIT-GC2 morphant larvae during 405 nm stimulation.

### Two unconventional guanylate cyclases are expressed in the cPRCs

Next we aimed to identify the NO receptor in the cPRCs. NO generally acts via soluble guanylate cyclases (sGC), belonging to the guanylate cyclase family with a CYC domain (PFAM domain: PF00211). NO binding to the heme group of sGC leads to increased cyclic guanosine monophosphate (cGMP) production. Analysis of sGCs in *Platynereis* showed that these genes were not expressed in cPRC or INNOS cells, we only detected expression of an sGCβ subunit in the INRGWa cells (Figure 6B).

**Figure 5.**
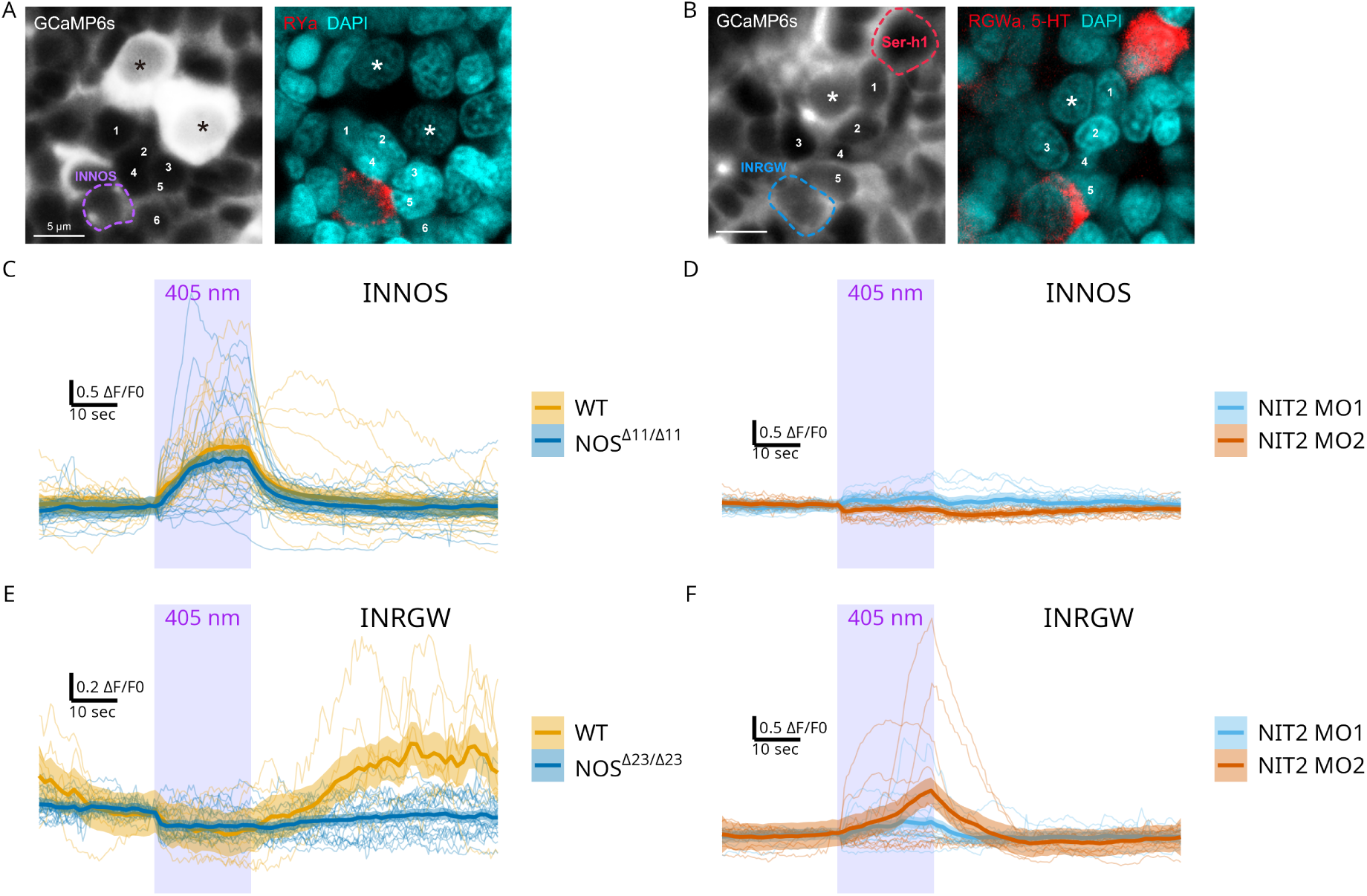
NOS- and NIT-GC2-dependent dynamics of the cPRC circuit. **(A, B)** GCaMP6s imaging from cPRCs and INNOS cells (left panels) followed by on-slide immunostaining for (A) RYamide to label INNOS and (B) RGWamide+serotonin to label INRGWa and Ser-h1 (red). Nuclei are stained with DAPI (cyan). Asterisks indicate cPRC nuclei. Numbers mark the same cells in the GCaMP and immunostaining images matched by position. **(C)** GCaMP6s fluorescence in INNOS cells in wild type (WT) and *NOSΔ11/Δ11* mutant larvae during 405 nm stimulation of the cPRC cilia. **(D)** GCaMP6s fluorescence in INNOS cells in *NIT-GC2*- morphant larvae during 405 nm stimulation. **(E)** GCaMP6s fluorescence in INRGWa cells in wild type and *NOSΔ23/Δ23* mutant larvae during 405 nm stimulation. **(F)** GCaMP6s fluorescence in INRGWa cells in *NIT- GC2*-morphant larvae during 405 nm stimulation. Figure 5 – source data 1. GCaMP6s fluorescence data forINNOS in wild type and *NOS* mutant larvae. Figure 5 – source data 2. GCaMP6s fluorescence data for INNOS in wild type and NIT-GC2 morphant larvae. Figure 5 – source data 3. GCaMP6s fluorescence data for INRGWa in wild type and *NOS* mutant larvae. Figure 5 – source data 4. GCaMP6s fluorescence data for INRGWa in wild type and NIT-GC2 morphant larvae.

**Figure 6.**
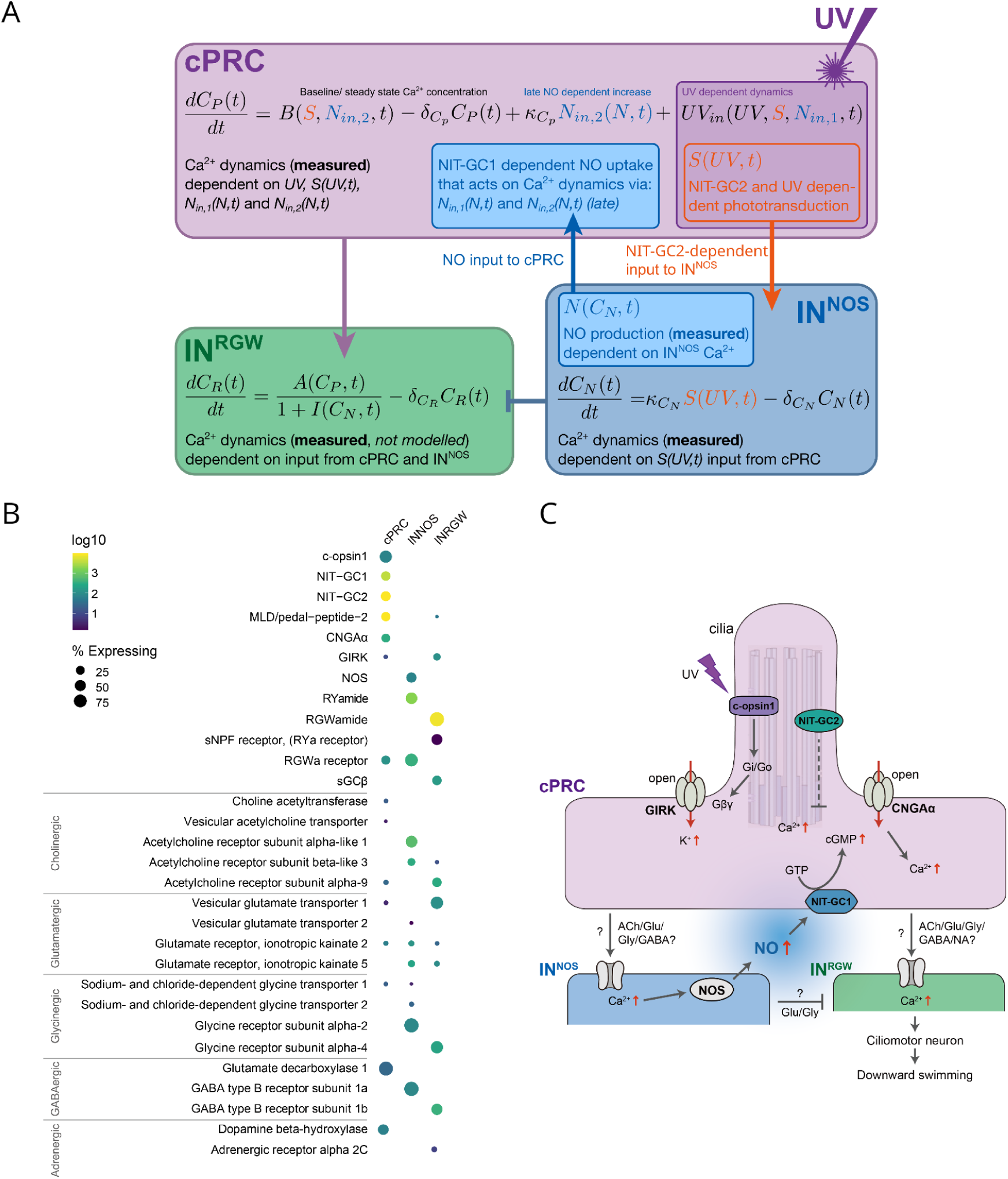
Mathematical modelling and signalling mechanisms of the cPRC circuit. **(A)** Diagram of the mathematical model with the componenets, interactions, parameters and equations used to model Ca^2+^ dynamics. **(B)** Dot plot of genes (columns) expressed in three types of cells (rows) in the cPRC circuit using single cell RNA-Seq. The size of the dots is expressed in proportion to the percentage of cells expressing that gene relative to all cells. The colours represent the normal logarithm of the number of transcripts in the cells expressing the gene. **(C)** Schematic diagram of the signalling pathway of the cPRC circuit, focusing on the NO feedback. Figure 6 – source data 1. TPM values for each gene and the percentage of expressed genes.

Recently, Moroz and coworkers reported an atypical but widely conserved family of guanylate cyclases with a NIT (nitrite/nitrate sensing) domain (PF08376) (NIT-GC) as potential mediators of NO signalling (Moroz et al., 2020). To identify NIT-GCs in *Platynereis*, we searched transcriptome resources and retrieved 15 potential NIT-GC homologs (Figure 4—figure supplement 1 and 2).

To analyse the relationship of these sequences to metazoan NIT-GCs, we retrieved protein sequences with a CYC domain from the transcriptome and genome databases of 45 metazoan and 2 choanoflagellate species. We carried out cluster analysis and phylogenetic reconstruction on a group of membrane-bound guanylate cyclases with sGCs as an outgroup. In agreement with Moroz et al. (Moroz et al., 2020), we found a group of GCs with NIT domains with representatives in placozoans, cnidarians, some ecdysozoans, echinoderms, and lophotrochozoans. The 15 *Platynereis* sequences belonged to several deeply diverged clades in the phylogenetic tree (Figure 4—figure supplement 1 and 2).

To characterise the expression of NIT-GCs, we used previously published spatially-mapped single-cell transcriptome data (Achim et al., 2015). Among the 15 NIT-GCs, two showed high and specific expression in the cPRCs and one was expressed in the INNOS cells (Figure 4—figure supplement 2). In the single-cell data, we could identify the cPRCs by the specific expression of the markers *c-opsin1* and the *pedal-peptide2 neuropeptide precursor* (*MLD proneuropeptide*) (Arendt et al., 2004; Williams et al., 2017) (Figure 4—figure supplement 3A). The INNOS cells were identified by NOS expression and spatial mapping in the brain (Achim et al., 2015). We decided to focus on two NIT-GCs expressed in the cPRCs and with a full-length sequence, *NIT- GC1* and *NIT-GC2*. To confirm the single-cell data, we first carried out in situ hybridisation chain reaction (HCR) with probes for *NIT-GC1* and *NIT-GC2* mRNA. Both genes were specifically expressed in the four cPRCs, as confirmed by co-labeling with an acetylated α-tubulin antibody and with an HCR probe against *pedal peptide 2/MLD proneuropeptide* (Figure 4C, D and Figure 4— figure supplement 3A-C). To analyse the subcellular localisation of NIT-GC1 and NIT-GC2 at the protein level, we raised and affinity-purified polyclonal antibodies against a specific peptide sequence from both proteins. In immunostainings, we found that NIT-GC1 was localise to the region corresponding to the axonal projections of the cPRCs in the anterior neurosecretory plexus (Figure 4E). In contrast, NIT-GC2 specifically labelled the ramified sensory cilia of the cPRCs (Figure 4F). These different subcellular localisations suggest that the two NIT-GCs are involved in different intracellular signalling processes in the ciliary and axonal regions of the cPRCs.

### NIT-GC1 produces cGMP in an NO-dependent manner

To further characterise these atypical guanylate cyclases, we focused on NIT-GC1 and carried out in vitro experiments. In bacteria, NIT domains are thought to regulate cellular functions in response to intra- or extracellular nitrate and nitrite. NIT-GC1 has a NIT domain and a highly conserved cyclase domain that is expected to catalyse cGMP synthesis (Figure 4G). The NIT domain may render NIT-GC1 dependent on NO signals. To test this, we co-expressed the cGMP indicator Green cGull (Matsuda et al., 2016) and NIT-GC1 in cultured COS-7 (monkey kidney) cells, a cell line with minimal endogenous sGC activity (Matsuda et al., 2016). For balanced expression, we used a single plasmid with the two open-reading frames separated by the 2A self-cleaving peptide (Figure 4H). Application of the NO donor SNAP lead to increased Green cGull fluorescence, an effect we did not observe when cells were exposed to DMSO or when Green cGull was expressed alone (Figure 4I-K). To test whether this effect is dependent on the NIT domain, we also tested a deletion construct of NIT-GC1 lacking the NIT domain (Figure 4G). Cells expressing this construct and Green cGull did not show an increased fluorescence of the cGMP reporter when exposed to SNAP (Figure 4L). These results indicate that NIT-GC1 is able to catalyse cGMP production in an NO-dependent manner and this function requires the NIT domain. These results establish NIT-GC1 as a biochemical sensor of NO or its derivatives.

### NIT-GC1 is required for NO-mediated retrograde signalling to cPRCs during the UV response

To test the in vivo function of NIT-GC1 and NIT-GC2 in cPRC responses, we combined Ca^2+^ imaging with morpholino-mediated knockdowns. We used two translation-blocking morpholinos for each *NIT-GC* gene and tested knockdown efficiency by immunostaining with the NIT-GC1 and NIT-GC2 antibodies (Figure 4—figure supplement 3D,E). For both genes, the morpholinos led to a strong reduction in the respective antibody signal, confirming efficient knockdown and antibody specificity (Figure 4—figure supplement 3F).

In NIT-GC1 morphant larvae, the delayed activation of cPRCs following 405 nm stimulation did not occur (Figure 4M). This phenotype is similar to the phenotype of *NOS* mutants suggesting that NIT-GC1 acts as the NO sensor in cPRCs to drive their delayed activation. This could occur via increased cGMP production and the opening of a cyclic-nucleotide-gated ion channel (CNG) specific to cPRCs (Tosches et al., 2014). *NIT-GC2*-morphant larvae, in contrast, showed a step-up increase in Ca^2+^ following light stimulation (Figure 4N). The Ca^2+^ signal decayed during stimulation and was off after light off. These data support an essential role for ciliary-localised NIT-GC2 in suppressing cPRC Ca^2+^ following its transient rise at stimulus onset. Overall, these knockdown experiments revealed different signalling mechanisms for the two NIT-GCs that may be due to their different subcellular localisations.

### NO signalling shapes the dynamics of the cPRC circuit

To investigate how NO and NIT-GC signalling influence the dynamics of the cPRC circuit, we imaged Ca^2+^ signals from the INNOS, INRGWa and Ser-h1 neurons in wild type, mutant and morphant larvae. To unambiguously identify the neurons from which we recorded Ca^2+^ signals, we developed an on-slide immunostaining method (Figure 5—figure supplement 1A). We used the cell-specific antibody markers against RYamide (INNOS) (Figure 5—figure supplement 1B-D), RGWamide (INRGWa) and serotonin (Ser-h1) (Conzelmann et al., 2011) to immunostain agar-embedded larvae following Ca^2+^ imaging. Based on the position of the nuclei, we could correlate live and fixed samples at a single-cell precision (Figure 5A, B). Due to the stereotypy of the larvae, we could also identify neurons based on their position and Ca^2+^ activity in activity-correlation maps (Figure 5 – figure supplement 2A).

We first quantified the responses of the INNOS and INRGWa interneurons during 405 nm stimulation of the cPRCs. In both wild type and *NOS*-mutant larvae, INNOS cells showed an increase in Ca^2+^ during stimulation (Figure 5C). In contrast, the INNOS response was flat or slightly negative in *NIT-GC2*-morphant larvae (Figure 5D) revealing an essential role for NIT-GC2- mediated cPRC suppression in INNOS activation. The INRGWa cells were initially inhibited during cPRC stimulation, followed by a delayed activation paralleling the second activation phase of the cPRCs. This late INRGWa response was lacking in *NOS*-mutants (Figure 5E). In *NIT-GC2*-morphants, INRGWa cells showed a transient increase in Ca^2+^ that decayed after light off and a delayed activation was not present (Figure 5F).

Next, we imaged Ca^2+^ signals from Ser-h1 neurons in wild type and *NOS* mutant larvae. Ser-h1 cells showed mixed responses. In some larvae the activation correlated with cPRC activity, including a reduction in Ca^2+^ during stimulation followed by a rebound. In other larvae, Ser-h1 did not show the post-stimulus rebound. In *NOS* mutants Ser-h1 in most larvae showed no post- stimulus rebound (Figure 5 – figure supplement 2B). These data suggest that during 405 nm stimulation the Ser-h1 cells can be activated, and this regulation is NO-dependent. This pattern is expected to increase ciliary activity in the prototroch but not in the other ciliary bands (with no Ser-h1 synapses), triggering faster swimming under UV light. The variability may be due to the endogenous rhythmic activity and regular inhibition of Ser-h1 neurons that are part of a larger ciliomotor circuit (Verasztó et al., 2017).

### Mathematical modelling of cPRC-circuit dynamics

To further analyse the dynamics of responses to UV light and formally describe cPRC phototransduction, synaptic connections, and NO retrograde signalling, we developed a mixed cellular-circuit-level mathematical model. We used our Ca^2+^ imagining data of cPRC, INNOS and INRGWa cells collected in wild type, *NOS* knockout and *NIT-GC2* morphant larvae to formulate assumptions that are the basis of the proposed model.

We modelled a c-opsin1-dependent Ca^2+^-response to UV in the cPRCs (Figure 6A). This response depends on UV-dependent phototransduction, inhibition from NIT-GC2 activity, and NO-dependent activation. Biochemically, the pathways could include — besides c-opsin1 and NIT-GC2 — Gβγ-biased signalling to a GIRK channel (Tsukamoto and Kubo, 2023a), cGMP, a cGMP- gated CNGAα channel, and other unknown components.

The NIT-GC2-dependent branch of the phototransduction signal is one input to Ca^2+^ levels in the cPRC and leads to the activation (Ca^2+^ increase) of the postsynaptic INNOS cells (via synaptic transmission) (Figure 6A). In the INNOS cells, we assumed that excitatory synaptic input from cPRC (or rebound from tonic inhibition) leads to a rise in Ca^2+^ leading to NO production via NOS. We do not explicitly incorporate diffusion of NO into the model.

The effect of NO in cPRC cells is captured by additional terms that describe a NIT-GC1 and NO- dependent increase in Ca^2+^. This step is also dependent on the previously mentioned second branch phototransduction pathway. The effect on calcium may be mediated by cGMP, not modelled here.

Based on the synaptic connectome, we inferred a feedforward coupling between the INNOS and INRGWa cells. We modelled this as an inhibitory connection, given the decrease in Ca^2+^ in the INRGWa cells at the onset of simulation. This inhibition is also dependent on NIT-GC2 (Figure 5E). We further assumed the existence of direct excitatory coupling between the NIT-GC1- dependent NO response in cPRC and INRGWa Ca^2+^ to account for the late effects of cPRC on INRGWa (Figure 6A).

Finally, in all variables, we assumed a linear decay and constant production to set a steady state (Figure 6A).

Since the aim of the model is to capture the normalised fluorescence data, the model is nondimensionalised. UV stimulation is modelled as a square pulse. To find model parameters producing an output fitting the experimental data we employed a global optimisation method known as a genetic algorithm (GA).

Through this optimisation procedure, we found parameters that gave a fit to our experimental data under all conditions (Figure 6–figure supplement 1-5). We could fit to mean Ca^2+^ profiles but also individual Ca^2+^ transients (Figure 7; Figure 6 – figure supplement 1). In addition, we could fit our model to the experimentally characterised NO signals in the neuropil (Figure 2D; Figure 6 – figure supplement 2).

**Figure 7.**
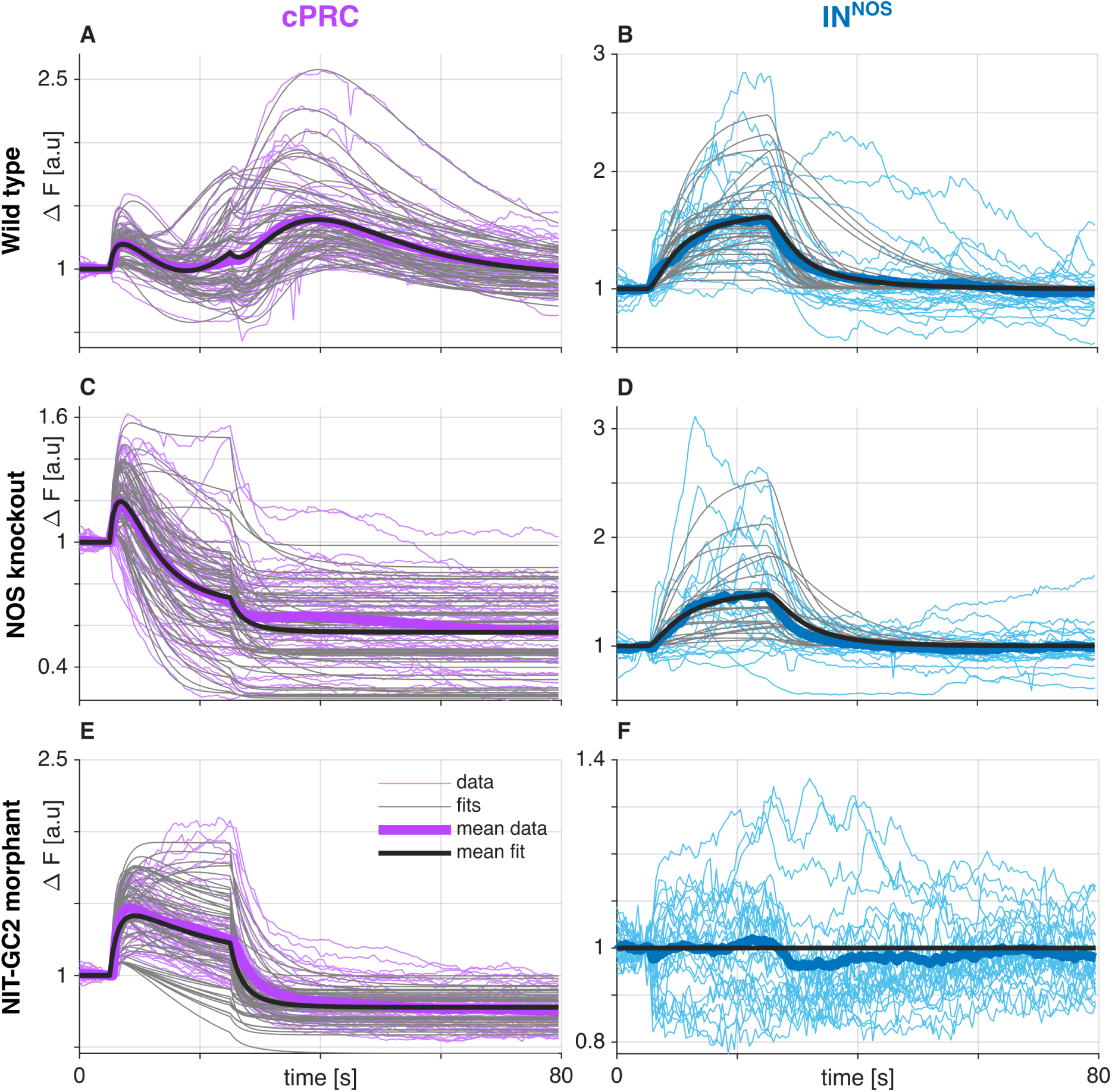
Simulated Ca^2+^ traces for parameter sets fitted independently to each Ca^2+^-recording collected in wild type, *NOS* knockout, and *NIT-GC2* morphant larvae. (A, B) Simulated Ca^2+^ traces in cPRC (A) and INNOS (B) cells in the wild-type condition. (C, D) Simulated Ca^2+^ traces in cPRC (C) and INNOS (D) cells in the *NOS* knockout condition. (E-F) Simulated Ca^2+^ traces in cPRC (E) and INNOS (H) cells in the *NIT-GC2*-morphant condition. Thin coloured curves indicate individual recordings (cPRC - purple and INNOS - blue), thin grey curves indicate simulated Ca^2+^ traces based on model parameters fitted to the individual recording, thick coloured curves indicate averages of the recordings, and thick black curves represent the average of the fits. Figure 7 – source data 1. Best fitted parameters `best_pars.mat`. Figure 7 – source data 2. Recorded traces (as used in earlier figures) `wt_ko_mo_data.mat`. Figure 7 – source data 3. script to generate the figure `fig_all_fits.m`

Our model only includes a minimal set of parameters and assumptions of interactions that are required to describe the dynamics of cPRC, INNOS and INRGWa in wild type and loss-of-function conditions (Figure 6A). The model highlights that cPRC phototransduction employs distinct pathways that operate on different time scales and differentially influence cPRC Ca^2+^ levels. The coupling between cPRCs and interneurons also requires different signalling mechanisms, either through distinct neurotransmitters or receptors expressed in the different cells.

Many of the couplings in our model are constrained (e.g. UV leads to INNOS activation) and e.g. reversing the sign of some of these couplings would not arrive at the same result. Several earlier variants of the model could not be fit to the data, The model is thus well constrained by our physiological experiments and the circuit map.

To identify possible molecular pathways, we analysed single-cell transcriptome data for cPRC, INNOS and INRGWa identified by spatial mapping (Achim et al., 2015) and unique marker genes (Williams et al., 2017). In cPRCs, we found transporters and synthetic enzymes indicating cholinergic, glutaminergic, glycinergic, GABAergic and adrenergic neurotransmission (Figure 6B). For each neurotransmitter, we also found receptor-encoding genes expressed in INNOS and INRGWa (Figure 6B). The two types of interneurons often expressed different subunits or types of these receptors, indicating differential signalling (Figure 6B).

Based on these data, our experimental results and the mathematical model, we assembled a minimal phototransduction and circuit diagram. The components for which experimental data are available are shown in bold. Other potential molecular players are also indicated (Figure 6C).

To further explore the model, we carried out a global sensitivity analysis. This analysis measures how the output of the model depends on parameter values (Tables 1–5). We also analysed structural identifiability and observability in the model and found most parameters to be identifiable (more details in the Methods section).

**Table 1:**
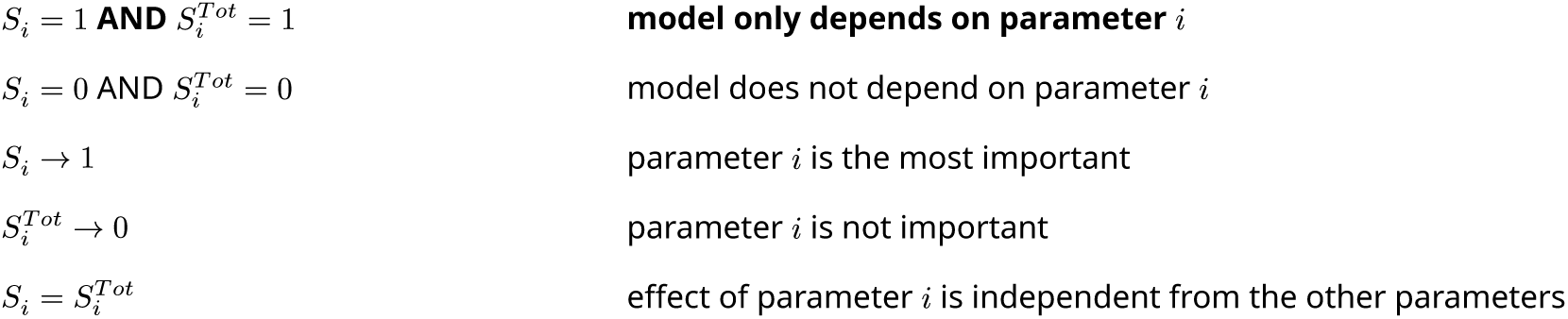
Summary of the significance of global sensitivity indexes.

**Table 2:** Results of the global sensitivity analysis of the model consisting of *C_p_*(*t*), *B*(*t*) and *S*(*t*) variables; fitting to *NOS*-knockout recordings.

|  |  |  |  |  |  |  |  |
| --- | --- | --- | --- | --- | --- | --- | --- |
| 1292 | | $\delta_{C_P}$ | $\kappa_{UV}$ | $\kappa_S$ | $\delta_B$ | $B_0$ | $\delta_S$ |
| 1293 | $S_i$ | 0.0007 | 0.0648 | 0.08 | 0.0127 | 0.3172 | 0.0106 |
| 1294 | ± CI of $S_i$ | <0.0001 | 0.0003 | 0.0004 | 0.0003 | 0.0001 | 0.0016 |
| 1295 | $S_i^{Tot}$ | 0.0277 | 0.2692 | 0.2418 | 0.2295 | 0.6734 | 0.1721 |
| 1296 | ± CI of $S_i^{Tot}$ | 0.0057 | 0.0042 | 0.0044 | 0.0044 | 0.0019 | 0.0048 |

**Table 3:**
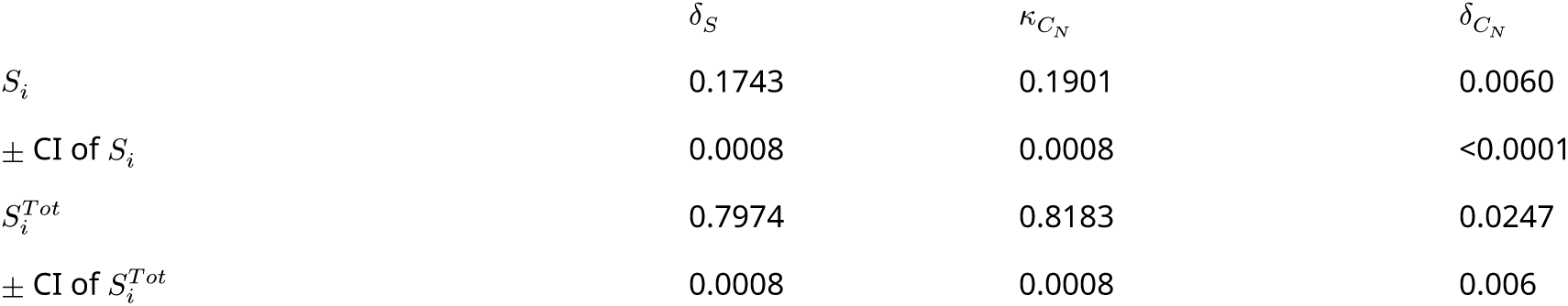
Results of the global sensitivity analysis of the model consisting of *S*(*t*) and *C_N_* (*t*) variables; fitting to WT and *NOS*-knockout INNOS Ca^2+^ recordings.

|  |  |  |  |  |
| --- | --- | --- | --- | --- |
| 1299 | | $\delta_S$ | $\kappa_{C_N}$ | $\delta_{C_N}$ |
| 1300 | $S_i$ | 0.1743 | 0.1901 | 0.0060 |
| 1301 | ± CI of $S_i$ | 0.0008 | 0.0008 | <0.0001 |
| 1302 | $S_i^{Tot}$ | 0.7974 | 0.8183 | 0.0247 |
| 1303 | ± CI of $S_i^{Tot}$ | 0.0008 | 0.0008 | 0.006 |

**Table 4:** Results of the global sensitivity analysis of the model consisting of *S*(*t*), *C_N_* (*t*) and *N*(*t*) variables; fitting to WT NO recordings.

|  |  |  |  |  |  |  |  |
| --- | --- | --- | --- | --- | --- | --- | --- |
| 1306 | | $\delta_S$ | $\kappa_{C_N}$ | $\delta_{C_N}$ | $\kappa_{N,1}$ | $\kappa_{N,2}$ | $\delta_N$ |
| 1307 | $S_i$ | 0.0012 | 0.0056 | 0.0025 | 0.0712 | 0.0059 | 0.5146 |
| 1308 | ± CI of $S_i$ | <0.0001 | <0.0001 | <0.0001 | 0.0004 | <0.0001 | 0.0037 |
| 1309 | $S_i^{Tot}$ | 0.0097 | 0.0506 | 0.0192 | 0.4083 | 0.0401 | 0.8950 |
| 1310 | ± CI of $S_i^{Tot}$ | 0.0081 | 0.0077 | 0.0080 | 0.0043 | 0.0078 | 0.0006 |

**Table 5:**
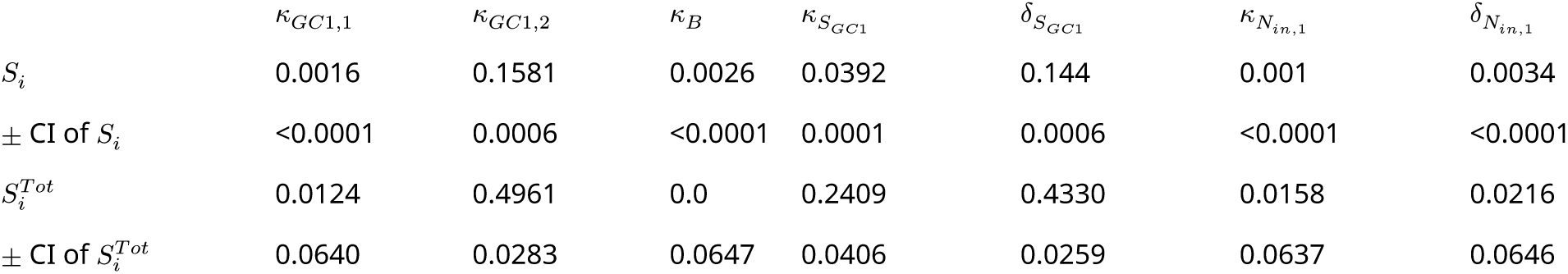
Results of the global sensitivity analysis of the full WT model (remaining parameters); fitting to WT recordings.

As additional validation of the wild-type model, we recorded Ca^2+^-dynamics from the cPRCs in wild-type larvae subject to UV stimulation with different duration. The model captures the time-invariant appearance of the delayed Ca^2+^ peak as observed in our experiments (Figure 6 – figure supplement 7).

The model also allowed us to systematically vary stimulus conditions that would not have been feasible in single-larva experiments. We simulated Ca^2+^ traces under stimulation with two UV pulses (Figure 6 – figure supplement 8 and Materials and Methods) revealing a dependence on inter-stimulus interval. Overall, the model suggests an integratory function of NO signalling to tune circuit output to the strength of UV exposure.

## Discussion

Our work revealed an essential role for NO-mediated signalling in driving UV/violet avoidance behaviour in larval *Platynereis*. NO, produced by postsynaptic INNOS interneurons, signals retrogradely to presynaptic cPRCs via NIT-GC1 leading to delayed and sustained cPRC activation. This late-phase activation of the photoreceptors drives circuit output through projection interneurons and ciliomotor neurons. In the cPRC circuit, synaptic connectivity alone is thus not sufficient to account for circuit dynamics and behavioural change, similar to other fully mapped circuits (Bargmann and Marder, 2013; Imambocus et al., 2022).

### Localised NO signalling in the neurosecretory plexus

NO is a free radical with a millisecond-to-second half life and thus a limited signalling range. In neuronal signalling, NOS is often localised to neurites (Kuntz et al., 2017) close to sGC at synapses (Burette et al., 2002; Garthwaite, 2015). In the cPRC circuit, NOS is localised to the dendritic compartment of INNOS cells where also cPRC postsynaptic sites occur. NIT-GC1, the target of NO signalling is localised to cPRC projections. NO-mediated retrograde signalling thus likely occurs in the neurosecretory plexus where NOS- and NIT-GC1-containing projections are in close proximity and where we detected NO production after UV stimulation. In contrast, INNOS to INRGWa synapses occur outside the neurosecretory plexus in the ventral brain neuropil.

Compartmentalised signalling also occurs in peptidergic modulatory systems and can enable selective network activity during specific behaviours. In the UV-avoidance circuit of *Drosophila* larvae, a peptidergic hub neuron Dp7 links UV-sensory neurons (v’td2) and motor circuits. Dp7 expresses an insulin-like peptide Ilp7 that is required for acute and sustained UV avoidance. Ilp7 signalling occurs at a functionally and morphologically distinct dendritic compartment of Dp7, segregated from other sensory-motor pathways involving Dp7 (Imambocus et al., 2022).

### Functional diversity of NIT-GCs

NO signalling is commonly mediated by sGCs. We identified 12 sGCs in *Platynereis*, but none of these is expressed in the cPRC based on the available scRNAseq data. Instead, we identified an unconventional cPRC-expressed NIT-domain containing GC, NIT-GC1 as the mediator of NO retrograde signalling. In an in vitro assay, we could show that NIT-GC1 can produce cGMP following the addition of an NO donor and that this activity requires the NIT domain.

NIT domains were first identified in bacteria and animal NIT-GCs have only recently been reported (Moroz et al., 2020; Shu et al., 2003). Bacterial NIT domains regulate cellular functions in response to changes in extracellular and intracellular nitrate and/or nitrite concentrations (Camargo et al., 2007). NO is readily converted to nitrate and nitrite (Garthwaite, 2015; Möller et al., 2019; Santos et al., 2011) and these molecules accumulate in placozoans and cnidarians in cells and tissues with high NOS activity (Moroz et al., 2020, 2004). NIT domains in NIT-GCs may also sense nitrate and nitrite, as in bacteria, a possibility we cannot rule out based on our cellular assays with NIT-GC1. If different NIT-GCs have different sensitivities to NO, nitrite and nitrate, then a range of activation timings may be possible due to the different half-lives of these molecules (Lundberg et al., 2011).

In *Platynereis*, we identified 15 NIT-GCs, suggesting a wide range of functions. NIT-GC1 and NIT- GC2 showed specific cellular co-expression but very different subcellular localisation and function. Differences in subcellular localisation and signalling thus seem to also contribute to the diversity of NIT-GC functions in addition to differences in expression.

### Mechanism of phototransduction and neurotransmission in the cPRC circuit

Based on our data herein and previous work we can now propose a more detailed model of cPRC phototransduction and neurotransmission. The cPRCs have high basal Ca^2+^ and respond to UV/violet light dependent on c-opsin1 (Verasztó et al., 2018). c-opsin1 forms a bistable photopigment and signals through Gi/oα and Gβγ (Tsukamoto et al., 2017; Tsukamoto and Kubo, 2023b; Veedin Rajan et al., 2021). In heterologous systems, the Gβγ subunits released following c-opsin1 activation open GIRK channels inducing K+ efflux (hyperpolarisation) (Tsukamoto et al., 2017; Tsukamoto and Kubo, 2023b) and close voltage-gated Ca^2+^ channels, thereby reducing intracellular Ca^2+^ levels (Tsukamoto and Kubo, 2023b). A GIRK channel is also expressed in the cPRCs (Figure 6A). The pathway for the first rapid cPRC-activation phase following UV/violet stimulus is not known but may involve the activation of a CNGα channel expressed in the cPRCs (Tosches et al., 2014). For the second inhibitory phase of phototransduction, we identified a key requirement for ciliary-localised NIT-GC2. The mechanisms may involve c-opsin1-dependent inhibition of tonic NIT-GC2 activity and the reduction of ciliary cGMP. Clarifying how NIT-GC2 signalling interacts with G-protein and GIRK signalling will require further work.

NIT-GC2-dependent signalling is required for the feedforward activation of the INNOS cells through an unknown transmitter. The activation of INNOS leads to NOS activation and NO release, potentially through a canonical Ca^2+^-calmodulin pathway. Our model predicts feedforward inhibition from INNOS to INRGWa, possibly mediated by glycine transmission. NO released by INNOS neurites activates NIT-GC1 in cPRC projections that could lead to cGMP production and the opening of CNGAα. This late-phase cPRC activation does not directly depend on the UV/violet signal and can happen post-stimulus. The late-phase cPRC activation leads to INRGWa activation via an unknown transmitter that is likely different from the one acting on INNOS.

In addition, the cPRC circuit expresses several neuropeptides and their receptors, suggesting further neuromodulatory signals. For example, the INRGWa cells express the proneuropeptide RGWamide and cPRCs and INNOS express its receptor, suggesting retrograde peptidergic signalling in the circuit.

We have recently shown that the cPRC circuit also mediates responses to hydrostatic pressure via the same motor system involving Ser-h1 and the prototroch ciliary band. Increased pressure leads to increased ciliary beating in the prototroch and upward swimming. The effect of pressure on cilia requires synaptic transmission by the serotonergic Ser-h1 neurons (Bezares- Calderón et al., 2023).

Pressure-induced Ca^2+^ transients in cPRCs lack an inhibitory phase and late activation and resemble UV responses in *NIT-GC2*-morphants (Bezares-Calderón et al., 2023). This indicates that the differentiation of sensory cues by the multisensory cPRCs occurs already at the level of sensory signal transduction. The different signalling pathways then likely result in the differential release of transmitters and modulators that are decoded by the postsynaptic interneuron circuit. The behavioural output is opposite, with UV inducing a diving response, while increased pressure inducing upward swimming. The complex activation dynamics and transmitter phenotype of cPRCs could underlie this differential signal processing leading to opposing circuit outputs.

### Nitric oxide confers short-term memory to circuit activity

Retrograde signalling by NO from INNOS to cPRC leads to the sustained activation of cPRCs and postsynaptic neurons even after the end of stimulation. This late-phase activation occurs at a fixed time after stimulus, independent of stimulus duration. The high-Ca^2+^-state is then maintained for several tens of second (Figure 4A, B). NO signalling thus induces a transient and time-invariant peak or short-term memory trace in the *Platynereis* larval brain. Because of the short life time of NO, this molecule may be well suited to encode transient memory traces (Kuntz et al., 2017).

In the ellipsoid body of the *Drosophila* central brain, NO signalling has a similar function. Here, NOS is specifically expressed in the R3 ring neurons and is required for the short-term (∼4 sec) visual memory of objects (Kuntz et al., 2017). NO is produced in the axons of R3 neurons and acts directly on sGC in the same axons. This autocrine signal leads to a CNG-dependent temporary increase in Ca^2+^ levels, carrying the working memory trace (Kuntz et al., 2017).

In the *Platynereis* circuit, our mathematical model indicates that the magnitude of the NO- dependent signal depends on the intensity and duration of the UV/violet stimulus. This suggests that the NO-dependent memory trace also encodes the intensity and duration of the stimulus.

During UV avoidance in the planarian *Schmidtea mediterranea*, neuropeptide signalling has a similar integratory function. Planarians exposed to UV light for >30 sec remain active for extended periods (several minutes) post-stimulation (Bray et al., 2023). If neuropeptide signalling is defective, animals can still respond to UV light, but do not maintain a latent memory state and do not display increased post-stimulus activity.

### The possible mechanism of UV-induced downward swimming

Non-directional UV/violet light induces downward swimming head down in *Platynereis* larvae (Verasztó et al., 2018). Since stimulus direction is not relevant, swimming direction must be determined by gravity.

Connectome reconstruction and whole-body cell annotation in the 3-day-old *Platynereis* larva has not revealed any balancer organ to sense orientation in the gravity field (Bezares-Calderón et al., 2019; Verasztó et al., 2020). Body orientation may thus be determined by physical parameters including the buoyancy, centre of mass and shape of the larva as well as differential ciliary activity. Here we only focused on the prototroch ciliary band and more work will be needed to unravel the process of graviorientation. For example, quantifying ciliary activity simultaneously in the prototorch and the telotroch (and also the paratrochs in nectochaete- stage larvae) during UV and pressure stimulation may reveal differential responses across ciliary bands.

### Future directions

The mathematical model could be further developed by more explicitly modelling cPRC phototransduction and synaptic transmission. For example, cGMP levels could also be directly measured in cPRCs by Green-cGull and built into the model. The contributions of the individual neurotransmitters in the circuit could also be analysed by knockdown/knockout experiments. The addition of an NO donor to larvae combined with calcium imaging could more directly test the relationship between NO levels and cPRC responses.

Beyond these details of cellular signalling, the biggest question remains the mechanism of graviorientation in the larvae. We do not understand what leads to differential behavioural output during the pressure on-response (upward swimming) (Bezares-Calderón et al., 2023) and the UV response (downward swimming). Under both conditions, the prototroch cilia increase their beating. The only difference we detected so far is in the dynamics of cPRC Ca^2+^ responses.

Identifying the neuronal and biophysical underpinnings of these opposite behavioural outcomes will require further genetic and biophysical experiments combined with mathematical modelling.

### UV-avoidance circuits of extraocular photoreceptors

Animals evolved distinct photosensory systems coupled to non-overlapping circuits and guiding unique behavioural responses. These sensory systems can employ different opsin molecules and be tuned to different wavelengths of light. The avoidance of noxious UV/violet light is often mediated by extraocular photoreceptors and their circuits, distinct from the pigmented visual eyes. These two types of systems co-exist in *Platynereis* where the pigmented eyes and eyespots guide phototaxis with a maximum sensitivity to cyan light (∼500 nm) (Gühmann et al., 2015; Randel et al., 2014; Verasztó et al., 2018). In free-swimming larvae in a vertical column, cyan light induces phototactic upward-swimming while UV induces diving. The two behaviours compete and the outcome depends on the ratio of UV-to-cyan light (Verasztó et al., 2018).

Planarians also have pigmented visual eyes mediating phototaxis to cyan light and peripheral extraocular photoreceptors mediating UV avoidance behaviour (Shettigar et al., 2021). *Drosophila* larvae have cerebral eyes called Bolwig’s organs involved in phototaxis (Kane et al., 2013) and several types of extraocular UV-sensory cells that tile the body wall and mediate UV avoidance (Imambocus et al., 2022).

One common feature of UV-avoidance circuits is their sustained post-stimulus activation following UV exposure. This can involve peptidergic signals as in planaria and maggots (Bray et al., 2023; Imambocus et al., 2022) or NO as in *Platynereis*. Volume transmission is well suited to integrate light exposure and maintain an internal state following noxious stimulation. The amount of modulator released can scale with stimulus intensity or duration and maintain an altered circuit state due to the slower decay of the diffusive signals relative to synaptic transmission.

Overall, we have revealed how NO shapes the dynamics of a fully mapped sensory-motor UV avoidance circuit. We identified an unconventional GC as the NO receptor and measured circuit activity in different genetic backgrounds. Finally, we could link circuit activity to light-avoidance behaviour. The richly modulated multi-transmitter cPRC circuit will serve as a fertile ground for future studies on how neuromodulators shape activity and behaviour.

## Key resources table

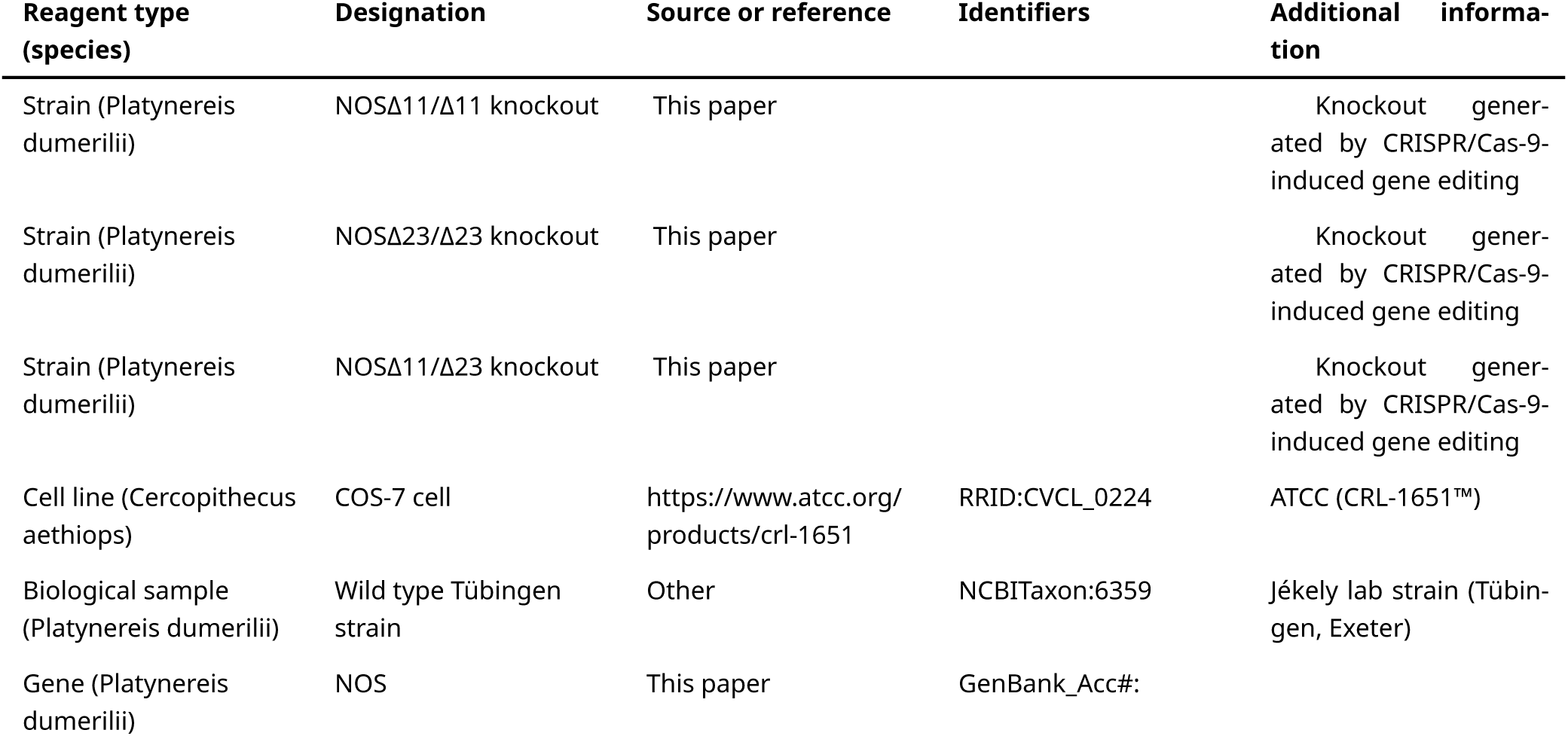
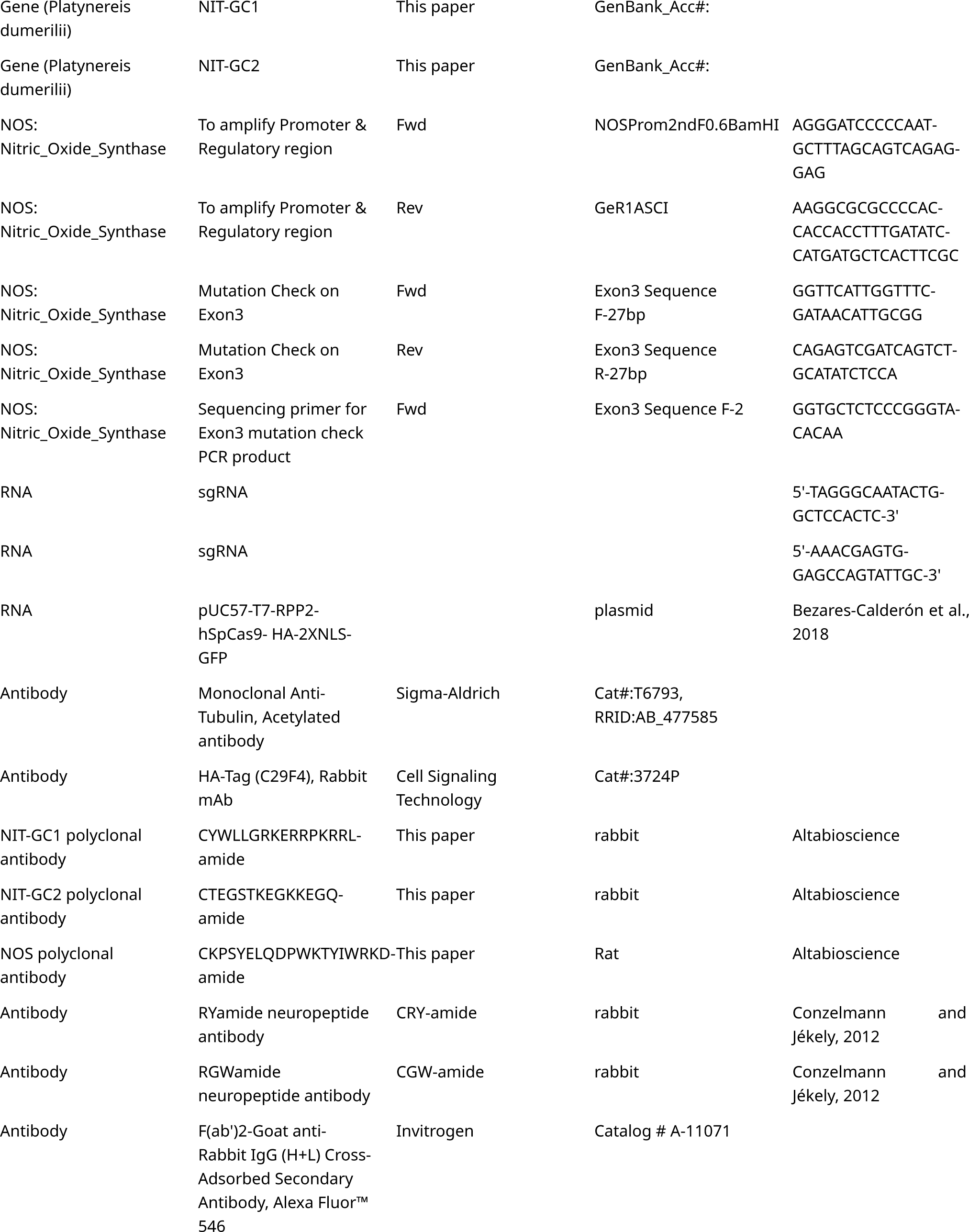
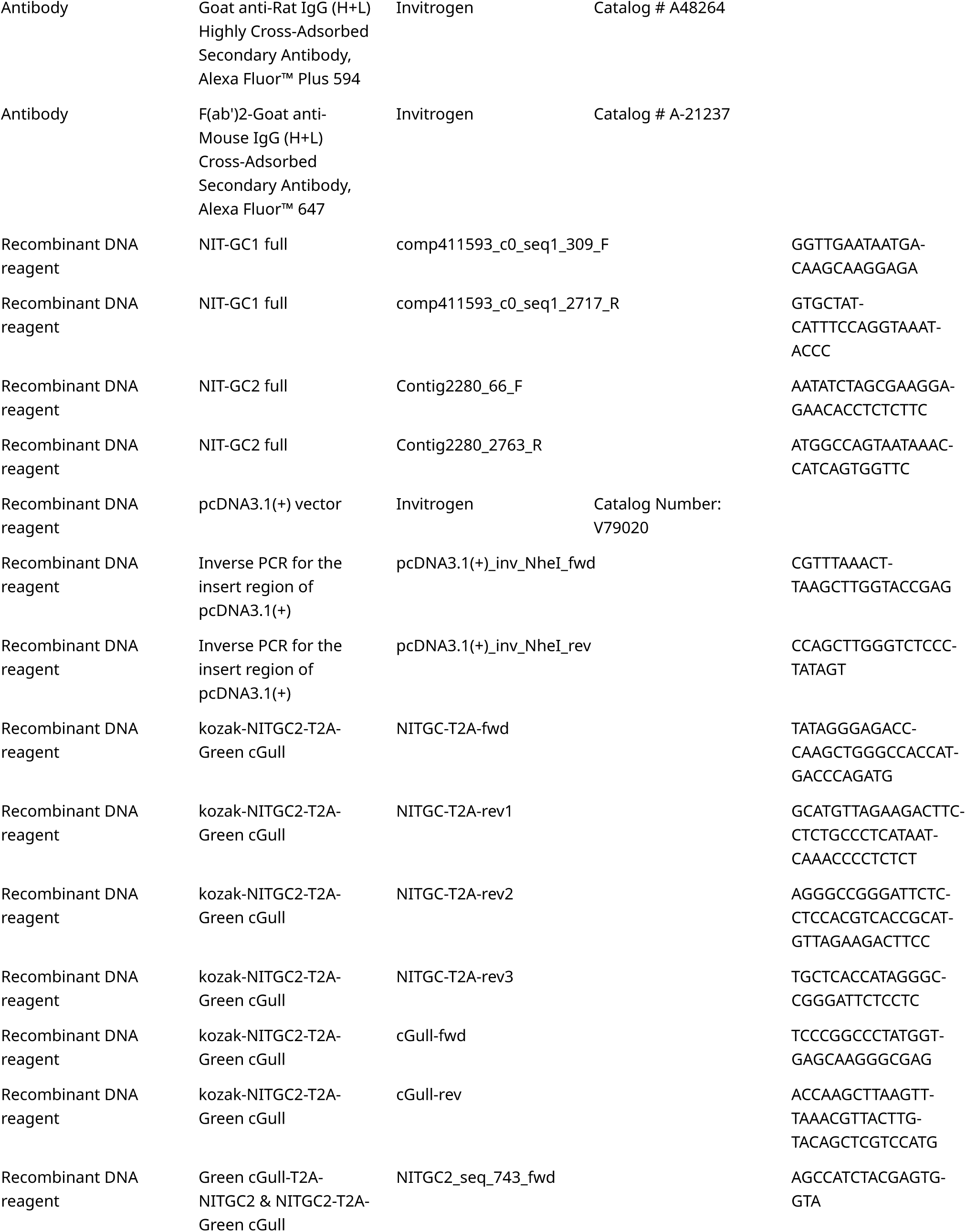
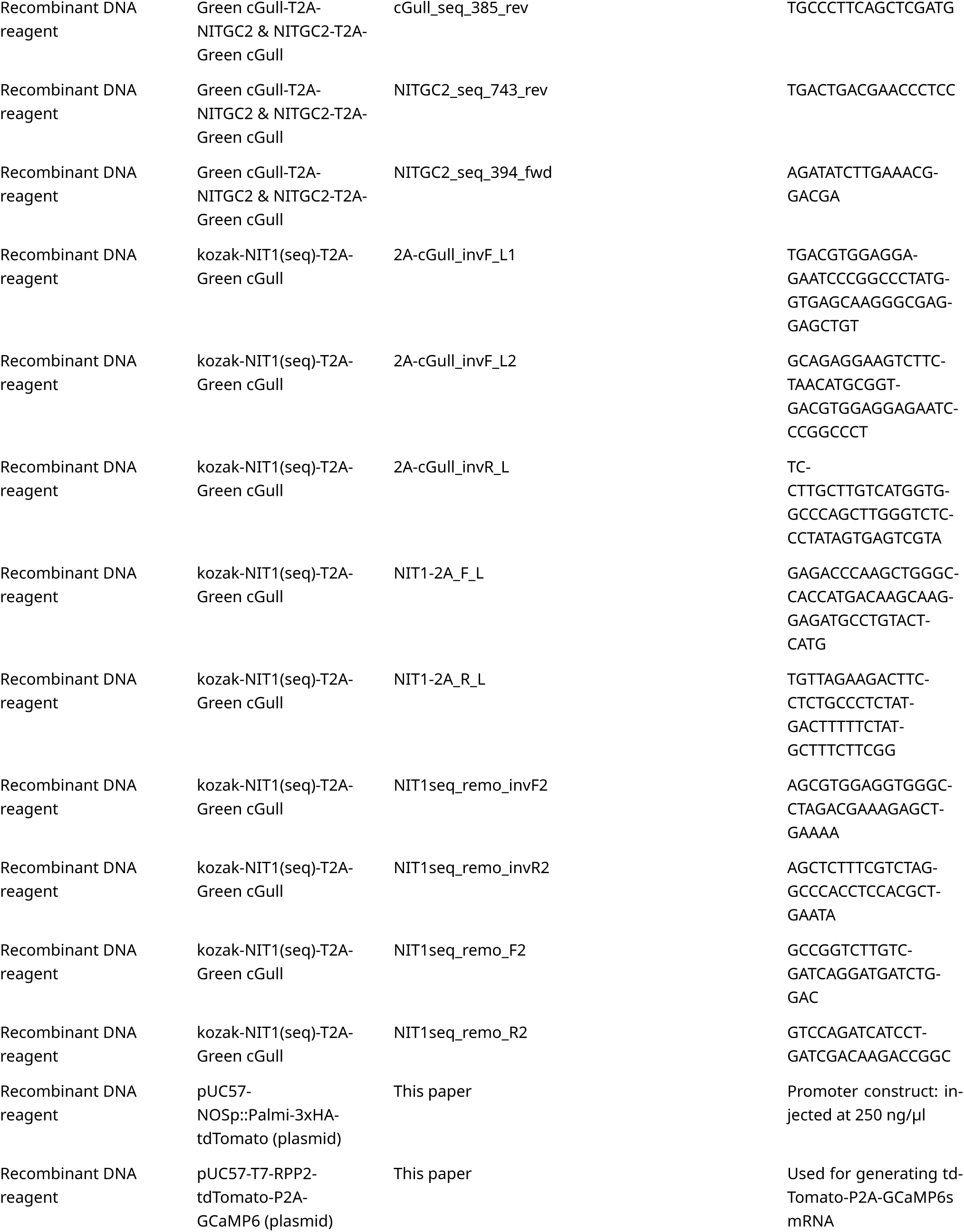
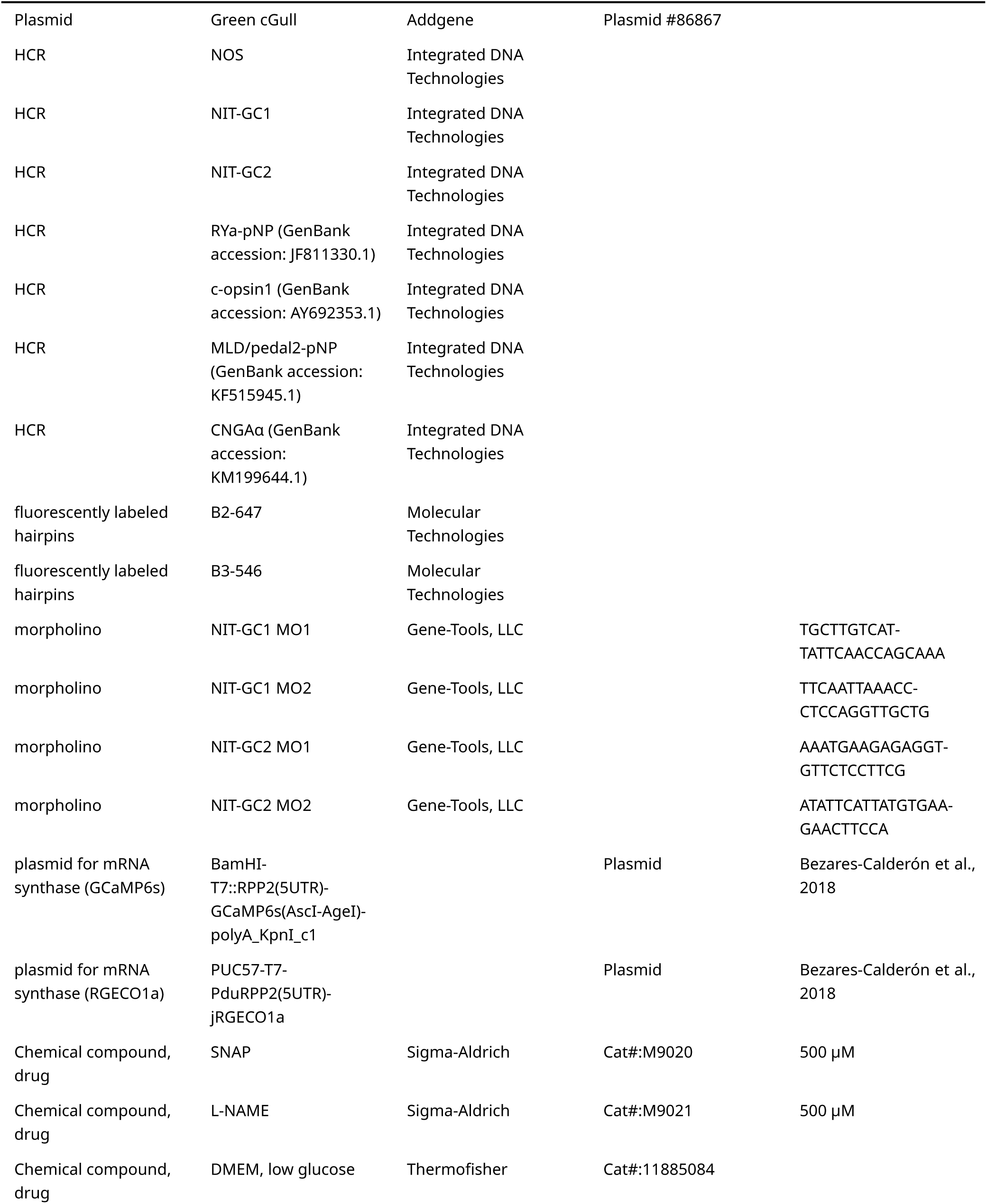
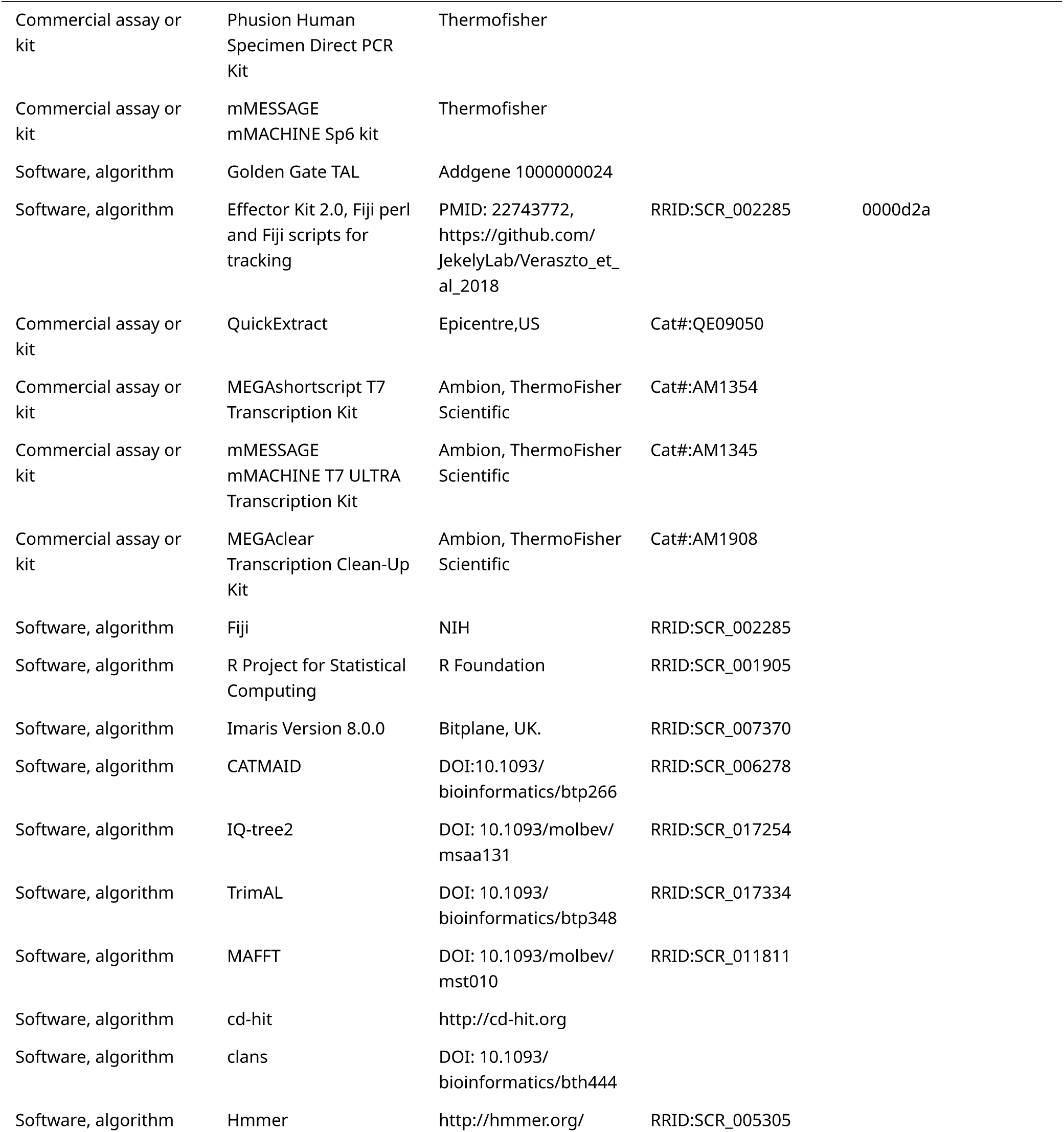

## Materials and Methods

### Animal culture

*Platynereis dumerilii* were cultured in an aquatic laboratory facility following established protocols. Larvae were raised in sterile artificial sea water (fASW, Tropic Marin) at 32 ppm and kept at 18°C in a 16/8-hr light–dark cycle. Worms were fed a mixed diet of microalgae, rotifers and spinach. For a detailed protocol see (Hird et al., 2024).

### CRISPR-Cas9 design and Microinjection

For the generation of a *NOS* knockout, we used an sgRNA targeting the third exon of the *Platynereis dumerilii NOS* gene (target site: 5′-GGGCAATACTGGCTCCACTC-3′). We selected the site to avoid polymorphic sites in our laboratory culture. The sgRNA was assembled from two annealed oligonucleotides (5′-TAGGGCAATACTGGCTCCACTC-3′, 5′- AAACGAGTGGAGCCAGTATTGC-3′) forming overhangs for cloning into the BsaI site of the plasmid pDR27456 @hwang2013, which contains next to the BsaI site a tracrRNA sequence. We used this plasmid to PCR amplify DNA (primers: T7, 5′-AAAAGCACCGACTCGGTGCC-3′) for synthesizing the sgRNA. To purify the PCR product, we used the QIAquick PCR Purification Kit (Qiagen). We then synthesized sgRNA with the MEGAshortscript Kit (Thermo Fisher Scientific) and purified it with the MEGAclear Kit (Thermo Fisher Scientific). To produce the Cas9-mRNA, we used the plasmid (pUC57-T7-RPP2-Cas9) containing the Cas9 ORF fused to 169 base pair 5′ UTR from the *Platynereis dumerilii* 60S acidic ribosomal protein P2. After in vitro transcription with the mMessage mMachine Kit (Thermo Fisher Scientific), we capped and polyA-tailed the Cas9-mRNA with the Poly(A) Tailing Kit (Thermo Fisher Scientific). We coinjected the sgRNA (18 ng/ml) and the Cas9-mRNA (180 ng/µl) into fertilized eggs of *Platynereis dumerilii* wild-type parents according to an established injection procedure (Conzelmann et al., 2013a). Eggs were kept at 18°C for 45 min before injection and were injected at 14.5°C. The injected individuals were kept at 18°C for 5 to 8 days in 6-well-plates (Nunc multidish no. 150239, Thermo Scientific) and then cultured at 22°C until sexual maturity. The mature worms were crossed to wild-type worms and the progeny was genotyped, resulting in two founder lines, which were bred to homozygosity.

### Genotyping of NOS alleles

For genotyping of the *NOS* locus, genomic DNA was isolated from single larvae, groups of 6-20 larvae, or from the tails of adult worms. The DNA was amplified by PCR (primers: 5′- GGTTCATTGGTTTCGATAACATTGCGG-3′, 5′-CAGAGTCGATCAGTCTGCATATCTCCA-3′) with the dilution protocol of the Phusion Human Specimen Direct PCR Kit (Thermo Scientific). The PCR product was sequenced directly with a nested sequencing primer (5′- GGTGCTCTCCCGGGTACACAA-3′). A mixture of wild-type and deletion alleles in a sample gave double peaks in the sequencing chromatograms, with the relative height of the double peaks reflecting the relative allele ratio in the sample.

### Morpholino-mediated knockdowns

Morpholino injections were performed as previously described (Conzelmann et al., 2013b). We used the following morpholinos to target the NIT-GC1 and NIT-GC2 genes (GeneTools, LLC). NIT-GC1 MO1: TGCTTGTCATTATTCAACCAGCAAA, NIT-GC1 MO2: TTCAATTAAACCCTCCAGGTTGCTG, NIT-GC2 MO1: AAATGAAGAGAGGTGTTCTCCTTCG, NIT-GC2 MO2: ATATTCATTATGTGAAGAACTTCCA.

### Vertical column setup to measure photoresponses

We assayed larval photoresponses in a vertical Plexiglas column (31 mm x 10 mm x 160 mm water height). For stimulation, we illuminated the column from above with a monochromator (Polychrome II, Till Photonics) controlled by AxioVision 4.8.2.0 (Carl Zeiss GmbH, Jena) via analog voltage. The light passed a collimator lens (LAG-65.0-53.0-C with MgF2 Coating, CVI Melles Griot) before entering the column. The column was illuminated from both sides with light-emitting diodes (LEDs). The LEDs on each side were grouped into two strips. One strip contained UV (395 nm) LEDs (SMB1W-395, Roithner Lasertechnik) and the other infrared (810 nm) LEDs (SMB1W-810NR-I, Roithner Lasertechnik). The UV LEDs were run at 4 V. The infrared LEDs were run at 8 V (overvoltage) to illuminate the larvae for imaging with a DMK camera (DMK 22BUC03, The Imaging Source). We recorded videos at 15 frames per second with the software IC Capture (The Imaging Source).

We compared the behavior of wildtype and *NOS*-knockout larvae in the vertical column. In the inhibitor experiments we compared wildtype larvae before and after treatment with 0.1 mM or 1.0 mM NOS inhibitor L-NAME (in seawater). All larvae were three days oldt. Before the experiment, the larvae were mixed and left in the dark for 5 min. The larvae were recorded for 1.5 min in the dark followed by exposure to diffuse UV light (395 nm) from the side for 2 min. Then the larvae were left for another 2 min in darkness followed by exposue to collimated cyan (480 nm) light from above for 2 min, then 2 min darkness, and finally collimated UV (395 nm) light from above for 2 min. Scripts are available at https://github.com/JekelyLab/Jokura_et_al_NOS.

The TrackMate7 plugin (“TrackMate 7,” n.d.) in Fiji ImageJ was used to trace the swimming trajectory of the larvae, using the DoG detector for 3 pixel size and the LAP Tracker, filtered at linking max distance: 10.0 pixels, gap closing max distance: 5.0 pixels.

### Identification and Phylogenetic Analysis of NOS sequences

To identify NOS sequences, we obtained a seed database of oxygenase domains from the Pfam database (PF02898). From these sequences, we generated a Hidden Markov Model (HMM) to mine the genomes and transcriptomes of 47 metazoan, 2 choanoflagellate and 2 filasterea species. The HMM model was run in HMMR3 (Eddy, 2011) with an e-value of 1e−15. We ran CD-Hit (Fu et al., 2012) to eliminate redundant sequences (at a 80% threshold). We aligned the sequences with MAFFT version 7, with the iterative refinement method E-INS-i (Katoh and Toh, 2008). The alignment was trimmed with TrimAl in gappy-out mode (Capella-Gutierrez et al., 2009). To calculate a maximum-likelihood tree, we used IQ-tree2 with the LG+G4 model with the following options: iqtree2 -s “FILENAME” -mset WAG, LG, Blosum62, Dayhoff, JTT, Poisson -B 1000 -alrt 1000 -T AUTO. Branch-support values are based on 1000 replicates with the ultrafast bootstrap approximation (UFBoot) (Hoang et al., 2018) and aLRT-SH-like (Guindon et al., 2010) methods.

### Identification and Phylogenetic Analysis of NIT-GC sequences

To identify NIT-GCs, we obtained a seed database of adenylate and guanylate cyclase catalytic domains from the Pfam database (PF00211). From these sequences, we generated a Hidden Markov Model (HMM) to mine the 45 metazoan, 2 choanoflagellate and 2 filasterea genomes and transcriptomes. The HMM model was run in HMMR3 (Eddy, 2011) with an e-value of 1e−15. We ran CD-Hit (Fu et al., 2012) to eliminate redundant sequences (at a 80% threshold). To identify clusters, we used sequence-similarity-based clustering in Clans (Frickey and Lupas, 2004) and the convex-clustering option with 100 jack-knife replicates. The NIT-GCs are extremely well conserved within the membrane-bound guanylate cyclases and form an easily recognizable cluster. To analyze the phylogeny of NIT-GCs, the cluster containing these GCs together with membrane-bound guanylate cyclases were parsed and used for tree building. We aligned the sequences with MAFFT version 7, with the iterative refinement method E-INS-i (Katoh and Toh, 2008). The alignment was trimmed with TrimAl in gappy-out mode (Capella-Gutierrez et al., 2009). To calculate a maximum-likelihood tree, we used IQ-tree2 with the LG+G4 model with the following options: iqtree2 -s “FILENAME” -mset WAG, LG, Blosum62, Dayhoff, JTT, Poisson -B 1000 -alrt 1000 -T AUTO. Branch-support values are based on 1000 replicates with the ultrafast bootstrap approximation (UFBoot) (Hoang et al., 2018) and aLRT-SH-like (Guindon et al., 2010) methods.

### Single-cell analysis

We used a single-cell RNA sequencing dataset from *Platynereis* larvae (Achim et al., 2015) that was further filtered to 107 cells (Williams et al., 2017). We normalized the raw read-count data to TPM and converted to log10. To calculate % expression, we used the sum of the expression levels in the 107 cells for each gene. The total TPM of each gene between the samples was used to calculate the percentage of expressed genes. We processed the data in Python and plotted in R. Individual cells were identified by suits of well-characterised marker genes and spatial mapping (Achim et al., 2015).

### In situ HCR

Larvae were fixed and treated with Proteinase K, according to the conventional WMISH protocol (Tessmar-Raible et al., 2005), with fixation in 4% paraformaldehyde/ PTW (PBS with 0.05% Tween20) for 2 hr at room temperature, and Proteinase K treatment in 100 µg/ml Proteinase K/PTW for 3 min (Tessmar-Raible et al., 2005). Specifically, for the HCR protocol, samples were processed in 1.5 ml tubes. Probe hybridization buffer, probe wash buffer, amplification buffer, and fluorescent HCR hairpins were purchased from Molecular Instruments (Los Angeles, USA). The hairpins associated with the b2 initiator sequence were labeled with Alexa Fluor 647, and the hairpins associated with the b3 initiator sequence were labeled with Alexa Fluor 546. To design probes for HCR, we used custom software (Kuehn et al., 2021) to create 20 DNA oligo probe pairs specific to *P. dumerilii NOS*, *NIT-GC1*, *NIT-GC2*, *RYa proneuropeptide* (GenBank accession: JF811330.1), and *MLD/pedal peptide 2 proneuropeptide* (GenBank accession: KF515945.1). The *NOS*, *NIT-GC1* and *NIT-GC2* probes were designed to be associated with the b2 initiator sequence, while the *RYa* and *MLD/pedal peptide 2* probes were designed to be associated with the b3 initiator sequence. For the detection stage, samples were pre-hybridized in 200 µl of probe hybridization buffer for 1 hr at 37°C, and then incubated in 250 µl hybridization buffer containing probe oligos (4 pmol/ml) overnight at 37°C. To remove excess probe, samples were washed 4× with 1 ml hybridization wash buffer for 15 min at 37°C, and subsequently 2× in 1 ml 5× SSCT (5× SSC with 0.1% Tween20) for 5 min at room temperature. For the amplification stage, samples were pre-incubated with 100 µl of amplification buffer for 30 min, room temperature, and then incubated with 150 µl amplification buffer containing fluorescently labelled hairpins (40nM concentration (2ul of 3uM stock in 150ul amplification buffer, snap-cooled as described; (Choi et al., 2018)) overnight in the dark at 25°C. To remove excess hairpins, samples were washed in 1 ml 5× SSCT at room temperature, twice for 5 min, twice for 30 min, and once for 5 min. During the first 30 min wash, samples were counterstained with DAPI (Cat. #40043, Biotium, USA).

### Immunohistochemistry

Whole-mount immunostaining of *Platynereis* larvae fixed with 4% paraformaldehyde was carried out using the primary antibodies listed in the KEY RESOURCES TABLE. Immunostainings were carried out as previously described (Conzelmann and Jékely, 2012).

### Antibody generation and purification

Antibodies were raised in rats or rabbits against synthetic peptides containing an N-term Cys residue (Altabioscience). The same Cys-containing synthetic peptides were used for affinity purification. Antibodies were affinity purified from sera on a SulfoLink Coupling Resin (Thermo Fischer) as previously described (Conzelmann and Jékely, 2012). View a detailed protocol here: https://bio-protocol.org/exchange/preprintdetail?id=2333&type=3

### Transient transgenesis

A 12 kb genomic region upstream of the start site of the *NOS* gene was amplified and cloned upstream of 3xHA-Palmi-tdTomato to yield the plasmid *NOS-3xHA-Palmi-tdTomato*. Larvae injected with the promoter construct (ca. 250 ng/ml) were analysed for reporter expression at three days post fertilization on an AxioImager Z.1 fluorescence microscope (Carl Zeiss GmbH, Jena) and if positive were fixed for immunostaining. The immunostaining for the HA-tagged reporter was done as described (Verasztó et al., 2017). Stained specimens were imaged on an LSM 780 NLO or LSM 880 Airysan Confocal Microscope (Carl Zeiss GmbH, Jena).

### Calcium imaging

For Ca^2+^ imaging, GCaMP6s mRNA (1 mg/ml) was injected into zygotes as described previously (Randel et al., 2014). At 49-55 hours post fertilisation, larvae were immobilised in 2.5% agarose in artificial sea water and mounted between a slide and coverslip spaced with adhesive tape. Larvae were imaged at room temperature on a Zeiss LSM 880 with Airyscan (with a C- Apochromat 63X/1.2 Corr - water) with a frame rate of 1.88 frame/sec and an image size of 512 x 512 pixels. The larvae were stimulated in a region of interest (a circle with 50 pixel diameter) with a 405 nm laser controlled by the bleaching mode of the Zeiss software. The imaging laser had a similar intensity than the stimulus laser but covered an area that was 10 times larger than the stimulus ROI.

### Ciliary beat-frequency analysis

Larvae were imaged at room temperature on a Zeiss AxioObserver Z1 Microscope (Carl Zeiss Microscopy, Germany) with a C-Apochromat 10X lens, 850 nm power LED (OSRAM OSLON® Black, SFH 4716S) and filter sets equipped with a 405/40 ET excitation Bandpass, Beamsplitter dichroic mirror 495LPXR and without an emission filter with a IR camera (UI-3360CP-NIR-GL Rev.2, IDS Imaging Development Systems GmbH, Germany) at a frame rate of 200 frame/sec with 0.2 msec pulse controlled by the LED stroboscopic illumination system synchronized with the exposure signals from the camera (TYM-002, developed by Dr.Shoji A. Baba, Ochanomizu University, Otsuka, Japan) and an image size of 640 x 480 pixels. The larvae were stimulated with a 395 nm LED (M395L5, Thorlabs). Kymographs of the cilia obtained with Fiji were used to measure the ciliary beat frequency.

### cGMP-production assay

To measure cGMP production in a cell culture assay we used the Green cGull reporter (Matsuda et al., 2016). Full-length *NIT-GC1* and *NIT-GC2* coding sequences were amplified by PCR from a *Platynereis dumerilii* cDNA library and cloned into the pcDNA3.1(+) vector with the T2A self- cleaving sequence. For the assay, COS-7 cells (Angio-proteomie, CAT no. cAP-0203) with a low endogenous level of soluble guanylate cyclase activity were used. Cells were not contaminated with mycoplasma. Cells were maintained at 37 °C in 35 mm dishes (Nunc™ Glass Bottom Dishes) containing 3 ml of DMEM, low-glucose medium (Thermo; Cat. No. 11885084) supplemented with 10% fetal bovine serum (Thermo; Cat. No. 10082147). Upon reaching approximately 85% confluency, cells were transfected with the pcDNA3.1(+)-Green-cGull-T2A-NITGC1 plasmid. Transfections were carried out with 150 ng of each plasmid and 0.3 μl of Lipofectamine 3000 Reagent (invitrogen; Cat. No. L3000001). Three hours post-transfection, the culture medium was substituted with fresh DMEM medium. For single-wavelength imaging experiments, cells in 35- mm dishes were washed twice and imaged in modified Ringer’s buffer (140 mM NaCl, 3.5 mM KCl, 0.5 mM NaH2PO4, 0.5 mM S-3 MgSO4, 1.5 mM CaCl2, 10 mM HEPES, 2 mM NaHCO3 and 5 mM glucose). The dishes were mounted on a stage heated to 37 °C and imaged on an inverted confocal microscope (LSM880, Carl Zeiss GmbH, Jena) equipped with an oil-immersion objective lens (UApo/340, 40x, NA = 0.17). The exposure time of the EM-CCD camera was controlled by the ZEN software (Carl Zeiss GmbH, Jena). Images were acquired every 15 s for 10 min and stimulation was initiated 2 min after starting image acquisition. Imaging data were processed with ImageJ (National Institutes of Health, Bethesda, MD, USA). As NO donor, we used S-Nitroso- N-acetyl-D, L-penicillamine (SNAP, Sigma-Aldrich, St.

### Mathematical modelling

#### Wild type model

The mathematical model for the wild type data comprises 11 dynamical equations:

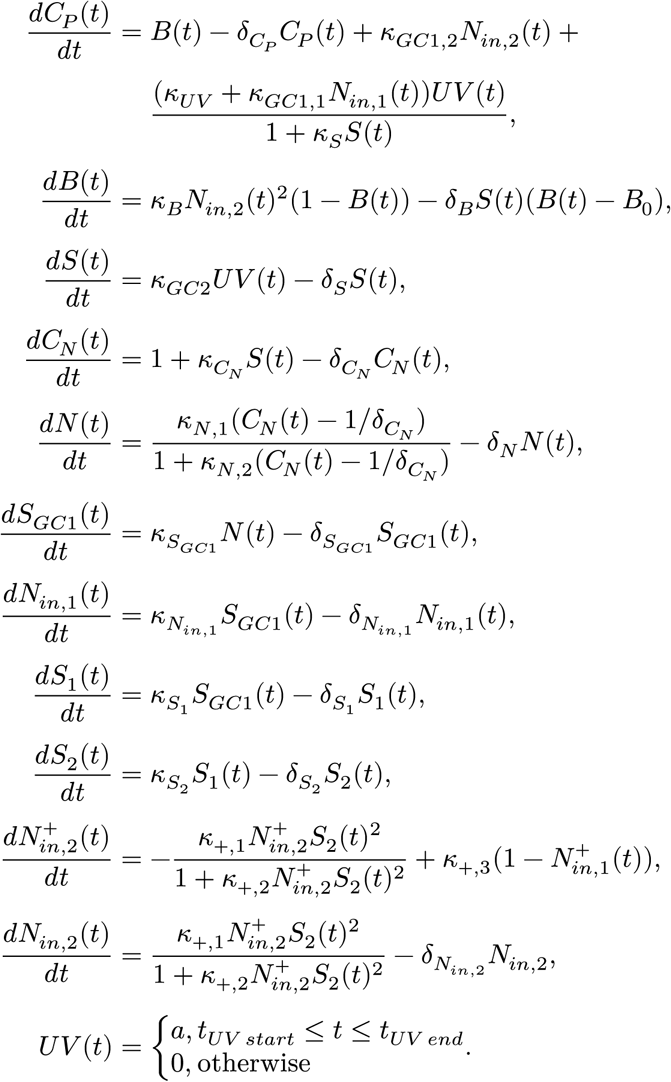

The *C_P_* (*t*) equation describes the dynamics of the Ca^2+^ concentration in the cPRC cells, the *B(t)* variable captures changes in the baseline level of the Ca^2+^, while the *κ*_*GC*1,1_*N_in_*_,2_(*t*) term captures the Ca^2+^ increase due to the late NO signalling. Finally, the term (*κ_UV_* + *κ*_*GC*1,2_*N_in_*_,1_(*t*))*UV* (*t*)/(1 + *κ_S_S*(*t*)) captures the UV-dependent dynamics. This dynamics has three components: direct UV-dependent phototransduction with rate *κ_UV_* ; inhibition from NIT-GC2 and UV-dependent phototransduction, captured by 1/(1 + *κ_S_S*(*t*)); and NO signalling dependent activation, given by *κ*_*GC*1,2_*N*_*in*,1_(*t*).

The *B*(*t*) equation is added to capture the decrease of the Ca^2+^ levels observed during the UV stimulation through the term −*δ_B_S*(*t*)(*B*(*t*) − *B*_0_); the mechanism of this decrease is unknown. It also describes the observed increase of the baseline Ca^2+^ levels due to late NO signalling through *κ_B_N_in_*_,2_(*t*)^2^(1 − *B*(*t*)); the mechanism of this increase is also unknown.

The *S*(*t*) equation describes effects of assumed NIT-GC2 and UV-dependent phototransduction on Ca^2+^ levels. The *S*(*t*) variable has two roles. It inhibits the UV dynamics and also represents synaptic coupling from the cPRC cells, which is assumed to act in an excitatory fashion on INNOS cells. Both of these processes are assumed to be suppressed in *NIT-GC2*-morphants. To this end, we introduce a parameter, *κ*_*GC*2_, which is kept fixed at *κ*_*GC*2_ = 1 in the *NOS*-knockout and WT models.

The *C_N_* (*t*) equation describes the Ca^2+^ dynamics in the INNOS cells. The 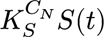 term describes the Ca^2+^ increase due to the synaptic signal *S*(*t*). The *N*(*t*) variable represents NO produced and secreted by the INNOS cells. The 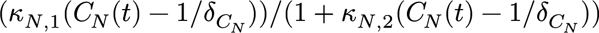 term describes the Ca^2+^-dependent NO production within the INNOS cells.

The *S_GC_*_1_(*t*) equation describes the NO signalling dependent on NIT-GC1. This signal activates two pathways. The first modulates phototransduction via the *N_in_*_,1_ variable. The second pathway captures the delayed time-limited increase in the Ca^2+^ in cPRC cells. The delay occurs because the signal *S_GC_*_1_(*t*) is transmitted via the *S*_1_(*t*) and *S*_2_(*t*) equations. To achieve time- bounded response to the NO signal, we introduce the 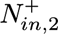 and *N_in_*_,2_ equations. The 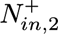 equation describes production of *N_in_*_,2_. The *N_in_*_,2_ equation captures the release of the *N_in_*_,2_ variable. The *N_in_*_,2_ variable increases the Ca^2+^ levels in the cPRC cells in two ways, one through a direct mechanism, the other by increasing the baseline level *B*(*t*).

Finally, the UV stimulation is described by a square pulse with amplitude *a* and duration *t_UV end_* − *t_UV start_*. The overall model structure is informed by the data collected in experiments with 20 seconds as well as with longer UV stimulation. Model parameters were fitted using exclusively experiments with 20 seconds UV stimulation. Experiments with longer UV stimulation were used to validate the estimated parameter values.

We do not model the dynamics of Ca^2+^ in the INRGWa cells, which depends on the other equations in a feed-forward manner. Phenomenologically, we observe increase in Ca^2+^ levels in the INRGWa cells that is correlated with increases of the Ca^2+^ levels in the cPRC cells; term *A*(*C_P_*, *t*) in Figure 6A. We also see an inhibitory effect correlated with the Ca^2+^ levels in the INNOS cells; term *I*(*C_P_*, *t*) in Figure 6A. However, for the following three reasons, we choose not model the INRGWa cells. First, the INRGWa cells were not a subject of direct experimental modifications. Second, the quality of the recordings from the INRGWa cells is lower than that of the other cells, for example, the wild-type recordings do not appear to be at a steady state before the UV stimulation. Additionally, recordings in the *NIT-GC2*-morphants are the only dataset that shows a large difference between mean and median time-series, which we take as a measure of robustness of response. Finally, the data collected in the experiments do not provide enough information to specify how the duration of the UV stimulation affects the dynamics of Ca^2+^ in the INRGWa cells.

#### *NOS*-knockout model

The *NOS*-knockout model is derived from the WT model by eliminating the *N*(*t*) variable, which represents the NO produced by the INNOS cells in a NOS-dependent manner, and terms that depend on it. This results in the system:

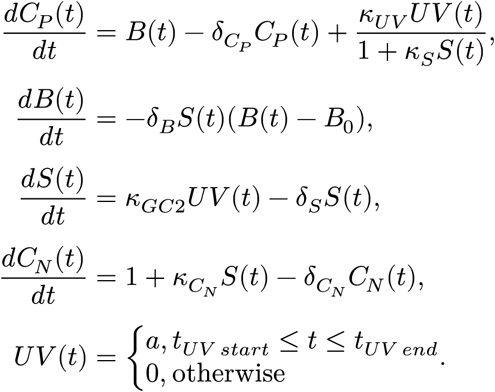

We use the same system of equations to describe the *NIT-GC2*-morphants. To this end, we lower the value of *κ_GC_*_2_ to 0.15; this value captures an average effectiveness of NIT-GC2-morpholino knockdown of 85%. Furthermore, in the NIT-GC2 morpholino condition, we do not fit the INNOS data since we assumed that the feedforward coupling between the cPRC cells and INNOS cell depends on NIT-GC2 activity (that is, we assume that the Ca^2+^ levels in INNOS cells are constant in this condition); this is on average consistent with the collected data.

#### Parameter fitting (genetic algorithm)

To find sets of model parameters that reproduce recordings from the Ca^2+^ imaging experiments, we define an optimisation problem to minimise the discrepancy between the modelled Ca^2+^ traces and the experimental data. To obtain the modelled Ca^2+^ traces, we analytically calculate the steady state in the absence of UV stimulation (in which many terms in the model equations vanish), and then use the explicit integration algorithm ode45 in Matlab to simulate the model with the computed steady state as the initial condition with UV stimulation reinstated.

We quantify the difference between the model output and the data using a vector-valued objective function. The first component of this function measures this difference during the UV stimulation, whilst the second component measures the difference from the end of the stimulation until the end of the recording. We thus define a vector-valued objective function, 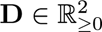, as:

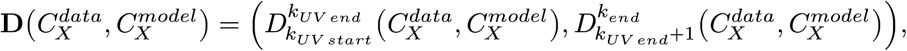

with

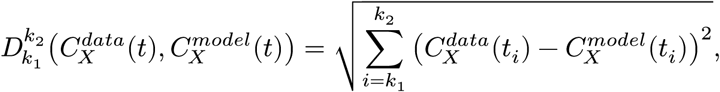

where the timepoints *t_i_* = *i*Δ*t* are equispaced with Δ = 0.5*s* and where *t_k_UV start__* = *t_UV start_* (and similar for the other time points). Here 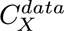 denotes normalised experimental data and 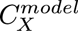 denotes normalised simulated Ca^2+^ data with *X* ∈ {*P*, *N*, *R*} indicating cell type. Simulated Ca^2+^ data are normalised in the same way as the experimental data, i.e., we divide all values in the data uniformly by the value at *t*_*k*_*UV start*_−1_; a time point immediately prior to the start of the UV stimulation. (Recall that *t_UV start_* is the time of the start of UV stimulation, *t_UV end_* is the time of the end of UV stimulation, and *t_end_* is the last time sample.) Simulations are sampled with the same sampling rate as the data.

To fit the *NOS*-knockout model to cPRC Ca^2+^ traces, we modify the second objective 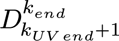. Instead of minimising the difference on the whole interval *t*_*k*_*UV end*_+1_ to *t_k_end__*, we minimise it on the last 10 seconds of this interval, i.e., *t*_*k*_*end*−10[*s*]__ to *t_k_end__*. We change the interval in this particular case because, based on the averaged data, we assume that the only dynamics after the end of UV stimulation in the *NOS*-knockout is linear decay to a steady state. Using the original, non- modified cost function for the recordings exhibiting more complicated dynamics than linear decay would bias the fitting process.

To solve the optimisation problem with the above objective function, we employ a global optimisation method known as a genetic algorithm (GA). GAs are a class of iterative, gradient- free algorithms inspired by processes of natural selection including breeding and mutation (Deb, 2001). With reference to natural selection, sets of candidate solutions in GAs are referred to as ‘populations’. As our algorithm of choice, we use the Matlab implementation of a genetic algorithm for multiobjective optimisation `gamultiobj`. The `gamultiobj` algorithm uses a controlled, elitist genetic algorithm (a variant of NSGA-II (Deb, 2001)). We use the default settings for most hyperparameters of the algorithm.

Here we outline the main steps of the gamultiobj implementation of the GA. The first step is to generate a set of random model parameter sets, known as the initial population. Next, we score the fitness of each parameter set in the population according to the objective function **D**. Using these scores, the `gamultiobj` algorithm employs a binary tournament selection function (Miller et al., 1995) to select ‘parents’ for the next population. A population of children (i.e., new parameter sets) is generated by ‘breeding’ randomly chosen pairs of parents (exchanging subsets of parameter values between them). Parameter values in these children can also be changed by ‘mutations’, which perturb said values randomly by a small amount. Breeding and mutation introduce a level of randomness to the optimisation process that allows exploring the objective function (fitting) landscape. The stochastic nature of GAs helps them to avoid local minima that can plague gradient-descent-based optimisation methods.

In the next step, the `gamultiobj` algorithm scores the parameter sets in the children population using the values of the objective function. Scores from the parent and child populations are combined. A subset of the size of the initial population is then selected as a new parent population based on the scores and the *Crowding Distance*, distance to other parameter sets in terms of a sum of element-wise differences between values of the objective function (used to ensure diversity of the parent population) (Deb, 2001; Konak et al., 2006). This iterative process is repeated until the convergence criteria are met, indicating that the GA has found a set of model parameters that best reproduce the data under consideration.

The `gamultiobj` algorithm returns a set of model parameter values on a Pareto front (Censor, 1977; Deb, 2001). The coordinates of points on the front are the values of the components of the objective function **D**. Points on the Pareto front are nondominated, meaning that improvements to one of the components cannot be achieved without worsening the other. To identify a single optimal solution, we select the parameter set along the Pareto front whose objective function value is the smallest with respect to the Euclidean norm.

Below we present a summary of the steps of our optimisation pipeline for fitting the experimental recordings.

- When fitting to any of the data we use a minimal set of equations necessary to capture the dynamics. This is feasible because a number of equations involve coupling in a feed forward manner, meaning that they do not directly affect the ‘up-stream’ dynamics.

‣ We use equations *C_P_* (*t*), *B*(*t*) and *S*(*t*) from the *NOS*-knockout model to estimate values of the following 6 parameters: *δ_C_P__*, *κ_UV_*, *κ_S_*, *δ_B_*, *B*_0_ and *δ_S_*. We fit them to the Ca^{2+} recordings from the *NOS*-knockout cPRC cells for 20 second UV stimulation.
‣ We use equations *S*(*t*) and *C_N_* (*t*) (equations are the same in the WT and *NOS*-knockout models) to estimate values of *κ_C_N__*, *δ_C_N__*. Estimation is based on the Ca^2+^ recordings from the WT and *NOS*-knockout INNOS cells for 20 second UV stimulation. We use both conditions together because they produce similar dynamics.
‣ We use the equations *S*(*t*), *C_N_* (*t*) and *N*(*t*) from the WT model to estimate values of *κ_N_*_,1_, *κ_N_*_,2_, *δ_N_*. Estimation is based on the NO recordings from the INNOS cells for 20 second UV stimulation. Due to the unknown dissociation dynamics of the NO fluorescent marker, we fit the model to the data using only a single objective, 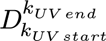.
‣ Finally, we use the full set of equations of the WT model to estimate the remaining 15 para- meters; 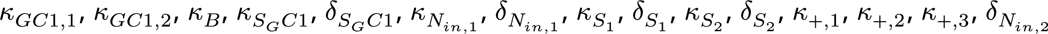.
Set the size of the initial population to 2,000. This value provides a compromise in terms of the coverage of the parameter space and run-time of the GA.
- Set bounds of random distributions of parameters. We draw the initial parameter values of *δ_C_P__*, *κ_UV_*, *κ_S_*, *δ_B_*, *B*_0_ from uniform distributions over the interval [0, 2]. For *δ_S_* we use an interval [0.2, 2]; we increase the lower bound to ensure that the *S*(*t*) returns quickly to a steady state after the end of the UV stimulation; without this restriction, good fits were obtained with the variable switching to a high value. For *κ_C_N__*, *δ_C_N__* we use an interval [0, 2]. For *κ_N_*_,1_, *κ_N_*_,2_, *δ_N_*, we use an interval [0, 50]. For 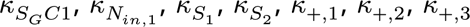 we use an interval [0, 2]. For *κ_GC_*_1,1_, *κ_GC_*_1,2_, *κ_B_* we use an interval [0, 35]. For 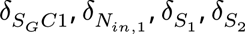 we use an interval [0.1, 2]; we increase the lower bounds to ensure that the variables return to steady states. For *κ*_+,3_ we use an interval [0, 0.0001]; we fix the value of this parameter because the recordings for the longer stimulation times are not consistent with a model in which 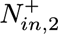 variable starts to return to levels before UV stimulation. For *δ*_*N*_*in*,2__ we use an interval [0.2, 2]; we increase the lower bound to ensure that the variable returns to a steady state.
- Evaluate the objective function on the initial population.
- Run the `gamultiobj` algorithm.
- Use the Euclidean norm to find the set of model parameters that best fits the data.

We fit the model to average recordings in a sequential manner, by which we mean that once we find the best fitting parameter values for a dataset (cell type, condition, signal), we keep them fixed when fitting the remaining parameters to the other datasets. Fixing a subset of parameters allows us to ensure that model parameters inferred from one kind of recordings stay relevant when fitting the model to the other data. This means that the fitted model is extended to include other signals, rather than being completely reparamatrised for each dataset. This is particularly important since technical restrictions mean that we do not have simultaneous recordings across multiple cell types. Furthermore, this sequential approach allows us to represent different experimental conditions by changing a value of a single parameter while keeping all the other parameters fixed. In particular, by lowering *κ_GC_*_2_ in the *S*(*t*) equation of the *NOS*-knockout model, we obtain the *NIT-GC2*-morphant model. Similarly, by setting *κ*_*S*_*GC*1__ to 0, we transform the WT model into the *NOS*-knockout model (Figure 6 – figure supplement 1).

In order to demonstrate the relevance of the parameters fitted to the average datasets, *p_avg_*, for fitting the individual datasets, we use them to build the intervals for fitting the individual datasets. The intervals are [0.25*p_avg_*, 1.75*p_avg_*] for *NOS*-knockout cPRC dataset as well as the parameters that are estimated using the other datasets for WT, *NOS*-knockout and NO INNOS datasets. For the parameters estimated using the other datasets the intervals are the ranges of parameters obtained from the individual fits.

### Predictions from a range of stimulation protocols

We validate the model by comparing model simulations with cPRC Ca^2+^ data collected in experiments with varying duration and intensity of the UV stimulation. The Ca^2+^ data exhibit the following general features:

- the NO-induced response begins following a fixed delay after the start of the UV stimulation;
- the NO-induced response restores the final steady state to the level prior to the UV stimulation (for approximately 20 sec UV stimulation);
- UV stimulation decreases the final steady state;
- after the end of the UV stimulation there is an exponential decay to the steady state;
- A nonlinear effect of UV stimulation intensity. Stimulation with 25% intensity induces lower delay response. Stimulations with 50-100% intensities induce delay responses with comparable amplitudes.

Model simulations with longer stimulation durations and varying stimulation intensity use the wild-type model and parameters fitted to the mean of wild type cPRC recordings under 100% intensity UV stimulation for 20 sec. The simulations capture all the main features of the data described above. The validation is limited by the available data (there is a single recording for each condition and these recordings are taken from different larvae). The data with variable stimulus duration were not used to fit the model parameters.

The model allows us to systematically vary stimulus conditions. To generate new insights from the model, we perform a series of simulations in which we simulate responses to a two-pulse UV stimulation protocol in which we systematically vary the pulse duration ∈ {2, 20, 40} and inter-pulse interval ∈ {10, 30} of the *UV* (*t*) variable in the WT model.

Figure 6 – figure supplement 8 shows how the results of simulations of the WT model depend on these two stimulation parameters. We observe that a single 2 sec pulse leads to a small delayed response. Depending on the inter-pulse interval, the second pulse can increase the amplitude of the delayed response and increase the final steady state value (10 sec interval) or behave almost like an independent stimulation (30 sec interval). In case of 20 sec pulses, the timing of the second pulse coincides with the delayed response and increases its amplitude. This increase is limited by the overlap between the second pulse and the delayed response. Increasing the inter-pulse interval from 10 to 30 sec results in a decrease of the final steady state. For the 60 sec pulse, we observe that the second pulse arrives after the delayed response and that it does not induce a second delayed response. We further observe that, as in the case of 20 sec pulse, the second pulse decreases the final steady state value. Overall, the simulations demonstrate sensitivity of the delayed NO-induced response to the timing and duration of the second pulse. They also illustrate relative independence of UV-phototransduction and delayed NO-induced response and that the presented model does not include a term that would replenish 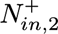(*t*) state variable; consistent with the data presented in Figure 6 – figure supplement 7.

### Assessment of model sensitivity

To analyse model sensitivity, we use Sobol’s indices using the Global Sensitivity Analysis Matlab Toolbox (Cannavó, 2012). Global sensitivity indices facilitate measuring the extent to which changes in model output depend on the changes in parameter values. The Sobol approach is based on the ANOVA (ANalysis Of VAriance) decomposition and computation of the variance of the terms in the decomposition. Here, we combine information from two kinds of indices. The first order indices *S_i_* represent the expected percentage reduction in output variance when parameter *i* has no uncertainty (Cannavó, 2012). By contrast, the total variance indices 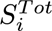 represent the expected percentage output variance that remains if all parameters except *i* are known (Cannavó, 2012). The summary of the significance of global sensitivity indexes is presented in Table 1. For our analysis we defined the output as the sum of the two components of the objective functions. To compute the indices the toolbox explores how the output changes as the model parameters are varied.

The results of the global sensitivity analysis are presented in Tables 2–6. The analysis identifies the parameter *B*_0_ as the most important parameter in the *NOS*-knockout model and *δ_C_P__* as the least important parameter. Fits to the Ca^2+^ recordings in the INNOS cells are found to depend on *δ_S_* and *K_C_N__*, while the most important parameter for fitting the NO recordings in the INNOS cells is *δ_N_*. In case of the wild-type cPRC recordings, the most important parameters are the ones that can be related to the NIT-GC1 dependent NO signals. For the quick NO response *N_in_*_,1_(*t*), the important parameters are *κ*_*S_GC_*1_ and *δ*_*S_GC_*1_, while, *κ*_*S_GC_*1_ and *δ*_*S*_GC_1_ control the dynamics of the *N_in_*_,1_(*t*) variable. For the late NO response *N_in_*_,2_(*t*), the important parameters include *κ*_*GC*1,2_, *K*_*S*_1__ and *K*_*S*_2__. The parameter *κ_GC_*_1,2_ modulates Ca^2+^ levels in the cPRC cells. Finally, *K*_*S*_1__ and *K*_*S*_2__ are important for regulating the signalling cascade that leads to the late NO response.

### Assessment of model identifiability and observability

The analysis of structural identifiability and observability provides an a priori assessment of whether a given model structure permits a unique solution to the parameter estimation procedure in an ideal scenario of continuous, noise-free observations. Structural identifiability determines whether the unknown parameters can, in principle, be uniquely recovered from this idealised input-output data. Observability evaluates whether the unmeasured state variables of the dynamic model can be accurately inferred from those same external observations (Díaz-Seoane et al., 2022).

To analyse identifiability in our model, we use the probabilistic ProbObsTest algorithm as implemented in the STRIKE-GOLDD 4.0 Matlab toolbox (Díaz-Seoane et al., 2022). The results of the ProbObsTest algorithm are guaranteed to be correct for identifiable/observable variables and have high probability of being correct for unidentifiable/unobservable variables.

This analysis revealed that the variables *S_GC_*_1_(*t*), *N_in_*_,1_(*t*), *S*_1_(*t*) and *S*_2_(*t*) are unobservable, meaning that it is theoretically impossible to deduce the true values of these variables, even with perfect experimental data. They either do not affect the experimental measurements at all, or their effects are mathematically indistinguishable from changes in other state variables or parameters (they are lumped together). Specifically, it implies that the amplitudes of these variables in our model cannot be uniquely determined. The algorithm further identified parameters *κ*_*S*_*GC*1__, *κ*_*N*_*in*,1__, *κ*_*S*_1__, *κ*_*S*_2__, *κ*_+,1_, *κ*_+,2_, *κ_GC_*_1,1_ as unidentifiable, meaning that there exist multiple combinations of these parameters that result in identical simulated traces; these parameters are not correlated with each other; in addition, *κ*_*S*_*GC*1__ and *κ*_*S*_1__ are not correlated with any other parameters (see Figure 6 - figure supplement 8). These results are to be expected considering the structure of the proposed model. First, all unidentifiable parameters scale the amplitudes of upstream signals (the input). Second, all the unobservable state variables are given by equations that represent the simplest low-pass filter (a linear transformation) of upstream signal (the input). These variables are included in the model to delay and scale inputs from other parts of the model. The ProbOpsTest algorithm suggests that eliminating parameters *κ*_*S*_*GC*1__, *κ*_*N*_*in*,1__, *κ*_*S*_1__ and *κ*_*S*_2__ would result in full state and parameter observability of the model. Since the structure of this part of the model is very likely to be modified by insights from future experiments, we accept the outcomes of the analysis without modifying the model. Furthermore, sensitivity analysis identified parameters *κ*_*S*_*GC*1__, *κ*_*S*_1__ and *κ*_*S*_2__ as the ones that are most important for fitting the WT model to the data. As such, removing these parameters from the model may impact the global sensitivity analysis for the remaining parameters.

## Data analysis and figure preparation

Data were analysed in ImageJ and R. All figures were assembled in R with the cowplot and patchwork packages. All scripts are available at https://github.com/JekelyLab/Joukra_et_al_NOS. The matlab code for the mathematical model is in the same repository under /code/matlab/.

## Acknowledgements

We thank Dr. Cameron Hird, Adam Johnstone, Rebecca Turner and Jamie Haddon for animal husbandry, Dr. Corin Liddle for advice on calcium imaging, and Dr. Emelie Brodrick and Dr. Elizabeth Williams for critical reading of the manuscript. This work was funded by the Wellcome Trust (214337/Z/18/Z). This project has received funding from the European Research Council (ERC) under the European Union’s Horizon 2020 research and innovation programme (grant agreement No 101020792). KJ has been supported by a JSPS Overseas Research Fellowship, LAYG by a BBSRC Discovery fellowship (BB/W010305/1), PS by a Wellcome Trust Institutional Strategic Support Award (204909/Z/16/Z), and KCAW by the EPSRC Hub for Quantitive Modelling in Healthcare (EP/T017856/1). We thank Dr. Francesca Carlisle for the mycoplasma testing of COS-7 cells.

## Figure supplements

**Figure 1 – figure supplement 1.**
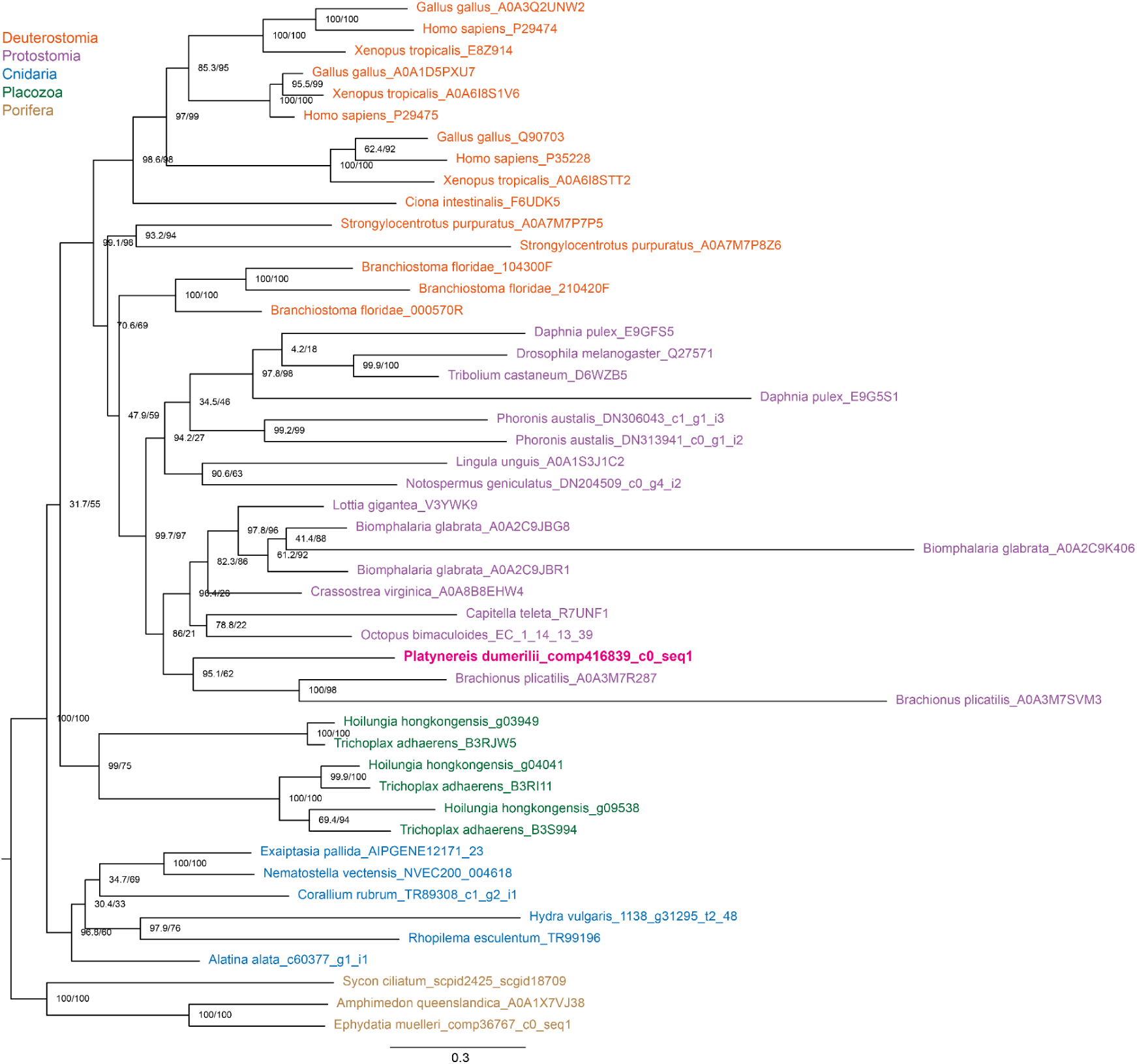
Maximum-likelihood phylogenetic tree of NOS protein sequences. Individual branches are coloured by taxonomy. Branch support values indicate UFBoot and aLRT-SH-like values. Figure 1 – figure supplement 1 – source data 1. NOS sequences used for the phylogenetic reconstruction. Figure 1 – figure supplement 1 – source data 2. Aligned and trimmed NOS sequences used for the phylogenetic reconstruction. Figure 1 – figure supplement 1 – source data 3. Tree file of the reconstructed NOS phylogeny.

**Figure 1 – figure supplement 2.**
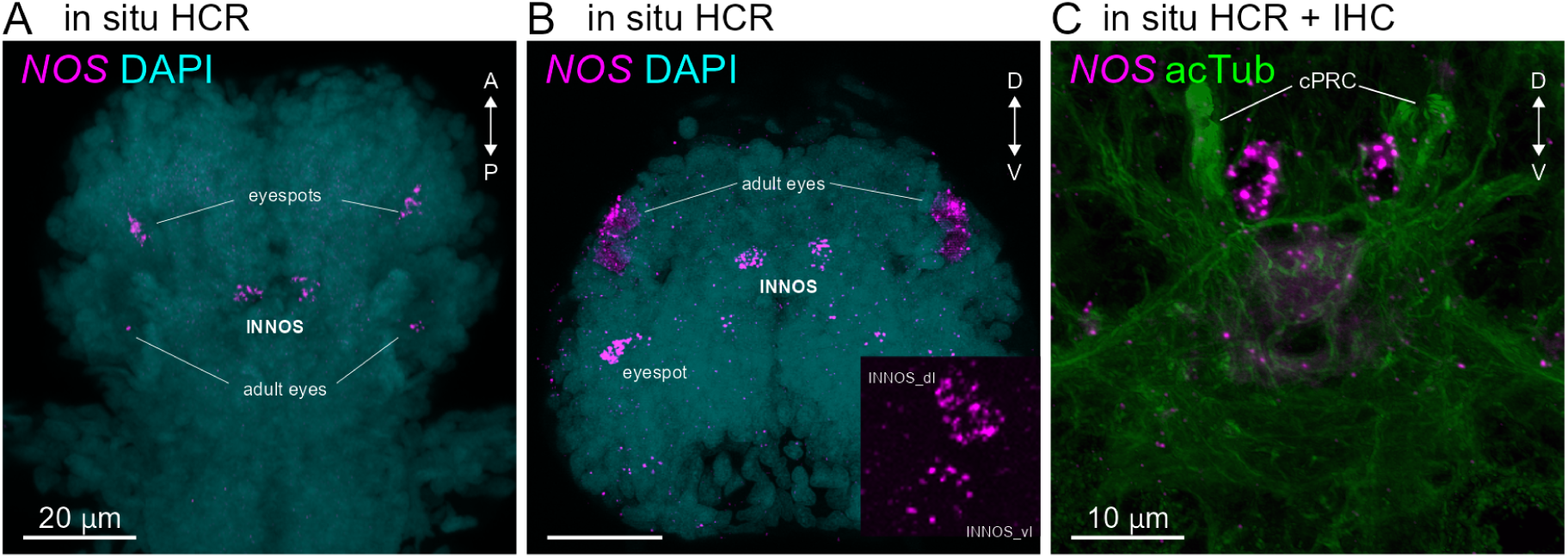
Expression of *NOS* in 3-day-old *Platynereis* larva. **(A)** HCR in situ for *NOS* (magenta), dorsal view. (B) HCR in situ for *NOS* (magenta), ventral view. Inset in **(B)** shows close-up of two *NOS*-positive cells (INNOS_dl and INNOS_vl) on the left side. Samples were also stained for DAPI (cyan) to label nuclei.(C) Expression of the *NOS* gene detected by in situ HCR (magenta) in a 3-day-old larva (anterior view). Antibody staining for acetylated α-tubulin (acTub: green) highlights cPRC cilia and the neuropil.

**Figure 1 – figure supplement 3.**
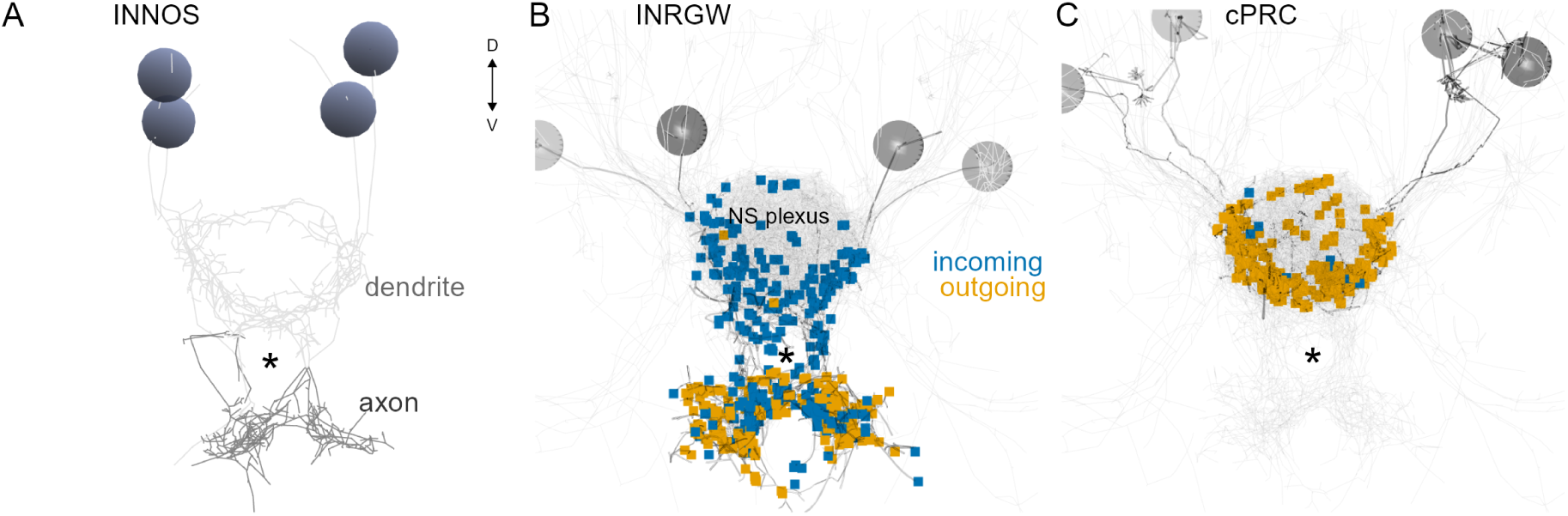
Synapse distribution in cPRCs and possynaptic interneurons. **(A)** Axo- dendritic splitting of the INNOS neurons. **(B)** Volume rendering of the four INRGWa neurons with incoming and outgoing synapses. **(C)** Volume rendering of the four cPRC neurons with incoming and outgoing synapses. All images are anterior views. Asterisk marks the same position across images for reference. Grey speheres indicate the position of cell nuclei. NS plexus; neurosecretory plexus. The radius of INNOS nuclei is set to 2 µm for scale.

**Figure 3 – figure supplement 1.**
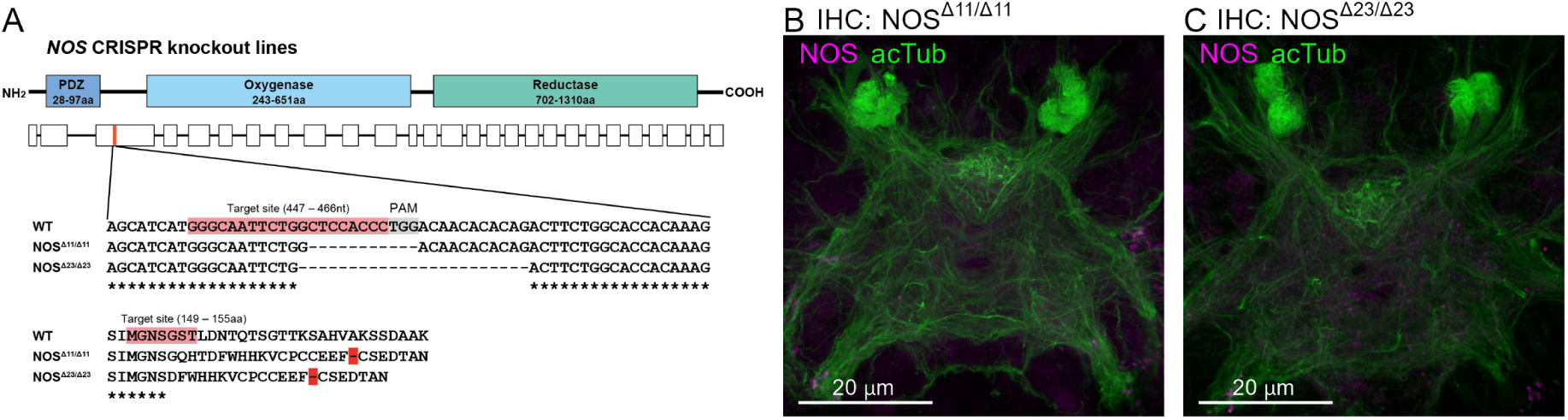
Generation and behavioural characterisation of *NOS* CRISPR knockouts. **(A)** Top: The domain organisation of *Platynereis* NOS protein and exon/intron structure of the *NOS* gene. Bottom: The genomic locus of *NOS* with the CRISPR target site, wild-type and knockout (NOSΔ11, NOSΔ23) sequences and predicted protein sequences. The PAM sequence is shown in grey, stop codons in red. **(B, C)** Immunostaining for NOS antibody in NOS mutant (B, NOSΔ11/Δ11; C, NOSΔ23/Δ23) larvae. Acetylated α-tubulin staining (green) highlights the neuropil and cPRC cilia. Anterior view of a 3- day-old larva.

**Figure 3 – figure supplement 2.**
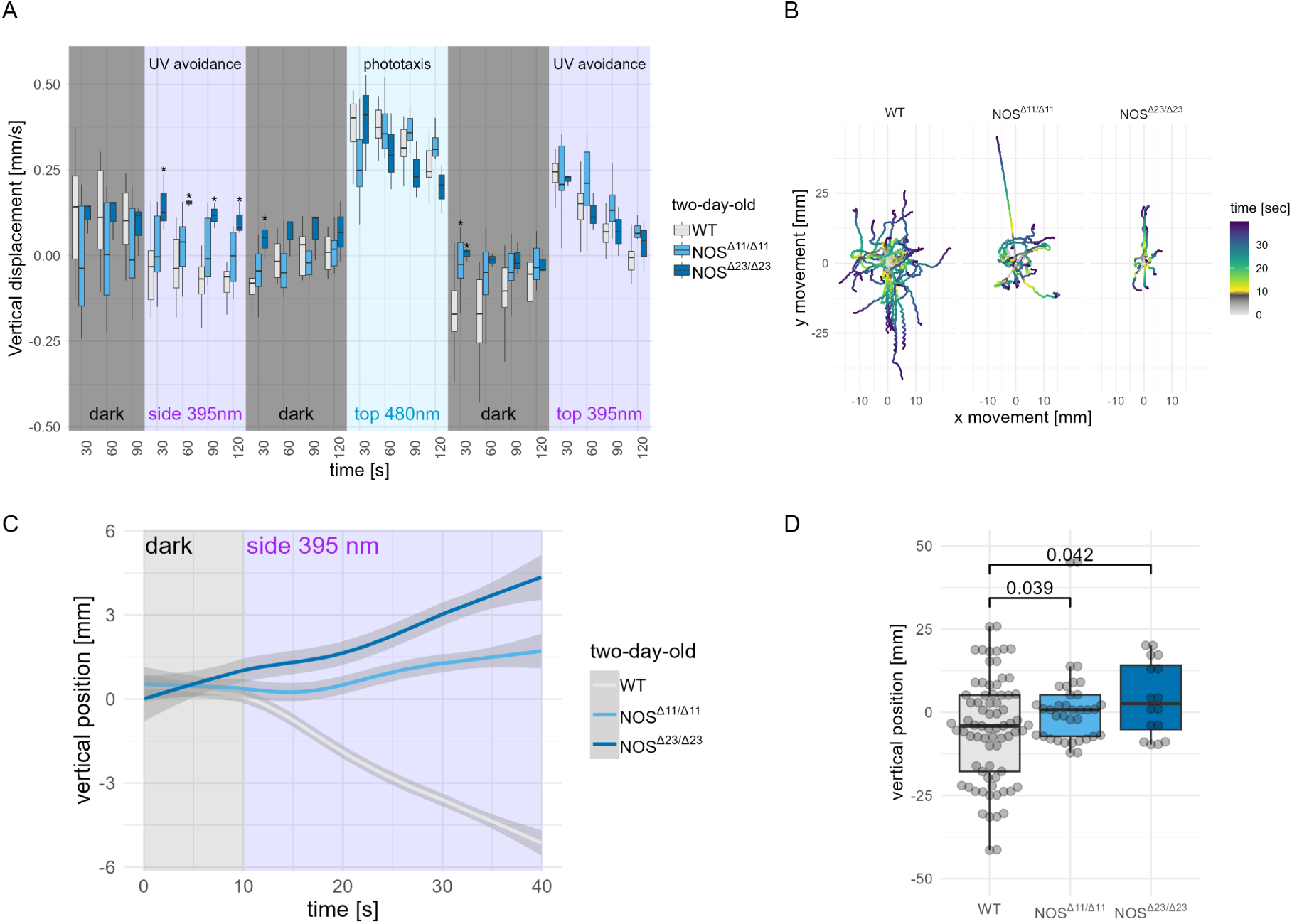
Generation and behavioural characterisation of *NOS* CRISPR knockouts. **(B)** Swimming trajectories of wild type (WT, n=37) and *NOS* mutant (NOSΔ11/Δ11, n=18 and NOSΔ23/Δ23, n=8) 2-day-old larvae. All trajectories start at 0 x and y position and time 0 corresponding to 10 sec before the onset of 395 nm stimulation from the side. **(C)** Vertical displacement in 30 sec bins of wild type (WT, n=37) and mutant (NOSΔ11/Δ11, n=18 and NOSΔ23/Δ23, n=8) 2-day-old larvae stimulated with 395 nm light from the side, 488 nm light from the top and 395 nm light from the top. **(D)** Comparison of vertical position of wild type and mutant 2-day-old larvae after 30 seconds of UV stimulation. One-way ANOVA with Dunnett’s multiple-comparison test are shown. The wild type (WT, n=52) and *NOS* mutant (*NOSΔ11/Δ11*, n=30 and *NOSΔ23/Δ23*, n=16) larvae. **(E)** Vertical position of batches of wild type (12 batches) and mutant (NOSΔ11/Δ11, 8 batches and NOSΔ23/Δ23, 3 batches) 2-day-old larvae over time under 395 nm UV stimulation. The starting position of each larval trajectory was set to 0. One-way ANOVA with Dunnett’s multiple-comparison test are shown. *P < 0.05. Figure 3 – – figure supplement 2 – source data 1. Vertical displacement data for 2-day-old wild type and *NOS* mutant larvae. Figure 3 – – figure supplement 2 – source data 2. Swimming trajectories of 2-day-old wild type and *NOS* mutant larvae. Figure 3 – – figure supplement 2 – source data 3. Vertical position of 2-day-old wild wild type and *NOS* mutant larvae. Figure 3 – – figure supplement 2 – source data 4. Vertical position of 2-day-old wild type and *NOS* mutant larvae after 30 sec UV exposure.

**Figure 3 – figure supplement 3.**
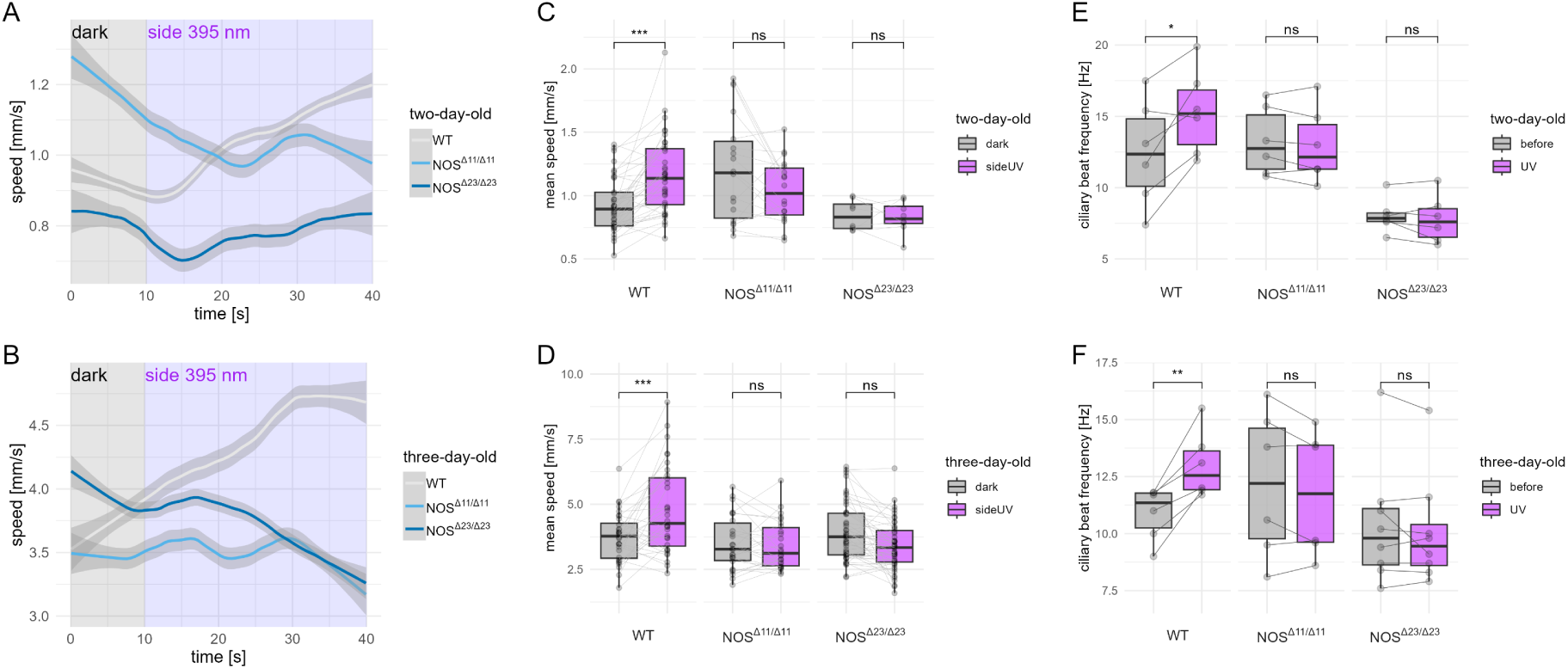
Generation and behavioural characterisation of *NOS* CRISPR knockouts. **(A,B)** Swimming speed of batches of wild type and *NOS* mutant 2-day-old (WT, n=37, NOSΔ11/Δ11, n=18 and NOSΔ23/Δ23, n=8) (A) and 3-day-old (WT, n=32, *NOSΔ11/Δ11*, n=26 and *NOSΔ23/Δ23*, n=47) (B) larvae under 395 nm UV stimulation. **(C,D)** Mean swimming speed of wild type and *NOS* mutant 2-day- old (WT, n=37, NOSΔ11/Δ11, n=18 and NOSΔ23/Δ23, n=8) (C) and 3-day-old (WT, n=32, *NOSΔ11/Δ11*, n=26 and *NOSΔ23/Δ23*, n=47) (D) larvae during 10 seconds before (dark) and 10 seconds after the start of 395 nm UV stimulation (sideUV). Data points for the same larva are joined by lines. One-tailed paired t-test; p- values (*** <0.001) are shown. **(E,F)** Average ciliary beat frequency of wild type and mutant 2-day-old (E) and 3-day-old (F) single larvae during 10 seconds before (gray) and 10 seconds after the start of 395 nm UV stimulation (purple). N=6 larvae each. Data points for the same larva are joined by lines. One-tailed paired t-test; p-values (** <0.01 or * <0.05) are shown. Figure 3 – figure supplement 3 – source data 1. Swimming speed data for wild type and *NOS* mutant 2-day-old larvae. Figure 3 – figure supplement 3 – source data 2. Swimming speed data for wild type and *NOS* mutant 3-day-old larvae. Figure 3 – figure supplement 3 – source data 3. Mean swimming speed of wild type and *NOS* mutant 2-day-old larvae. Figure 3 – figure supplement 3 – source data 4. Mean swimming speed of wild type and *NOS* mutant 3- day-old larvae. Figure 3 – figure supplement 3 – source data 5. Ciliary beat frequency data for wild type and *NOS* mutant 2-day-old larvae. Figure 3 – figure supplement 3 – source data 6. Ciliary beat frequency data for wild type and *NOS* mutant 3-day-old larvae.

**Figure 3 – figure supplement 4.**
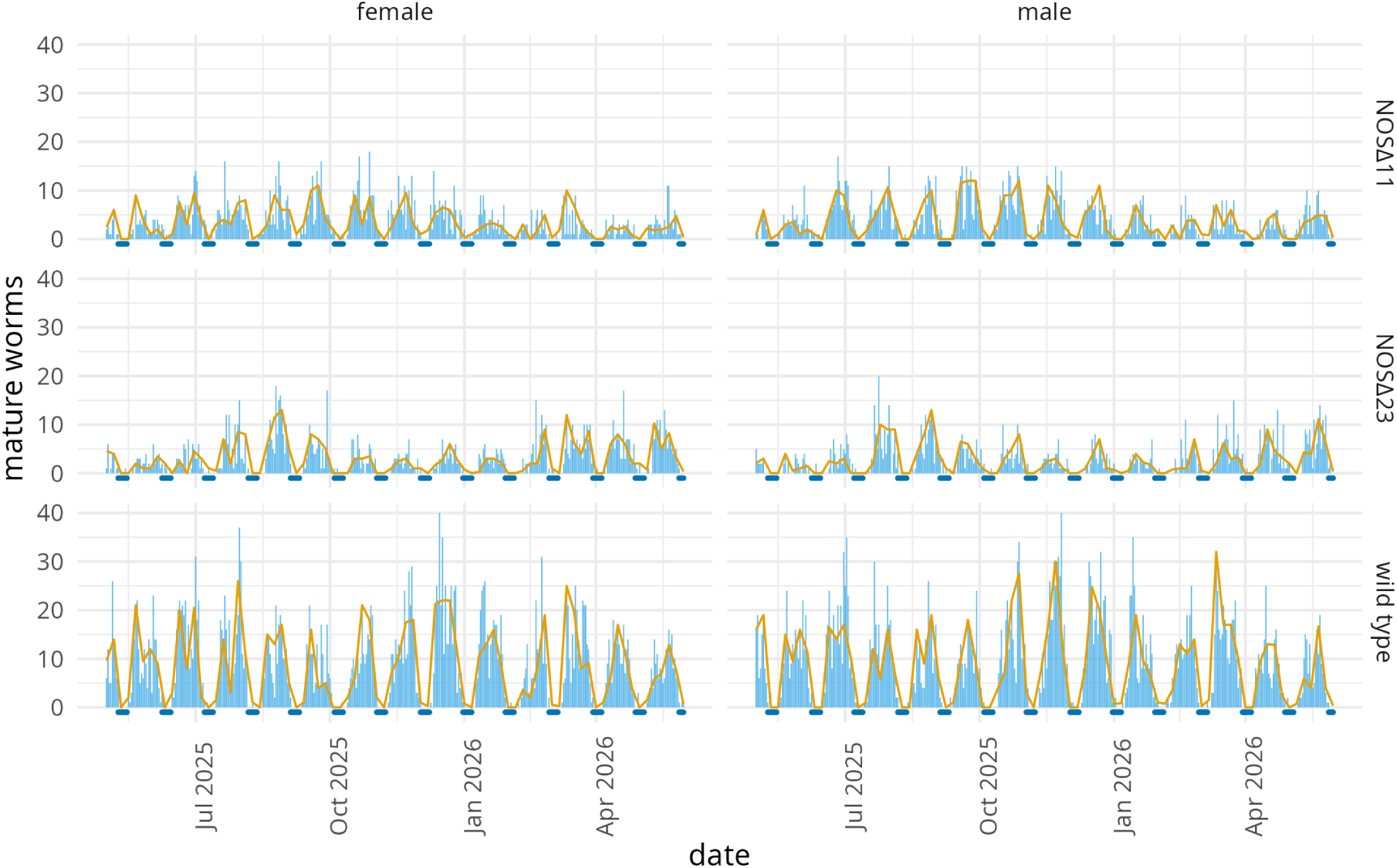
Lunar cycle of maturation in wild-type and *NOS* CRISPR knockout *Platynereis* worms. Data represent the number of swimming epitoks per day over a one-year period in the Heidelberg culture. Males and females are plotted separately. Data are shown for wild type, *NOSΔ11/Δ11* and *NOSΔ23/Δ23* worms as line plots and smoothed plots (with a 0.95 confidence band). Figure 3 – figure supplement 4 – source data 1. Maturation data for wild type and *NOS* knockout worms.

**Figure 4 – figure supplement 1.**
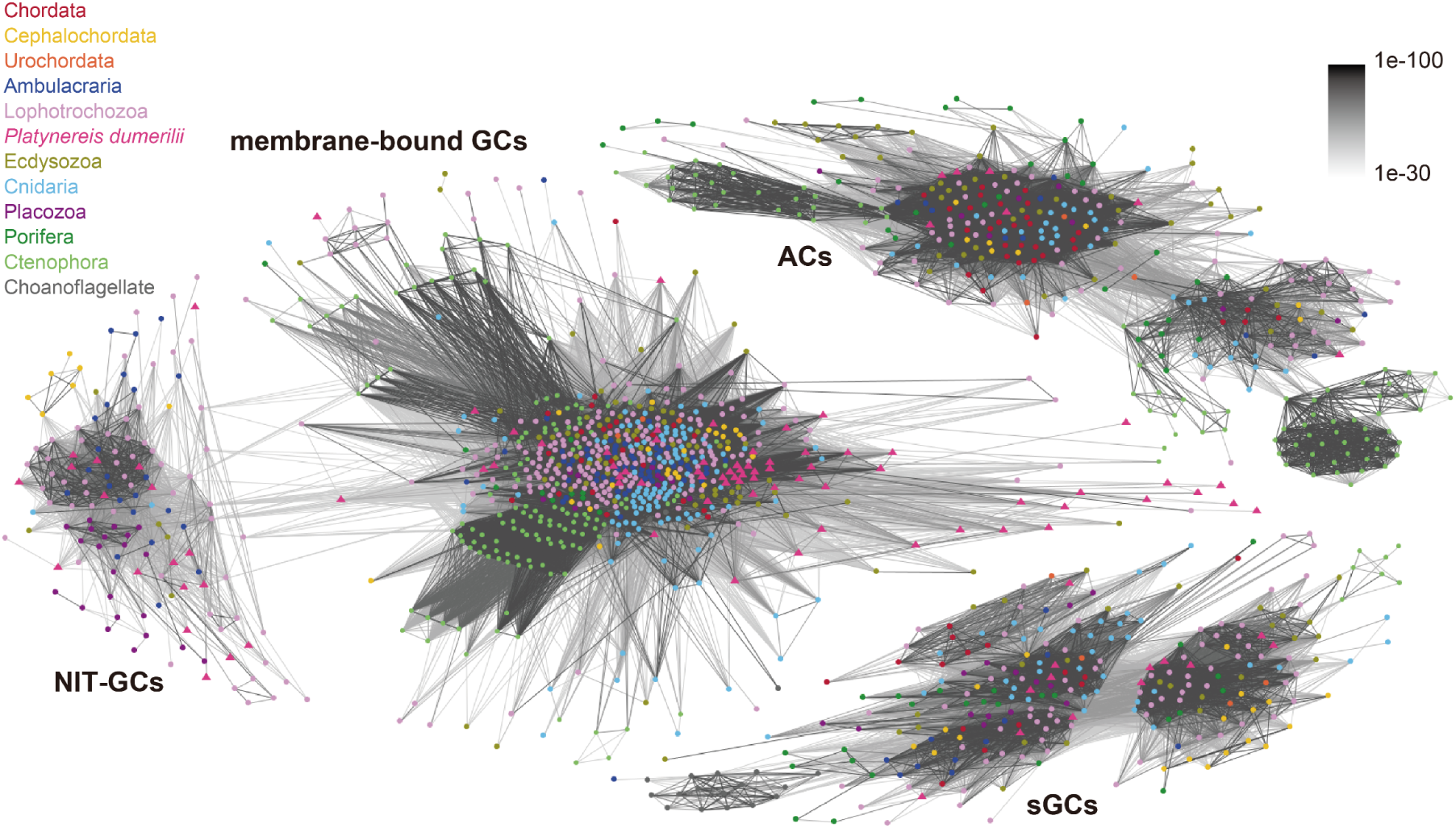
Cluster analysis of guanylate and adenylate cyclase sequences. Each node represents one sequence, colour-coded by taxonomy. Connections represent BLAST P-values of <1e-16. NIT-GCs, NIT domain containing guanylate cyclases; membrane-bound GCs, membrane-bound guanylate cyclases; sGCs, soluble guanylate cyclases; ACs, adenylate cyclases.

**Figure 4 – figure supplement 2.**
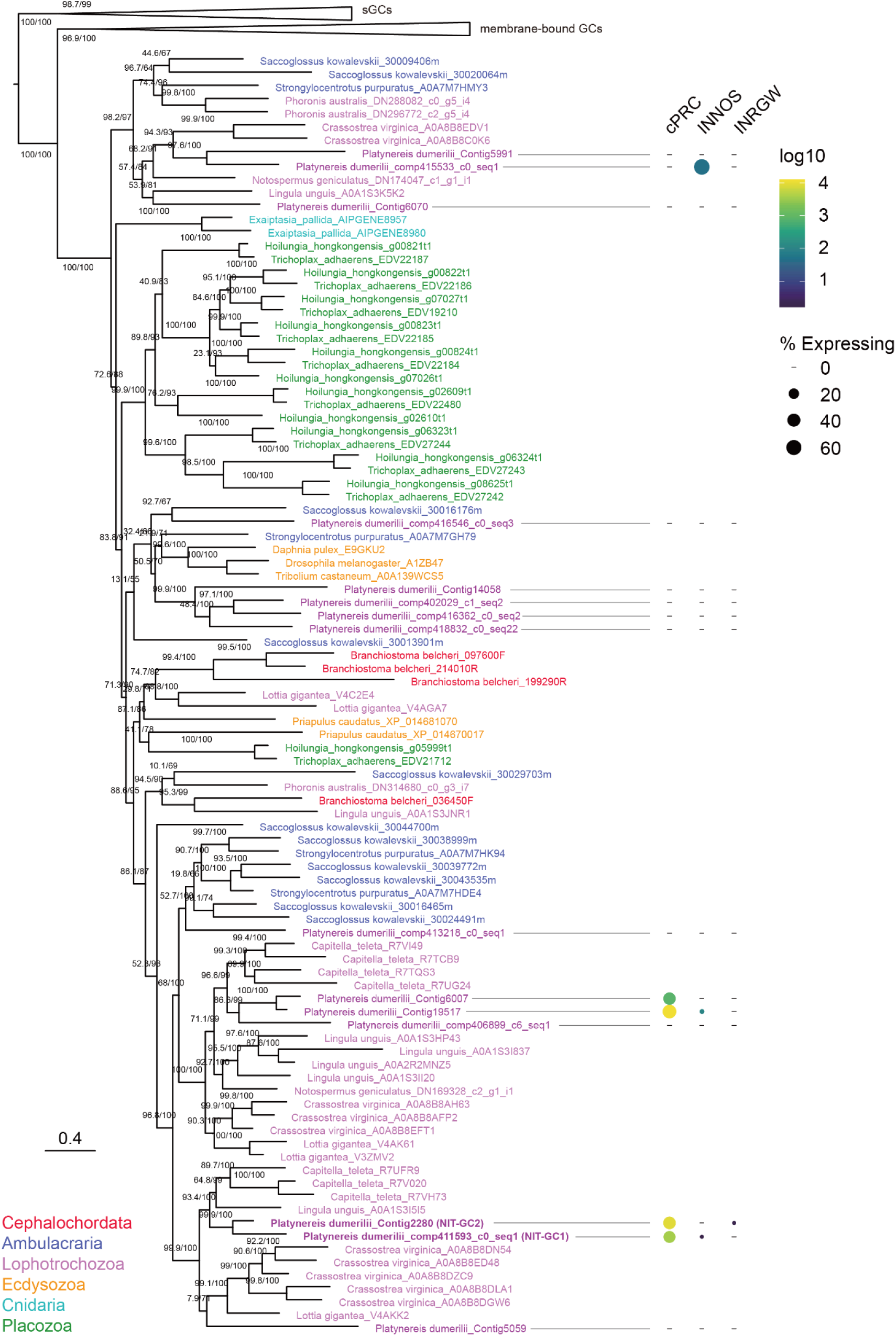
Maximum-likelihood phylogenetic tree of NIT-domain-containing guanylate cyclases. Membrane-bound and soluble guanylate cyclases (sGC) were included as outgroups. Guanylate cyclases with NIT domains are found in most animal phyla except Porifera, Ctenophora, Urochordata and Chordata. Branch support values indicate UFBoot and aLRT-SH-like values. The expression of *Platynereis* NIT-GC genes in cPRC, INNOS and INRGWa cells is indicated on the right side of the tree. The values represent expression based on single-cell sequencing data. Dot size indicates specificity (percent of transcripts expressed in the indicated cell across all cells). Dot colour represents the logarithm of the number of transcripts in the expressing cells. Figure 4 – figure supplement 2 – source data 1. GC sequences used for the phylogenetic reconstruction. Figure 4 – figure supplement 2 – source data 2. Aligned and trimmed GC sequences used for the phylogenetic reconstruction. Figure 4 – figure supplement 3 – source data 3. Tree file of the reconstructed GC phylogeny.

**Figure 4 – figure supplement 3.**
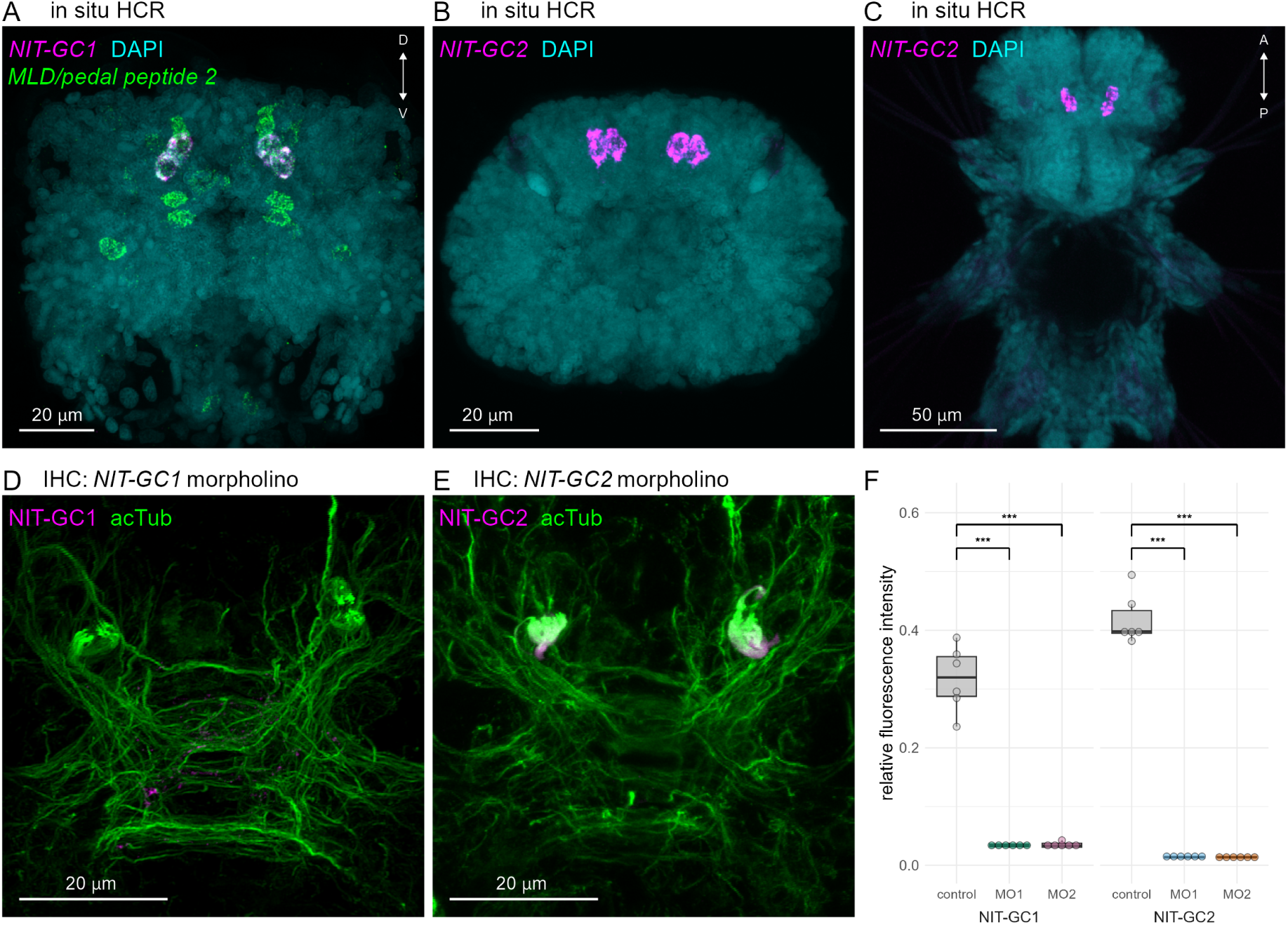
Expression of NIT-GC1 and NIT-GC2 in the cPRCs. **(A)** HCR co- expression analysis of *NIT-GC1* (magenta) and *MLD/pedal-peptide-2 proneuropeptide* (green). Anterior view of a 2-day-old larva. Nuclei were stained with DAPI (cyan). **(B, C)** Expression of *NIT-GC2* (magenta) detected by in situ HCR. Anterior (B) and ventral (C) views of a 3-day-old larva. Nuclei were stained with DAPI (cyan). **(D, E)** Immunostaining for NIT-GC1 (D) and NIT-GC2 (E) antibodies in larvae injected with NIT-GC1 (D) or NIT-GC2 (E) morpholinos. Acetylated α-tubulin staining (green) highlights the neuropil and cPRC cilia. Anterior view of a 2-day-old larva. **(F)** The fluorescence intensities of the NIT-GC1 or NIT-GC2 in the control and morphant (MO1 and MO2) larvae (n=6 each) were normalized to the acetylated α-tubulin signals and plotted as a relative fluorescence. One-way ANOVA with Dunnett’s multiple-comparison test; p-values (‘***‘ <0.001) are shown. Figure 4 – figure supplement 3 – source data 1. Comparison NIT-GCs signals readings.

**Figure 5 – figure supplement 1.**
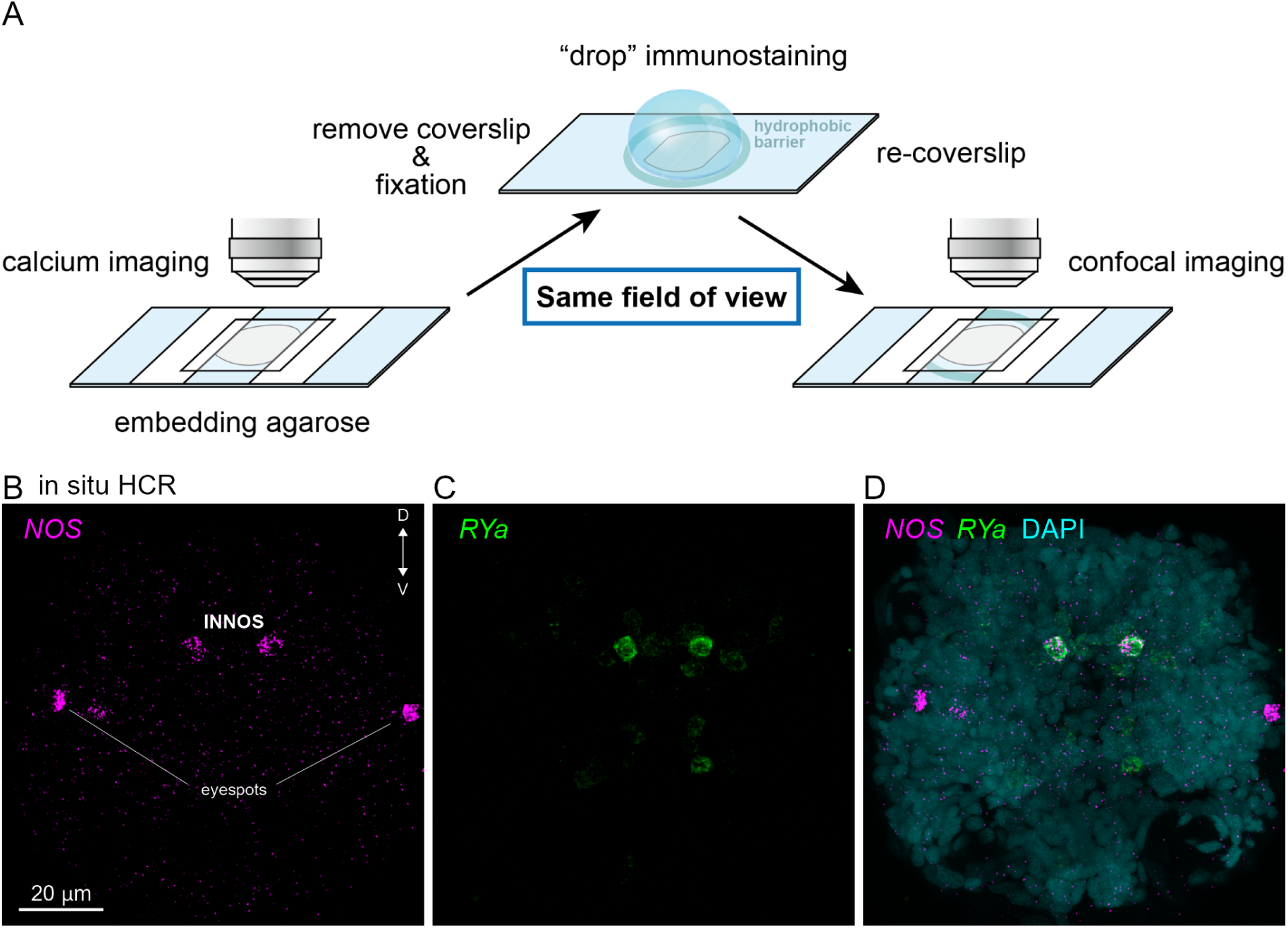
On-slide immunostaining and coexpression analysis of *NOS* and *RYamide proneuropeptide*. **(A)** Schematic diagram of the on-slide immunostaining procedure after Ca^2+^ imaging.**(B-D)** Co-expression analysis by HCR in situ of *NOS* (magenta) and *RYamide proneuropeptide* (green). Anterior view of a 2-day-old larva. Nuclei were stained by DAPI (cyan).

**Figure 5 – figure supplement 2.**
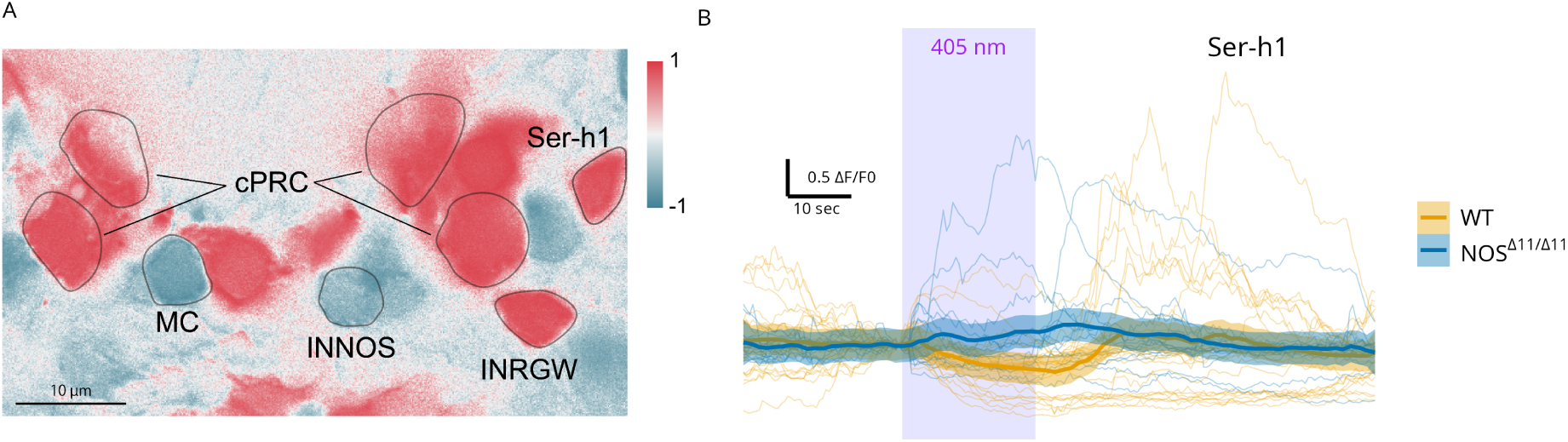
NOS-dependent dynamics of the Ser-h1 neurons. **(A)** Correlation map of neuronal activity of the cPRCs, INNOS, INRGWa, and Ser-h1 neurons. **(B)** GCaMP6s fluorescence in Ser- h1 cells in wild type and *NOSΔ11/Δ11* mutant larvae during 405 nm stimulation. Figure 5 – figure supplement 2 – source data 1. Calcium imaging data from Ser-h1 neurons.

**Figure 6 – figure supplement 1.**
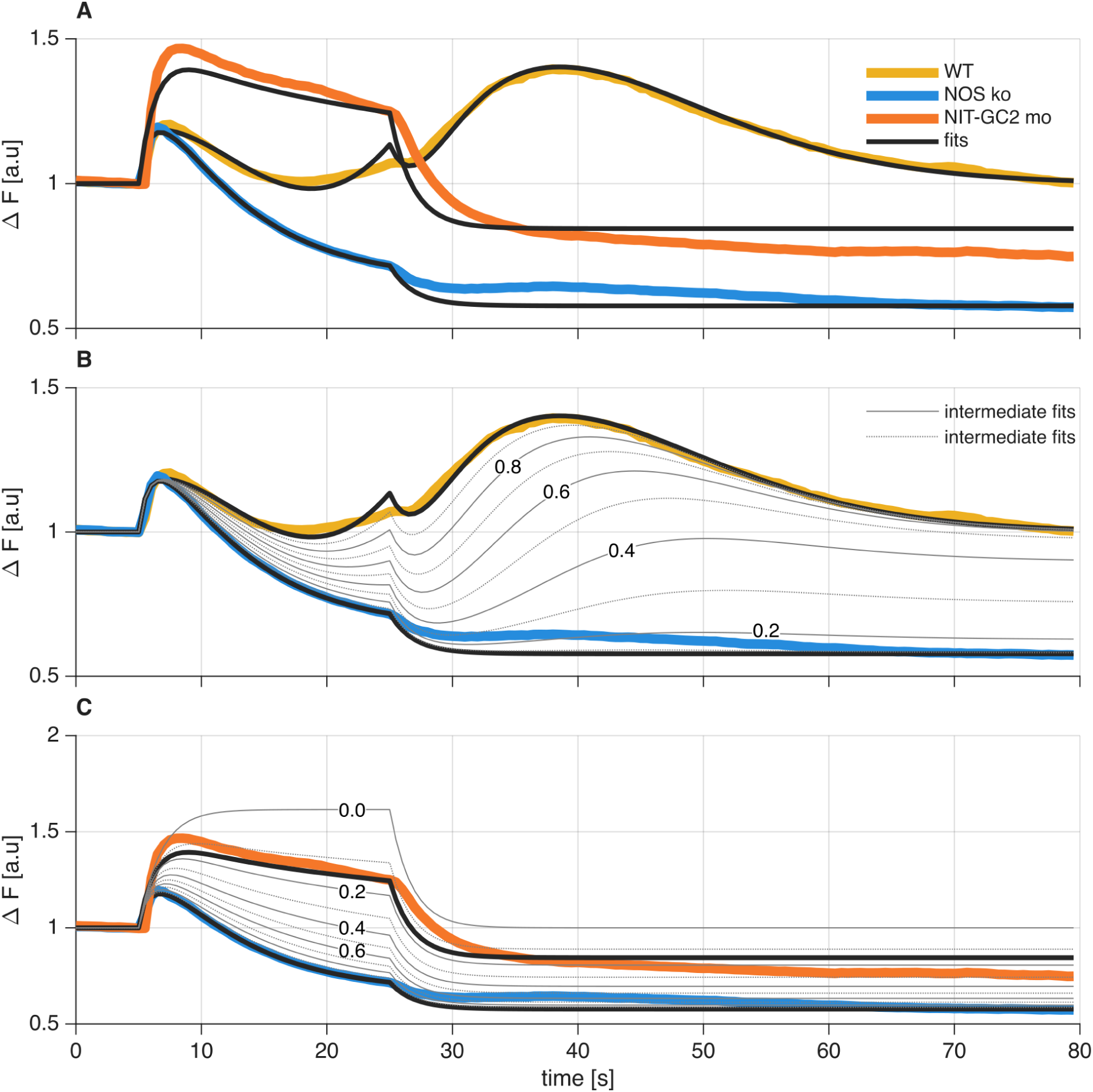
Simulated Ca^2+^ traces for parameter sets fitted to the means of individual cPRC recordings collected in wild-type,*NOS* knockout, and *NIT-GC2* morphant larvae. The NIT- GC2 has the same parameter values as NOS-knockout except *kappa*_*GC*2_ is decreased from 1 to 0.15. **(A)** Coloured curves indicate the mean of the recordings, thick black curves indicate simulated traces based on model parameters fitted to the mean of the recordings. **(B, C)** Thin grey curves (solid and dotted) represent simulated experimental conditions with different levels of efficacy of the NOS-knockout **(B)** and NIT-GC2 morpholino **(C)**. To generate interpolated condition curves in **(B)**, we multiplied *kappa*_*S*_*GC*1__ in the wild-type model by values from 0.1 to 0.9 with steps of 0.1. To generate interpolated condition curves in **(C)**, we use a set of *kappa*_*GC*2_ values from 0.1 to 0.9 with steps of 0.1. Figure 6 – figure supplement 1 – source data 1. Recorded traces (as used in earlier figures) `wt_ko_mo_data.mat`. Figure 6 – figure supplement 1 – source data 2. Script to generate the figure `fig_WT_to_MO.m`

**Figure 6 – figure supplement 2.**
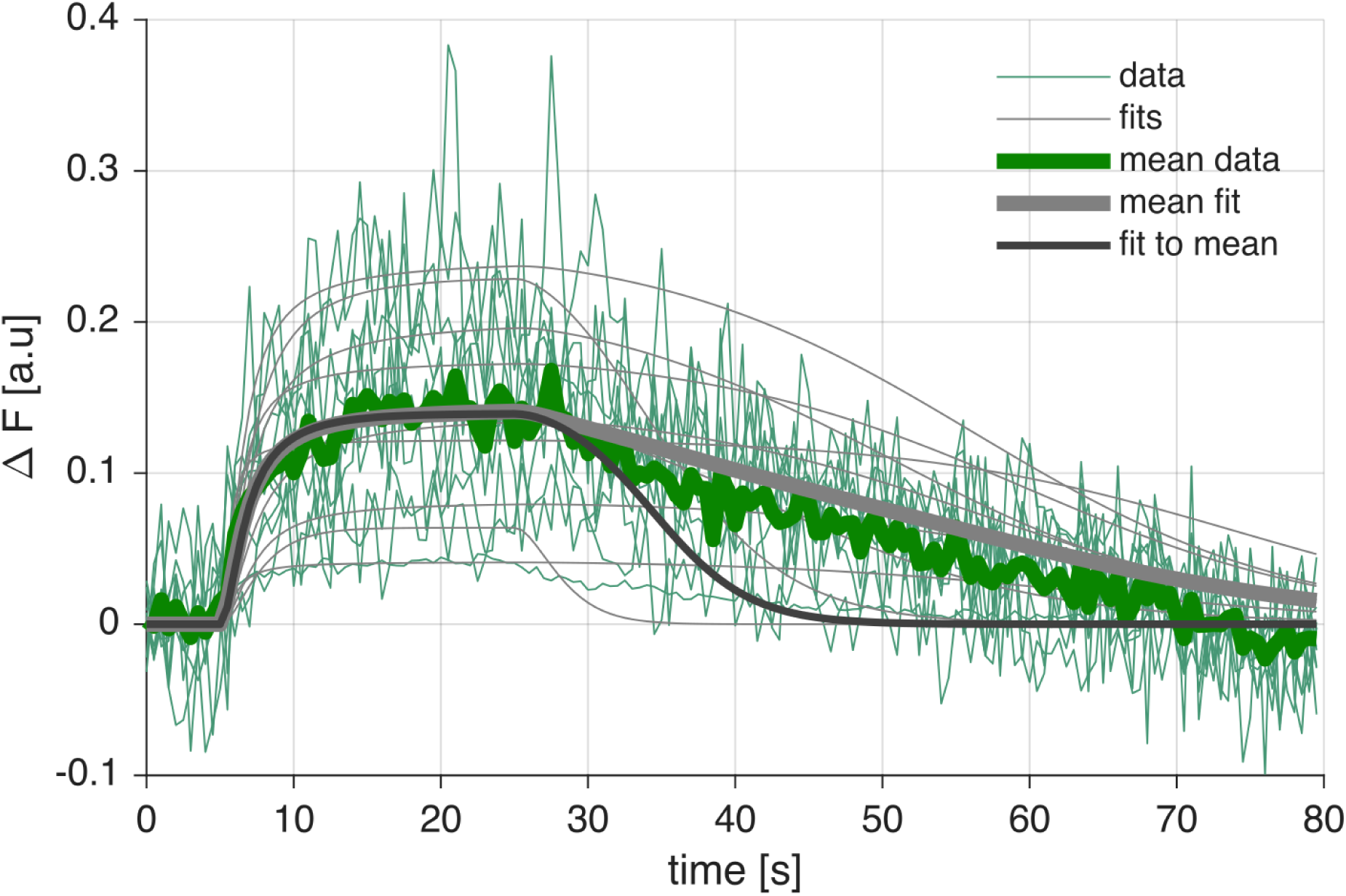
Simulated NO traces for parameter sets fitted to individual NO- recordings collected in wild type larvae. Thin coloured curves indicate individual recordings, thin grey curves indicate simulated NO traces based on model parameters fitted independently to each recording, the thick coloured curve indicates the average of the recordings, the thick grey curve represents the average of the fits, and the thick black curve represents the fit of the model parameters to the averaged recordings. Figure 6 – figure supplement 2 – source data 1. Best fitted parameters `best_pars.mat`. Figure 6 – figure supplement 2 – source data 2. Recorded traces (same as in Figure 2 but with removed bleaching and steady state set to 0) (`NO_data.mat`). Figure 6 – figure supplement 2 – source data 3. Script to generate the figure `fig_fit_NO.m`

**Figure 6 – figure supplement 3.**
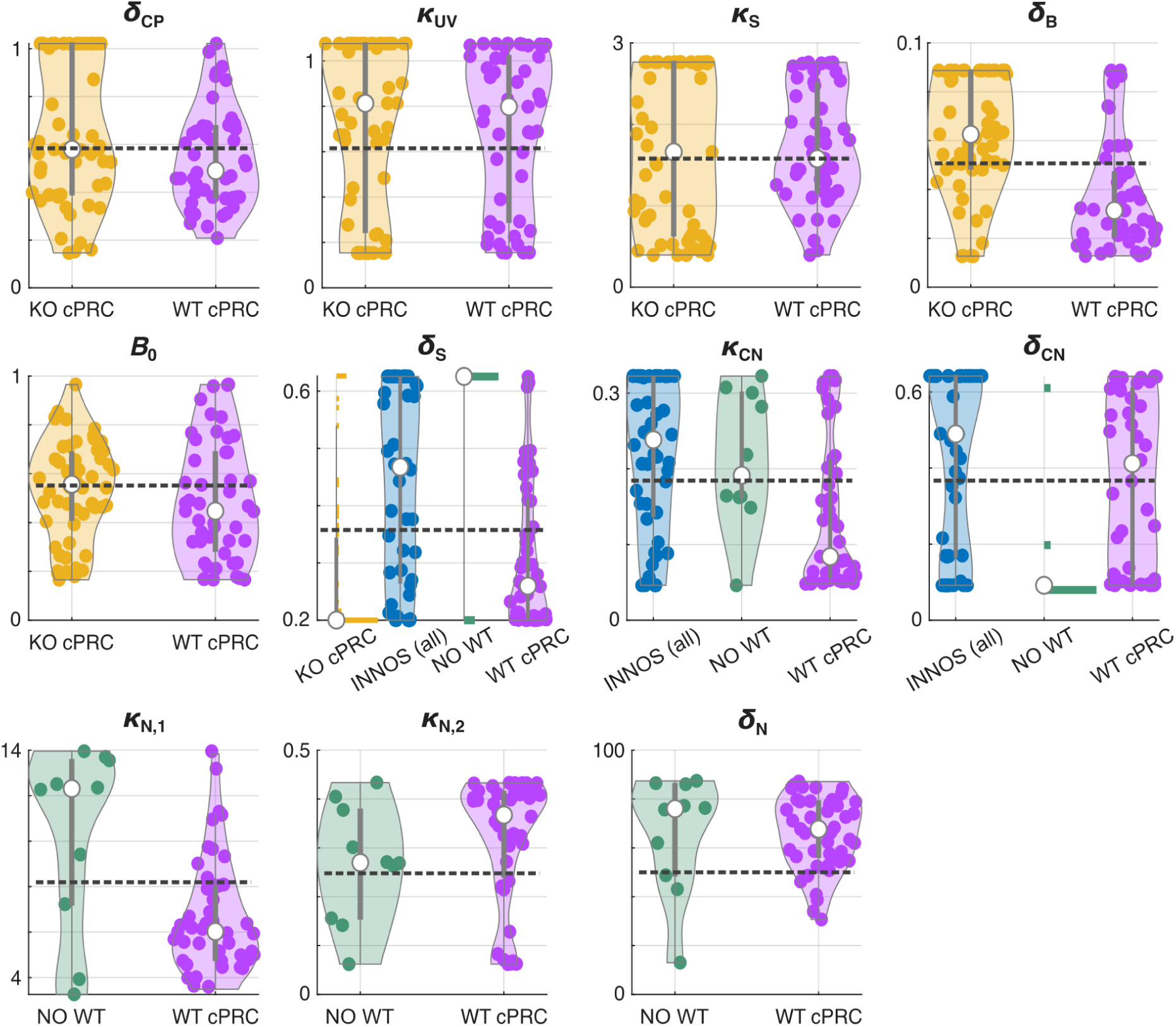
Distributions of the parameter values fitted to the individual recordings collected in different conditions. Violin plots (grey lines) are kernel density estimates of the underlying distributions computed using the Matlab function *ksdensity* with default settings, i.e., using normal kernel function; plots are trimmed to the observed range of data. Coloured markers represent individual parameter values, white markers indicate medians, vertical grey bars indicate interquartiles (ranges Q25 to Q75), and the dashed line indicates the value of the parameter fitted to the average of the recordings. We show histograms instead of violin plots for *delta_S_* when fitted to *NOS*-knockout recordings and for *delta_S_* and *delta_C_N* when fitted to NO recordings because these distributions are highly skewed and hence misrepresented by violin plots. Violin plots are presented in the order of our model-fitting procedure, i.e., first Ca^2+^ in *NOS*-knockout cPRC cells (yellow), then Ca^2+^ in wild type and *NOS*-knockout INNOS cells (blue), then NO in INNOS cells (green), and finally Ca^2+^ in wild type cPRC cells (magenta); (see Methods for details). Figure 6 – figure supplement 3 – source data 1. Best fitted parameters `best_pars.mat`. Figure 6 – figure supplement 3 – source data 2. Script to generate the figure `parameter_dist.m`

**Figure 6 – figure supplement 4.**
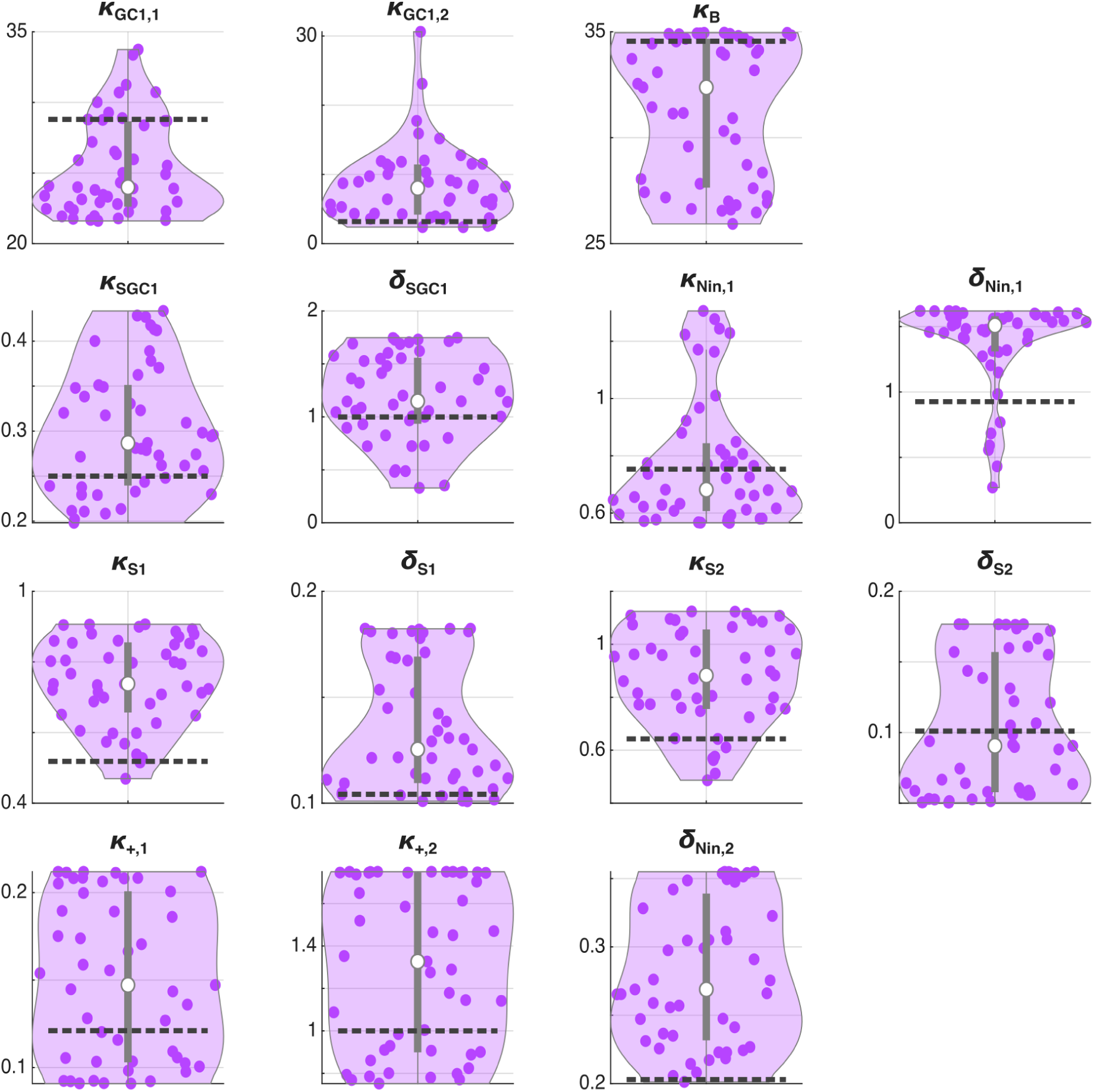
Distributions of the parameter values fitted to individual cPRC recordings collected in wild type larvae. The figure is in the same format as Figure 6 – figure supplement 4. Figure 6 – figure supplement 4 – source data 1. Best fitted parameters `best_pars.mat`. Figure 6 – figure supplement 4 – source data 2. Script to generate the figure `parameter_dist.m`.

**Figure 6 – figure supplement 5.**
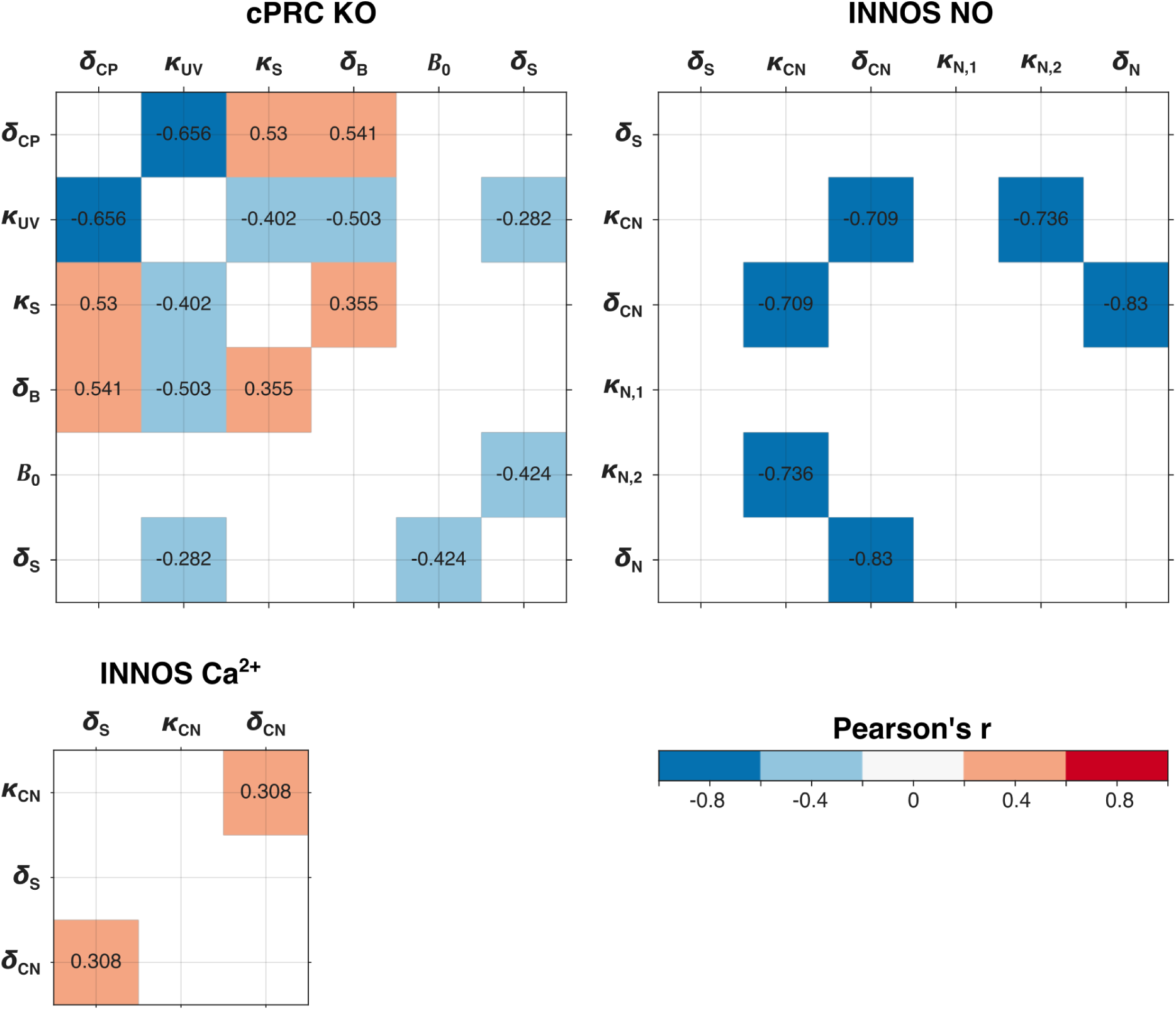
Pairwise correlations between parameters of the *NOS*-knockout model (state variables *C_P_* (*t*), *B*(*t*) and *S*(*t*)) fitted to *NOS*-knockout cPRC recordings; wild-type model (state variables *S*(*t*) and *C_N_* (*t*)) fitted to wild-type and *NOS*-knockout Ca^2+^ INNOS recordings. Wild type model (state variables *S*(*t*), *C_N_* (*t*) and *N*(*t*)) fitted to wild type INNOS NO recordings. All visible correlation values are significant at the 5% level, *p* < 0.05, (trivial correlations along the diagonal are excluded). Figure 6 – figure supplement 5 – source data 1. Best fitted parameters `best_pars.mat`. Figure 6 – figure supplement 5 – source data 2. Script to generate the figure `correlations.m`

**Figure 6 – figure supplement 6.**
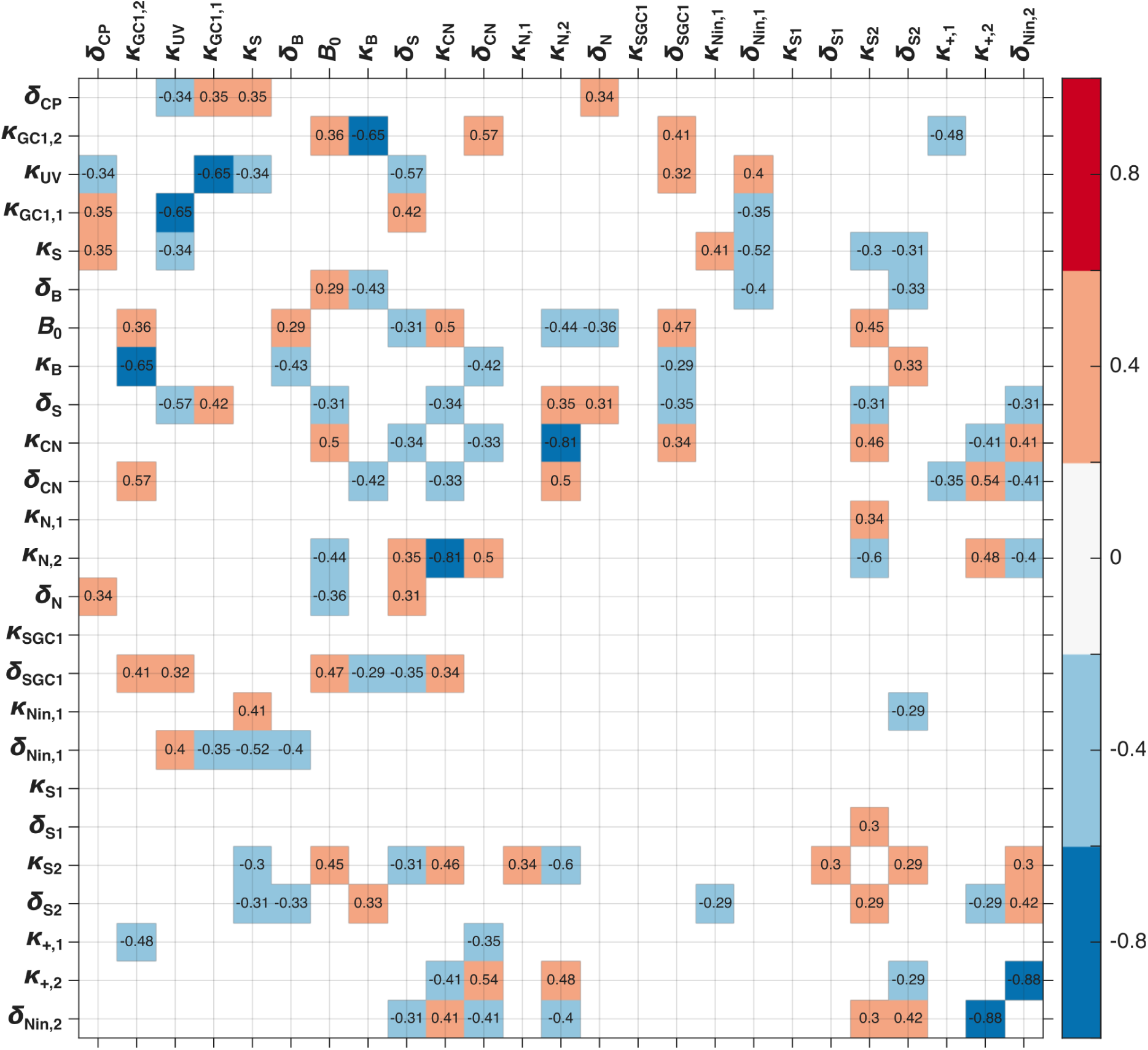
Pairwise correlations between parameters of the wild type model fitted to wild type cPRC recordings. All visible correlation values are significant at the 5% level, *p* < 0.05, (trivial correlations along the diagonal are excluded). Figure 6 – figure supplement 6 – source data 1. Best fitted parameters `best_pars.mat`. Figure 6 – figure supplement 6 – source data 2. Script to generate the figure `correlations.m`

**Figure 6 – figure supplement 7.**
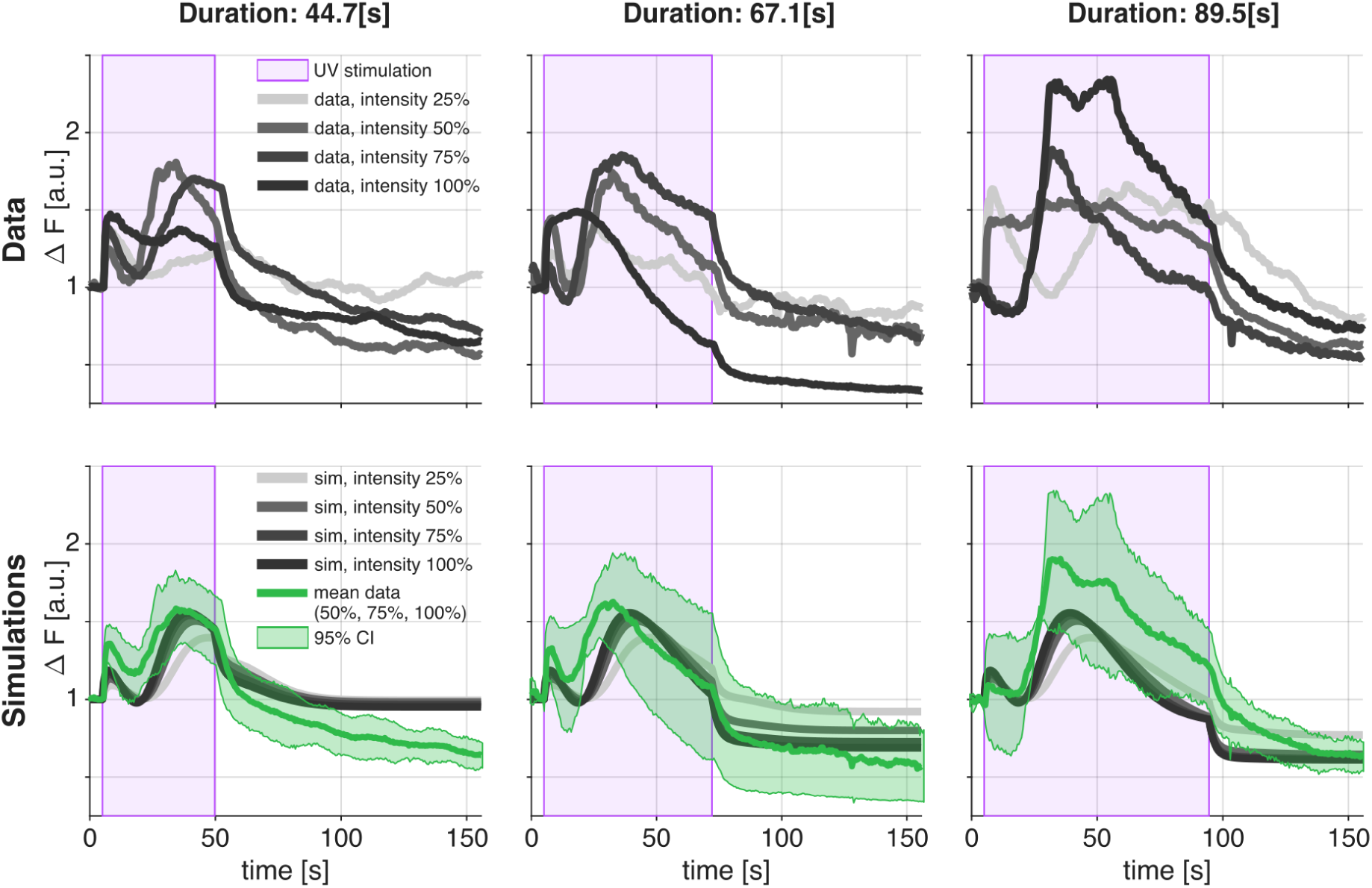
Validation of the wild-type model using cPRC recordings collected in wild-type larvae subject to UV stimulation with different intensity and duration. The top row displays recorded data; the bottom row shows model simulations. Different columns show different durations of UV stimulation. Simulated Ca^2+^ traces use parameter sets fitted to mean of Ca^2+^-recordings collected in wild-type larvae subject to 20s UV stimulation at 100% power. Stimulation duration is indicated by the shaded rectangle (violet) while stimulation intensity is indicated by the shade of grey of the line plot (darker shades indicate higher power). Simulations are compared with the mean of the 50%, 75% and 100% data and their 95% confidence interval (mean *pm*1.96*SE*, here is *SE* = *std*/*sqrtn* and *std* stands for standard deviation). We use the 50%, 75% and 100% recordings to compute the average because the data indicate the existence of a non-linear relation between response and the power of the stimulation and the 25% recordings would bias the average down. Figure 7 – source data 1. Recorded traces `longer_experiments_data.mat`. Figure 6 – figure supplement 7 – source data 1. Script to generate the figure `fig_longer_experiments.m`

**Figure 6 – figure supplement 8.**
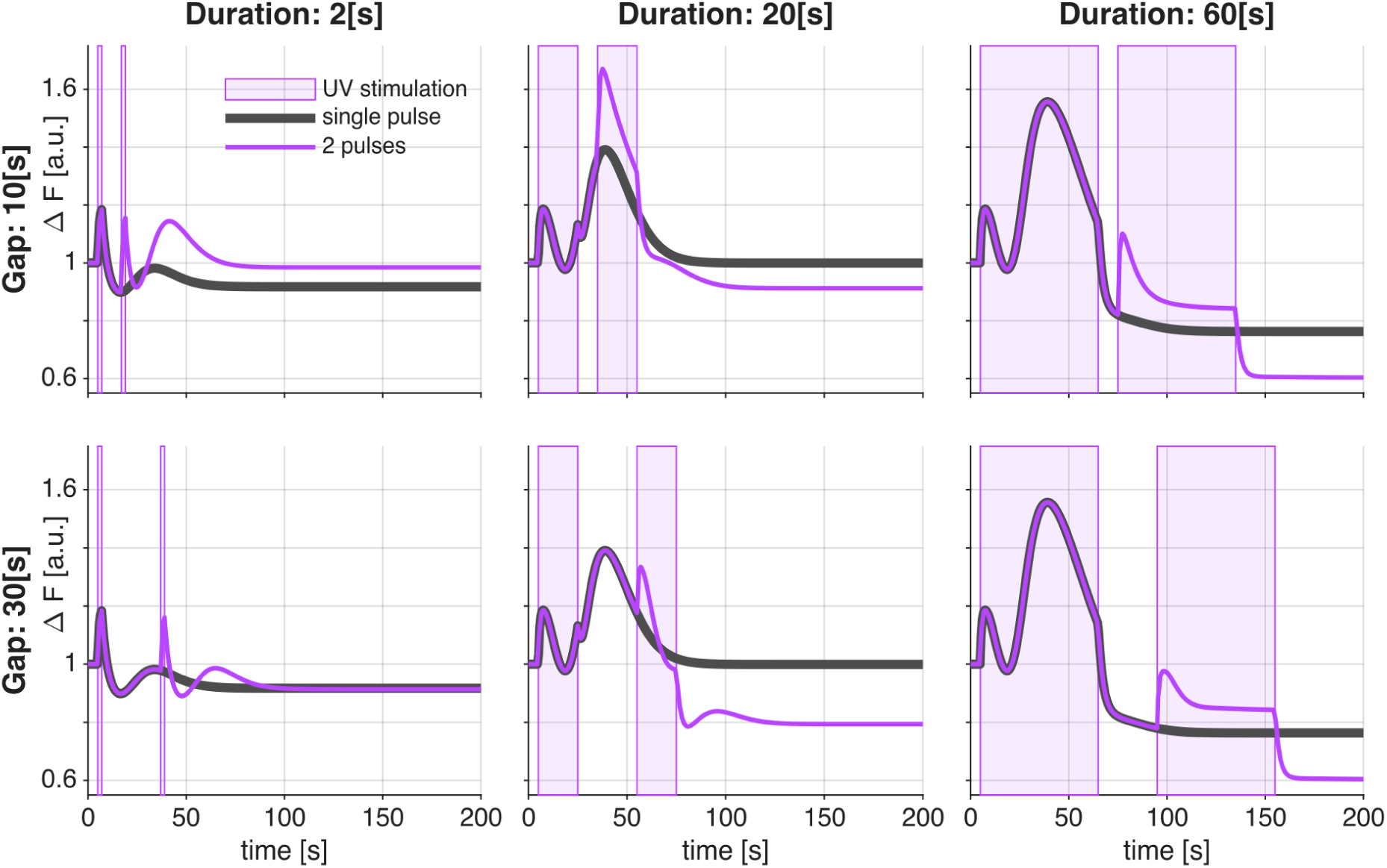
Simulated Ca^2+^ traces under stimulation with two UV pulses. The top row shows simulations with a 10s interval between pulses; the bottom row shows simulations with a 30s interval between pulses. Different columns show different durations of UV stimulation. The duration of UV stimulation is indicated by the shaded rectangle (violet). Coloured curves indicate simulated outcome of two pulse UV stimulation; thick black curves indicate simulated reference traces with a single UV pulse. Simulations of the Ca^2+^ traces use parameter sets fitted to the mean of individual Ca^2+^-recordings collected in wild-type larvae subject to 20s UV stimulation. Figure 6 – figure supplement 8 – source data 1. Script to generate the figure `fig_two_pulse.m`

## References

Achim K, Pettit J-B, Saraiva LR, Gavriouchkina D, Larsson T, Arendt D, Marioni JC. 2015. High- throughput spatial mapping of single-cell RNA-seq data to tissue of origin. Nature Biotechnology 33:503–509. doi:10.1038/nbt.3209

Arendt D, Tessmar-Raible K, Snyman H, Dorresteijn AW, Wittbrodt J. 2004. Ciliary photoreceptors with a vertebrate-type opsin in an invertebrate brain. Science 306:869–871. doi:10.1126/science.1099955

Aso Y, Ray RP, Long X, Bushey D, Cichewicz K, Ngo T-T, Sharp B, Christoforou C, Hu A, Lemire AL, Tillberg P, Hirsh J, Litwin-Kumar A, Rubin GM. 2019. Nitric oxide acts as a cotransmitter in a subset of dopaminergic neurons to diversify memory dynamics. eLife 8. doi:10.7554/elife.49257

Bargmann CI, Marder E. 2013. From the connectome to brain function. Nature Methods 10:483–490. doi:10.1038/nmeth.2451

Bauknecht P, Jékely G. 2017. Ancient coexistence of norepinephrine, tyramine, and octopamine signaling in bilaterians. BMC Biology 15. doi:10.1186/s12915-016-0341-7

Bezares-Calderón LA, Berger J, Jékely G. 2019. Diversity of cilia-based mechanosensory systems and their functions in marine animal behaviour. Philosophical Transactions of the Royal Society B: Biological Sciences 375:20190376. doi:10.1098/rstb.2019.0376

Bezares-Calderón LA, Shahidi R, Jékely G. 2023. A ciliary photoreceptor-cell circuit mediates pressure response in marine zooplankton.

Bishop CD, Brandhorst BP. 2007. Development of nitric oxide synthase-defined neurons in the sea urchin larval ciliary band and evidence for a chemosensory function during metamorphosis. Developmental Dynamics 236:1535–1546. doi:10.1002/dvdy.21161

Bray SR, Wyss LS, Chai C, Lozada ME, Wang B. 2023. Adaptive robustness through incoherent signaling mechanisms in a regenerative brain.

Bredt DS, Hwang PM, Snyder SH. 1990. Localization of nitric oxide synthase indicating a neural role for nitric oxide. Nature 347:768–770. doi:10.1038/347768a0

Burette A, Zabel U, Weinberg RJ, Schmidt HHHW, Valtschanoff JG. 2002. Synaptic localization of nitric oxide synthase and soluble guanylyl cyclase in the hippocampus. The Journal of Neuroscience 22:8961–8970. doi:10.1523/jneurosci.22-20-08961.2002

Camargo A, Llamas A, Schnell RA, Higuera JoseJ, Jì Gonzaĺez-Ballester D, Lefebvre PA, Fernańdez E, Galvań A. 2007. Nitrate signaling by the regulatory gene NIT2 in chlamydomonas. The Plant Cell 19:3491–3503. doi:10.1105/tpc.106.045922

Cannavó F. 2012. Sensitivity analysis for volcanic source modeling quality assessment and model selection. Computers & Geosciences 44:52–59. doi:10.1016/j.cageo.2012.03.008

Capella-Gutierrez S, Silla-Martinez JM, Gabaldon T. 2009. trimAl: A tool for automated alignment trimming in large-scale phylogenetic analyses. Bioinformatics 25:1972–1973. doi:10.1093/bioinformatics/btp348

Censor Y. 1977. Pareto optimality in multiobjective problems. Applied Mathematics and Optimization 4:41–59.

Choi HMT, Schwarzkopf M, Fornace ME, Acharya A, Artavanis G, Stegmaier J, Cunha A, Pierce NA. 2018. Third-generation in situ hybridization chain reaction: Multiplexed, quantitative, sensitive, versatile, robust. Development 145. doi:10.1242/dev.165753

Conzelmann M, Jékely G. 2012. Antibodies against conserved amidated neuropeptide epitopes enrich the comparative neurobiology toolbox. EvoDevo 3:23. doi:10.1186/2041-9139-3-23

Conzelmann M, Offenburger S-L, Asadulina A, Keller T, Münch TA, Jékely G. 2011. Neuropeptides regulate swimming depth of platynereis larvae. Proceedings of the National Academy of Sciences 108. doi:10.1073/pnas.1109085108

Conzelmann M, Williams Elizabeth A, Krug K, Franz-Wachtel M, Macek B, Jékely G. 2013a. The neuropeptide complement of the marine annelid platynereis dumerilii. BMC Genomics 14:906. doi:10.1186/1471-2164-14-906

Conzelmann M, Williams Elizabeth A, Tunaru S, Randel N, Shahidi R, Asadulina A, Berger J, Offermanns S, Jékely G. 2013b. Conserved MIP receptorligand pair regulates platynereis larval settlement. Proceedings of the National Academy of Sciences 110:8224–8229. doi:10.1073/pnas.1220285110

Cudeiro J, Rivadulla C. 1999. Sight and insight on the physiological role of nitric oxide in the visual system. Trends in Neurosciences 22:109–116. doi:10.1016/s0166-2236(98)01299-5

Deb K. 2001. Multi-objective optimization using evolutionary algorithms., Wiley-interscience series in systems and optimization. Wiley.

Díaz-Seoane S, Rey Barreiro X, Villaverde AF. 2022. STRIKE-GOLDD 4.0: User-friendly, efficient analysis of structural identifiability and observability. Bioinformatics 39. doi:10.1093/bioinformatics/btac748

Eddy SR. 2011. Accelerated profile HMM searches. PLOS Computational Biology 7:1–16. doi:10.1371/journal.pcbi.1002195

Frickey T, Lupas A. 2004. CLANS: A java application for visualizing protein families based on pairwise similarity. Bioinformatics 20:3702–3704. doi:10.1093/bioinformatics/bth444

Fu L, Niu B, Zhu Z, Wu S, Li W. 2012. CD-HIT: Accelerated for clustering the next-generation sequencing data. Bioinformatics 28:3150–3152. doi:10.1093/bioinformatics/bts565

Garthwaite J. 2015. From synaptically localized to volume transmission by nitric oxide. The Journal of Physiology 594:9–18. doi:10.1113/jp270297

Gibbs SM, Truman JW. 1998. Nitric oxide and cyclic GMP regulate retinal patterning in the optic lobe of drosophila. Neuron 20:83–93. doi:10.1016/s0896-6273(00)80436-5

Gühmann M, Jia H, Randel N, Verasztó C, Bezares-Calderón Luis A, Michiels Nico K, Yokoyama S, Jékely G. 2015. Spectral tuning of phototaxis by a go-opsin in the rhabdomeric eyes of platynereis. Current Biology 25:2265–2271. doi:10.1016/j.cub.2015.07.017

Guindon S, Dufayard J-F, Lefort V, Anisimova M, Hordijk W, Gascuel O. 2010. New algorithms and methods to estimate maximum-likelihood phylogenies: Assessing the performance of PhyML 3.0. Systematic Biology 59:307–321. doi:10.1093/sysbio/syq010

Hird C, Jékely G, Williams EA. 2024. Microalgal biofilm induces larval settlement in the model marine worm *platynereis dumerilii*. Royal Society Open Science 11. doi:10.1098/rsos.240274

Hoang DT, Chernomor O, Haeseler A von, Minh BQ, Vinh LS. 2018. UFBoot2: Improving the ultrafast bootstrap approximation. Molecular Biology and Evolution 35:518–522. doi:10.1093/molbev/msx281

Hölscher C. 1997. Nitric oxide, the enigmatic neuronal messenger: Its role in synaptic plasticity. Trends in Neurosciences 20:298–303. doi:10.1016/s0166-2236(97)01065-5

Imambocus BN, Zhou F, Formozov A, Wittich A, Tenedini FM, Hu C, Sauter K, Macarenhas Varela E, Herédia F, Casimiro AP, Macedo A, Schlegel P, Yang C-H, Miguel-Aliaga I, Wiegert JS, Pankratz MJ, Gontijo AM, Cardona A, Soba P. 2022. A neuropeptidergic circuit gates selective escape behavior of drosophila larvae. Current Biology 32:149–163.e8. doi:10.1016/j.cub.2021.10.069

Jacoby J, Nath A, Jessen ZF, Schwartz GW. 2018. A self-regulating gap junction network of amacrine cells controls nitric oxide release in the retina. Neuron 100:1149–1162.e5. doi:10.1016/j.neuron.2018.09.047

Jékely G, Colombelli J, Hausen H, Guy K, Stelzer E, Nédélec F, Arendt D. 2008. Mechanism of phototaxis in marine zooplankton. Nature 456:395–399. doi:10.1038/nature07590

Jékely G, Yuste R. 2024. Nonsynaptic encoding of behavior by neuropeptides. Current Opinion in Behavioral Sciences 60:101456. doi:10.1016/j.cobeha.2024.101456

Kane EA, Gershow M, Afonso B, Larderet I, Klein M, Carter AR, Bivort BL de, Sprecher SG, Samuel ADT. 2013. Sensorimotor structure of drosophila larva phototaxis. Proceedings of the National Academy of Sciences 110. doi:10.1073/pnas.1215295110

Katoh K, Toh H. 2008. Recent developments in the MAFFT multiple sequence alignment program. Briefings in Bioinformatics 9:286–298. doi:10.1093/bib/bbn013

Konak A, Coit DW, Smith AE. 2006. Multi-objective optimization using genetic algorithms: A tutorial. Reliability Engineering & System Safety 91:992–1007. 10.1016/j.ress.2005.11.018

Kuehn E, Clausen DS, Null RW, Metzger BM, Willis AD, Özpolat BD. 2021. Segment number threshold determines juvenile onset of germline cluster expansion in platynereis dumerilii. Journal of Experimental Zoology Part B: Molecular and Developmental Evolution 338:225–240. doi:10.1002/jez.b.23100

Kuntz S, Poeck B, Strauss R. 2017. Visual working memory requires permissive and instructive NO/cGMP signaling at presynapses in the drosophila central brain. Current Biology 27:613–623. doi:10.1016/j.cub.2016.12.056

Leise EM, Thavaradhara K, Durham NR, Turner BE. 2001. Serotonin and nitric oxide regulate metamorphosis in the marine snail ilyanassa obsoleta. American Zoologist 41:258–267. doi:10.1093/icb/41.2.258

Locascio A, Vassalli QA, Castellano I, Palumbo A. 2022. Novel insights on nitric oxide synthase and NO signaling in ascidian metamorphosis. International Journal of Molecular Sciences 23:3505. doi:10.3390/ijms23073505

Lundberg JO, Weitzberg E, Shiva S, Gladwin MT. 2011. The nitratenitritenitric oxide pathway in mammals. Humana Press. pp. 21–48. doi:10.1007/978-1-60761-616-0_3

Matsuda S, Harada K, Ito M, Takizawa M, Wongso D, Tsuboi T, Kitaguchi T. 2016. Generation of a cGMP indicator with an expanded dynamic range by optimization of amino acid linkers between a fluorescent protein and PDE5? ACS Sensors 2:46–51. doi:10.1021/acssensors.6b00582

Miller BL, Goldberg DE, others. 1995. Genetic algorithms, tournament selection, and the effects of noise. Complex systems 9:193–212.

Mobley RB, Ray EJ, Maruska KP. 2022. Expression and localization of neuronal nitric oxide synthase in the brain and sensory tissues of the african cichlid fish astatotilapia burtoni. Journal of Comparative Neurology 530:2901–2917. doi:10.1002/cne.25383

Möller MN, Rios N, Trujillo M, Radi R, Denicola A, Alvarez B. 2019. Detection and quantification of nitric oxidederived oxidants in biological systems. Journal of Biological Chemistry 294:14776– 14802. doi:10.1074/jbc.rev119.006136

Moroz LL, Meech RW, Sweedler JV, Mackie GO. 2004. Nitric oxide regulates swimming in the jellyfishAglantha digitale. The Journal of Comparative Neurology 471:26–36. doi:10.1002/cne.20023

Moroz LL, Romanova DY, Nikitin MA, Sohn D, Kohn AB, Neveu E, Varoqueaux F, Fasshauer D. 2020. The diversification and lineage-specific expansion of nitric oxide signaling in placozoa: Insights in the evolution of gaseous transmission. Scientific Reports 10. doi:10.1038/s41598-020-69851-w

Ozpolat BD, Randel N, Williams EA, Bezares-Calderón LA, Andreatta G, Balavoine G, Bertucci PY, Ferrier DEK, Gambi MC, Gazave E, Handberg-Thorsager M, Hardege J, Hird C, Hsieh Y-W, Hui J, Mutemi KN, Schneider SQ, Simakov O, Vergara HM, Vervoort M, Jékely G, Tessmar-Raible K, Raible F, Arendt D. 2021. The nereid on the rise: Platynereis as a model system. Zenodo. doi:10.5281/ZENODO.4907400

Randel N, Asadulina A, Bezares-Calderón LA, Verasztó C, Williams EA, Conzelmann M, Shahidi R, Jékely G. 2014. Neuronal connectome of a sensory-motor circuit for visual navigation. eLife 3. doi:10.7554/elife.02730

Santos RM, Lourenço CF, Pomerleau F, Huettl P, Gerhardt GA, Laranjinha J, Barbosa RM. 2011. Brain nitric oxide inactivation is governed by the vasculature. Antioxidants & Redox Signaling 14:1011–1021. doi:10.1089/ars.2010.3297

Shettigar N, Chakravarthy A, Umashankar S, Lakshmanan V, Palakodeti D, Gulyani A. 2021. Discovery of a body-wide photosensory array that matures in an adult-like animal and mediates eyebrain-independent movement and arousal. Proceedings of the National Academy of Sciences 118. doi:10.1073/pnas.2021426118

Shu CJ, Ulrich LE, Zhulin IB. 2003. The NIT domain: A predicted nitrate-responsive module in bacterial sensory receptors. Trends in Biochemical Sciences 28:121–124. doi:10.1016/s0968-0004(03)00032-x

Song H, Hewitt OH, Degnan SM. 2021. Arginine biosynthesis by a bacterial symbiont enables nitric oxide production and facilitates larval settlement in the marine-sponge host. Current Biology 31:433–437.e3. doi:10.1016/j.cub.2020.10.051

Tessmar-Raible K, Steinmetz PRH, Snyman H, Hassel M, Arendt D. 2005. Fluorescent two-color whole mount in situ hybridization in platynereis dumerilii (polychaeta, annelida), an emerging marine molecular model for evolution and development. BioTechniques 39:460–464. doi:10.2144/000112023

Thomas DD. 2015. Breathing new life into nitric oxide signaling: A brief overview of the interplay between oxygen and nitric oxide. Redox Biology 5:225–233. doi:10.1016/j.redox.2015.05.002

Tooker RE, Lipin MY, Leuranguer V, Rozsa E, Bramley JR, Harding JL, Reynolds MM, Vigh J. 2013. Nitric oxide mediates activity-dependent plasticity of retinal bipolar cell output via s- nitrosylation. The Journal of Neuroscience 33:19176–19193. doi:10.1523/jneurosci.2792-13.2013

Tosches MA, Bucher D, Vopalensky P, Arendt D. 2014. Melatonin signaling controls circadian swimming behavior in marine zooplankton. Cell 159:46–57. doi:10.1016/j.cell.2014.07.042

TrackMate 7: Integrating state-of-the-art segmentation algorithms into tracking pipelines. n.d.

Tsukamoto H, Chen I-S, Kubo Y, Furutani Y. 2017. A ciliary opsin in the brain of a marine annelid zooplankton is ultraviolet-sensitive, and the sensitivity is tuned by a single amino acid residue. Journal of Biological Chemistry 292:12971–12980. doi:10.1074/jbc.m117.793539

Tsukamoto H, Kubo Y. 2023b. A self-inactivating invertebrate opsin optically drives biased signaling toward gβγ-dependent ion channel modulation. Proceedings of the National Academy of Sciences 120. doi:10.1073/pnas.2301269120

Tsukamoto H, Kubo Y. 2023a. A self-inactivating invertebrate opsin optically drives biased signaling toward gβγ-dependent ion channel modulation. Proceedings of the National Academy of Sciences 120. doi:10.1073/pnas.2301269120

Ueda N, Richards GS, Degnan BM, Kranz A, Adamska M, Croll RP, Degnan SM. 2016. An ancient role for nitric oxide in regulating the animal pelagobenthic life cycle: Evidence from a marine sponge. Scientific Reports 6. doi:10.1038/srep37546

Veedin Rajan VB, Häfker NS, Arboleda E, Poehn B, Gossenreiter T, Gerrard E, Hofbauer M, Mühlestein C, Bileck A, Gerner C, Ribera d’Alcala M, Buia MC, Hartl M, Lucas RJ, Tessmar-Raible K. 2021. Seasonal variation in UVA light drives hormonal and behavioural changes in a marine annelid via a ciliary opsin. Nature Ecology & Evolution 5:204–218. doi:10.1038/s41559-020-01356-1

Verasztó C, Gühmann M, Jia H, Rajan VBV, Bezares-Calderón LA, Piñeiro-Lopez C, Randel N, Shahidi R, Michiels NK, Yokoyama S, Tessmar-Raible K, Jékely G. 2018. Ciliary and rhabdomeric photoreceptor-cell circuits form a spectral depth gauge in marine zooplankton. eLife 7. doi:10.7554/elife.36440

Verasztó C, Jasek S, Gühmann M, Bezares-Calderón LA, Williams EA, Shahidi R, Jékely G. 2025. Whole-body connectome of a segmented annelid larva. eLife 13. doi:10.7554/elife.97964.3

Verasztó C, Jasek S, Gühmann M, Shahidi R, Ueda N, Beard JD, Mendes S, Heinz K, Bezares- Calderón LA, Williams E, Jékely G. 2020. Whole-animal connectome and cell-type complement of the three-segmented platynereis dumerilii larva.

Verasztó C, Ueda N, Bezares-Calderón LA, Panzera A, Williams EA, Shahidi R, Jékely G. 2017. Ciliomotor circuitry underlying whole-body coordination of ciliary activity in the platynereis larva. eLife 6. doi:10.7554/elife.26000

Vergara HM, Pape C, Meechan KI, Zinchenko V, Genoud C, Wanner AA, Mutemi KN, Titze B, Templin RM, Bertucci PY, Simakov O, Dürichen W, Machado P, Savage EL, Schermelleh L, Schwab Y, Friedrich RW, Kreshuk A, Tischer C, Arendt D. 2021. Whole-body integration of gene expression and single-cell morphology. Cell 184:4819–4837.e22. doi:10.1016/j.cell.2021.07.017

Vielma AH, Agurto A, Valdés J, Palacios AG, Schmachtenberg O. 2014. Nitric oxide modulates the temporal properties of the glutamate response in type 4 OFF bipolar cells. PLoS ONE 9:e114330. doi:10.1371/journal.pone.0114330

Wang G-Y, List DA van der, Nemargut JP, Coombs JL, Chalupa LM. 2007. The sensitivity of light- evoked responses of retinal ganglion cells is decreased in nitric oxide synthase gene knockout mice. Journal of Vision 7:7. doi:10.1167/7.14.7

Wei T, Schubert T, Paquet-Durand F, Tanimoto N, Chang L, Koeppen K, Ott T, Griesbeck O, Seeliger MW, Euler T, Wissinger B. 2012. Light-driven calcium signals in mouse cone photoreceptors. Journal of Neuroscience 32:6981–6994. doi:10.1523/jneurosci.6432-11.2012

Wildemann B, Bicker G. 1999. Developmental expression of nitric oxide/cyclic GMP synthesizing cells in the nervous system ofDrosophila melanogaster. Journal of Neurobiology 38:1–15. doi:10.1002/(sici)1097-4695(199901)38:1<1::aid-neu1>3.0.co;2-l

Williams EA, Verasztó C, Jasek S, Conzelmann M, Shahidi R, Bauknecht P, Mirabeau O, Jékely G. 2017. Synaptic and peptidergic connectome of a neurosecretory center in the annelid brain. eLife 6. doi:10.7554/elife.26349

Zhang Y, He L-S, Zhang G, Xu Y, Lee O-O, Matsumura K, Qian P-Y. 2012. The regulatory role of the NO/cGMP signal transduction cascade during larval attachment and metamorphosis of the barnacle balanus (=amphibalanus) amphitrite. Journal of Experimental Biology. doi:10.1242/jeb.070235

